# Spurious model comparisons are widespread in biomedical artificial intelligence

**DOI:** 10.64898/2026.05.17.724301

**Authors:** Tianchu Zeng, Hetu Li, Shaoshi Zhang, Yan Quan Tan, Fang Tian, Csaba Orban, Lijun An, Wanyu Che, Jingwen Cheng, Joanna Su Xian Chong, Niousha Dehestani, Zijian Dong, Xin Li, Zhizhou Li, Mervyn Jun Rui Lim, Yi Lin, Qinrui Ling, Zijie Ling, Xi Zhi Low, Sina Mansour L., Eric Kwun Kei Ng, Thuan Tinh Nguyen, Leon Qi Rong Ooi, Shreya Pande, Xing Qian, Jingxuan Ruan, Ziwen Wang, Yapei Xie, Chen Zhang, Yichi Zhang, Kaustubh Patil, Linden Parkes, Elvisha Dhamala, Sidhant Chopra, Andrew Zalesky, Avram J. Holmes, Simon Eickhoff, Juan Helen Zhou, Olivier Renaud, Nico U.F. Dosenbach, Konrad Kording, Danilo Bzdok, Thomas E. Nichols, B.T. Thomas Yeo

## Abstract

Cross-validation is routinely used to compare performance in biomedical artificial intelligence. Standard tests ignore correlation across cross-validation folds, inflating false-positive rates. In a PRISMA-guided meta-analysis of 184 studies (impact factor ≥15) across 30 biomedical fields, 97% use invalid tests. Among studies with abstract-level claims supported by invalid tests, 59% rely on a spurious comparison: one that loses significance after correcting for this correlation. On average, regaining significance requires a 43% larger effect. Extending these findings to the broader literature, we estimate that spurious comparisons occur in nearly two in three studies and support abstract-level claims in nearly one in three studies. Simulations confirm false-positive rates paradoxically approach 100% when cross-validation is repeated to improve stability. We introduce SHARP, a redesign of cross-validation, which best balances false-positive control and power among 13 benchmarked tests. These results reveal widespread fragility in biomedical artificial intelligence and offer a practical route to valid comparisons.

## Introduction

Biomedical research increasingly relies on artificial intelligence to extract predictive information from complex, high-dimensional data (Rajpurkar et al., 2022; Moor et al., 2023; Bzdok et al., 2024; Perez-Lopez et al., 2024). As algorithms and data modalities proliferate, comparisons of predictive performance guide which algorithms to deploy and biomarkers to prioritize (Greene et al., 2018; Wagner et al., 2023; Carrasco-Zanini et al., 2024; Yoo et al., 2025). Consider a hypothetical study in which the same algorithm is trained to predict dementia progression from blood or MRI biomarkers (Fig. 1a). Because the algorithm is held constant, superior performance of the MRI model would indicate that MRI carries more information about dementia progression than blood. Comparing model performance can therefore inform scientific and clinical applications, not merely benchmark algorithms.

**Figure 1.**
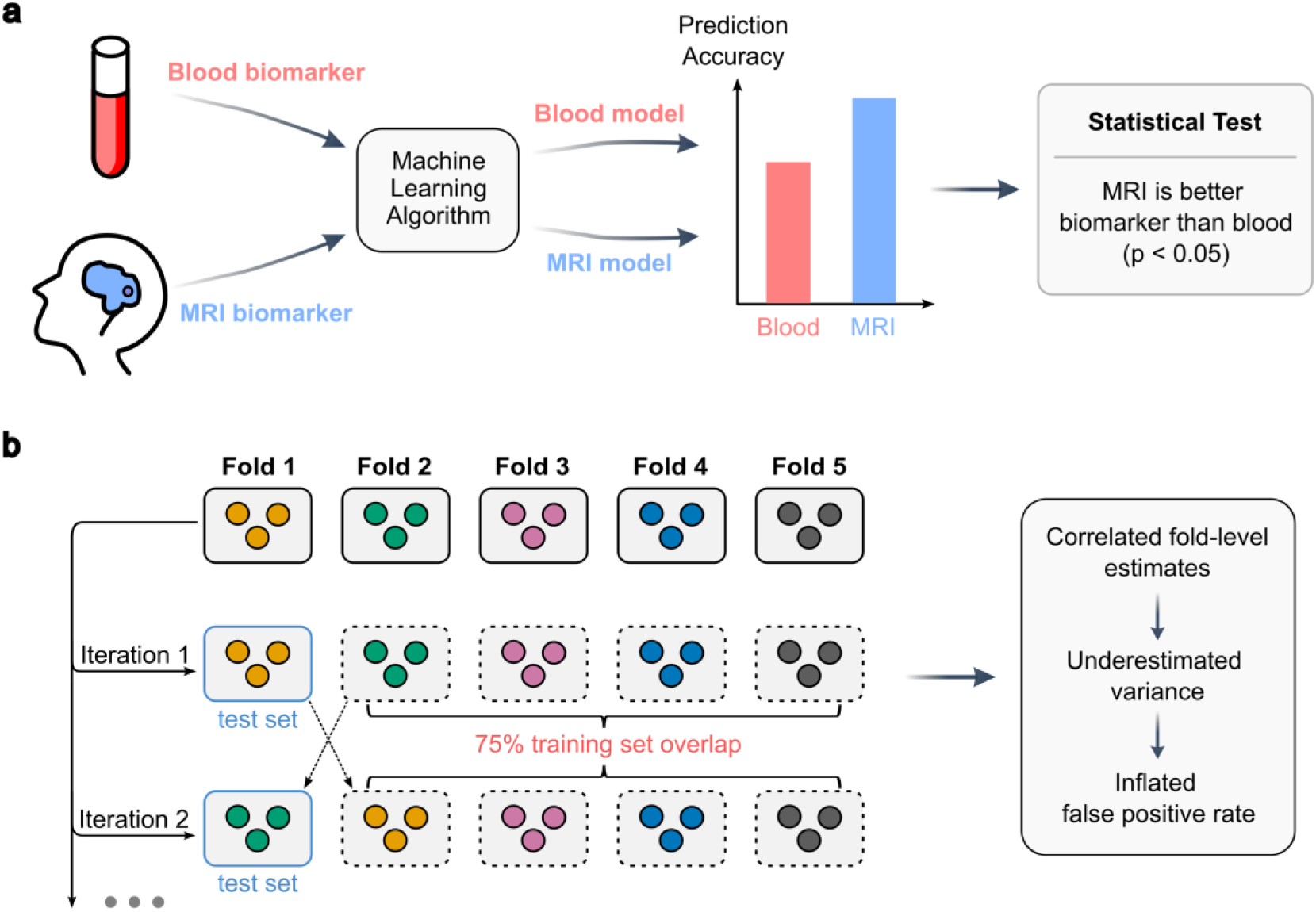
Comparing biomarkers using artificial intelligence and statistical dependence (correlation) induced by cross-validation. **a.** Comparing the predictive performance of artificial intelligence models can inform scientific and clinical applications. In this hypothetical example, the same algorithm is trained to predict dementia progression from blood or MRI biomarkers. A statistical test compares their prediction performance. Since the algorithm is held constant, superior performance of the MRI model would suggest that MRI is more informative than blood. **b.** Five-fold cross-validation partitions the dataset into five folds. In each iteration, one fold serves as the test set (blue outline) and four folds form the training set (dashed outline). Training sets overlap substantially across iterations, and each test fold contributes to the training set in all other iterations. This induces positive correlations among fold-level performance differences. Treating the fold-level differences as independent underestimates the variance of the average performance difference, thus inflating false positive rates (Nadeau & Bengio, 2003). Other common cross-validation variants are described in Methods; the Discussion considers the complementary role of external validation.

Predictive performance is commonly compared with K-fold cross-validation. The dataset is partitioned into *K* non-overlapping subsets (“folds”) of observations, such as brain scans or blood samples. In each iteration, one fold serves as the test set while the remaining *K* – 1 folds form the training set (Fig. 1b). When applied to two competing biomarkers (or algorithms), this procedure yields *K* pairs of prediction performance estimates that can be statistically compared. Although test folds do not overlap across iterations, training sets overlap substantially, and each test fold is reused for training in other iterations (Fig. 1b). The performance estimates are therefore correlated across folds (Dietterich, 1998; Bates et al., 2024). Conventional tests (e.g., paired t-tests) assume the correlated performance estimates are independent, thus underestimating uncertainty in the model comparison and inflating false positive rates (Fig. 1b; Nadeau & Bengio, 2003; Jafrasteh et al., 2025). Repeated cross-validation, often recommended to improve stability (Bouckaert & Frank, 2004; Varoquaux et al., 2017), reuses data more extensively and worsens the problem (Nadeau & Bengio, 2003).

Seminal studies dating back more than two decades have identified this statistical problem and introduced tests that account for correlation across folds (Dietterich, 1998; Nadeau & Bengio, 2003). Yet the prevalence and consequences of incorrectly assuming fold independence in biomedical research remain unclear. We ourselves have inadvertently made this error (Yeo et al., 2010; Mansour L. et al., 2021; Chopra et al., 2024), raising two broader questions. First, how often do biomedical studies ignore correlation induced by cross-validation, and does this vary across scientific fields? If publishing safeguards – editorial selectivity, statistical review, methodological reporting standards, open data and code – are effective, prevalence should be lower where safeguards are stronger. Second, and more consequentially, what proportion of published findings will remain significant after accounting for between-fold correlation? An invalid test can still lead to the same decision as a valid test, so the effect on published findings must be measured directly.

Accounting for correlation across folds is not as simple as applying a standard correction. When fold-level estimates are highly similar, this could reflect genuinely low fold-level variance or high between-fold correlation or a combination of the two. Standard cross-validation cannot distinguish among these possibilities, creating a fundamental statistical ambiguity (Nadeau & Bengio, 2003; Bengio & Grandvalet, 2004). Existing correlation-aware tests therefore make different compromises: some sacrifice power for more reliable control of the false positive rate (FPR; Dietterich, 1998; Parkes et al., 2021), whereas others preserve power by pre-specifying the between-fold correlation (Nadeau & Bengio, 2003) but risk inconsistent FPR control. This motivates redesigning cross-validation to estimate both variance and correlation directly.

To address these issues, we conduct a PRISMA-guided meta-analysis of 184 studies spanning 30 biomedical fields in journals with impact factor at least 15. For each study, we review the reporting and all available data and code. Where sufficient information is provided, we re-analyze originally significant model comparisons evaluated using invalid tests to determine whether they retain significance after accounting for between-fold correlation. To identify which analysis choices further inflate FPR when between-fold correlation is ignored, we perform simulations across 420 scenarios spanning image recognition, neuroimaging, ecology, and systems biology. We then introduce SHARP (Split-Half Analysis of Repeated Performance), a redesign of cross-validation that estimates variance and correlation directly, and benchmark it against 12 tests with respect to FPR control, power, and confidence interval coverage. Finally, we provide reporting recommendations and practical guidance on combining cross-validation with external validation for rigorous model development and evaluation.

## Results

### Invalid statistical tests in 97% of studies

Following the Preferred Reporting Items for Systematic Reviews and Meta-Analyses (PRISMA) guidelines (Page et al., 2021), we searched PubMed for original research articles published from 1 June 2020 to 1 June 2025 that used cross-validation to compare predictive performance. We restricted inclusion to journals with impact factor ≥ 15, thereby providing a conservative assessment of prevalence because these venues typically have stronger reporting standards, statistical review, and transparency policies.

The search yielded 2,400 studies, of which 1,875 compared model performance (Fig. 2a). Of these, 1,008 studies (54%) did not clearly apply a statistical test or report confidence intervals. Further exclusions for unclear procedures, absent fold-level statistics, or confidence-interval-only reporting yielded a final set of 184 studies (Fig. 2a; see Methods). The substantial winnowing reflects broader deficiencies in statistical reporting and motivates the minimum reporting standards outlined in the Discussion.

**Figure 2.**
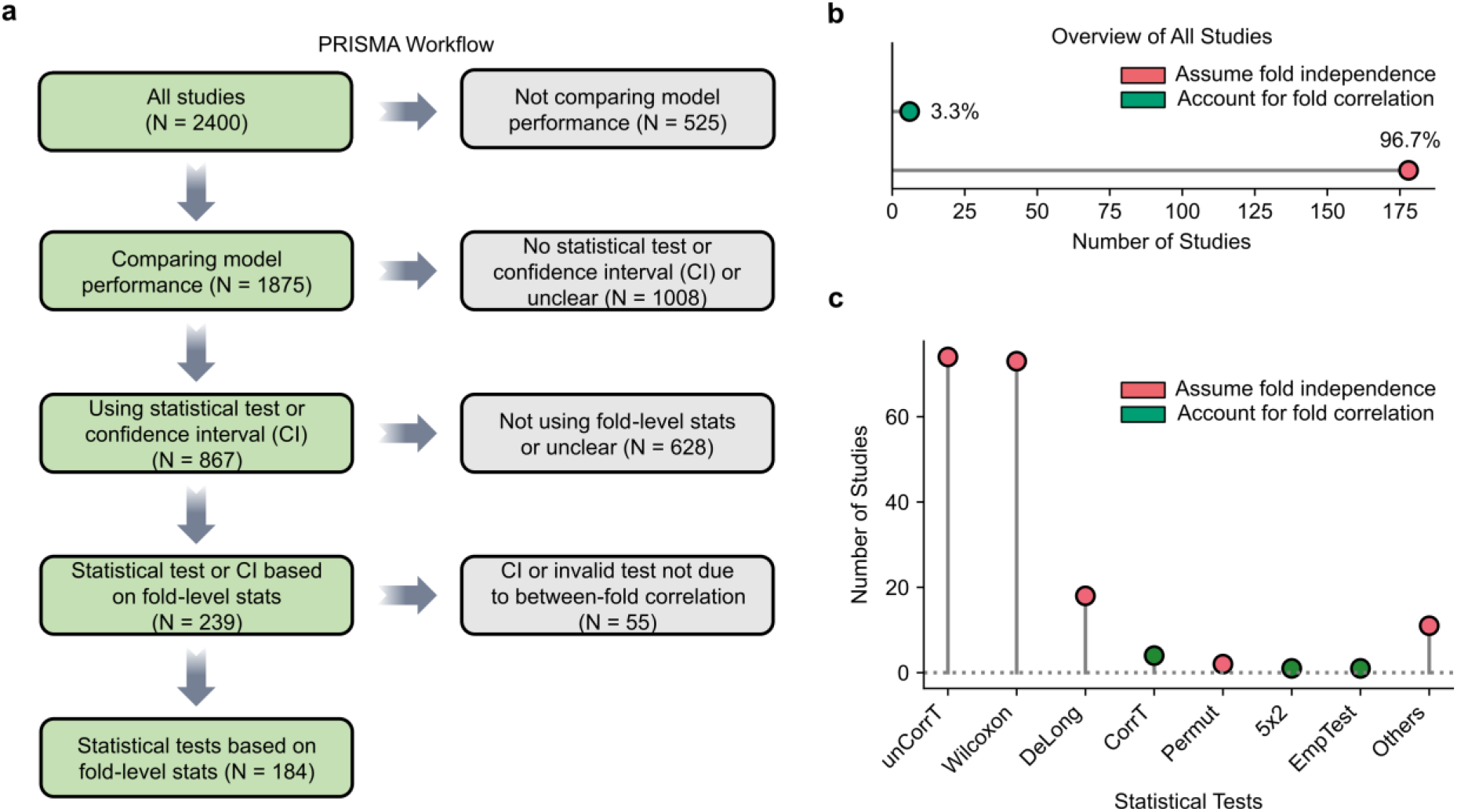
Widespread use of invalid statistical tests in biomedical studies using cross-validation to compare predictive performance. **a.** PRISMA flow diagram. A PubMed search (1 June 2020–1 June 2025; journals with impact factor ≥15) identified 2,400 records, of which 1,875 compared model performance. Of these, 867 clearly applied a statistical test or reported confidence intervals. We excluded 628 studies without fold-level statistics or with unclear procedures. We further excluded one study using a permutation test invalid for other reasons (see Supplementary Results) and 54 studies reporting only confidence intervals because the method of computation was rarely clear (see Methods). This resulted in a final set of 184 studies. **b.** Breakdown of the 184 studies by statistical approach: tests assuming fold independence (red) versus tests accounting for between-fold correlation (green). **c.** Breakdown of specific statistical tests across the 184 studies. Red indicates tests that assume fold independence, i.e., invalid tests. Green indicates tests that account for between-fold correlation. “unCorrT” and “CorrT” refer to the uncorrected and corrected resampled t-tests, respectively (Nadeau & Bengio, 2003); “Wilcoxon” refers to either the Wilcoxon signed-rank test (Wilcoxon, 1945) or the Wilcoxon rank-sum test (Mann & Whitney, 1947); “DeLong” refers to the DeLong test (DeLong et al., 1988); “Permut” refers to a paired permutation test (Edgington & Onghena, 2007); “5×2” refers to the “5×2 paired t-test” (Dietterich, 1998); and “EmpTest” refers to a variant of the empirical test of differences (Parkes et al., 2021). Further breakdown of the invalid tests grouped under “Others” is provided in Supplementary Results. Supplementary File 1 provides summary information for each study, while Supplementary File 2 provides detailed results for each study.

Of the 184 studies (Fig. 2b), 178 (97%) used invalid statistical tests that incorrectly assumed fold independence. Invalid testing was similarly prevalent among studies using classical machine learning, deep neural networks, or both: 95%, 97%, and 100%, respectively. The naïve (uncorrected) t-test and Wilcoxon tests were the most common invalid tests (Fig. 2c). As we show later, many comparisons deemed significant by these invalid tests lose significance under valid re-analysis.

### Invalid testing is prevalent regardless of year, scientific field or publishing safeguards

We next tested whether invalid-test prevalence varied by publication year, scientific field or publishing safeguards. Scientific fields were defined using Web of Science Journal Citation Reports subject categories, and journals were assigned a rigor score from 0 to 14 spanning transparency, reproducibility, and statistical rigor.

The number of studies grew by 14.5% per year from 1 June 2020 to 1 June 2025 (Poisson regression *p* = 0.009; Fig. 3a). Over the same period, the proportion of studies assuming fold independence (i.e., ignoring between-fold correlation) showed no detectable change and remained high throughout (permutation test *p* = 0.63; Fig. 3b).

**Figure 3.**
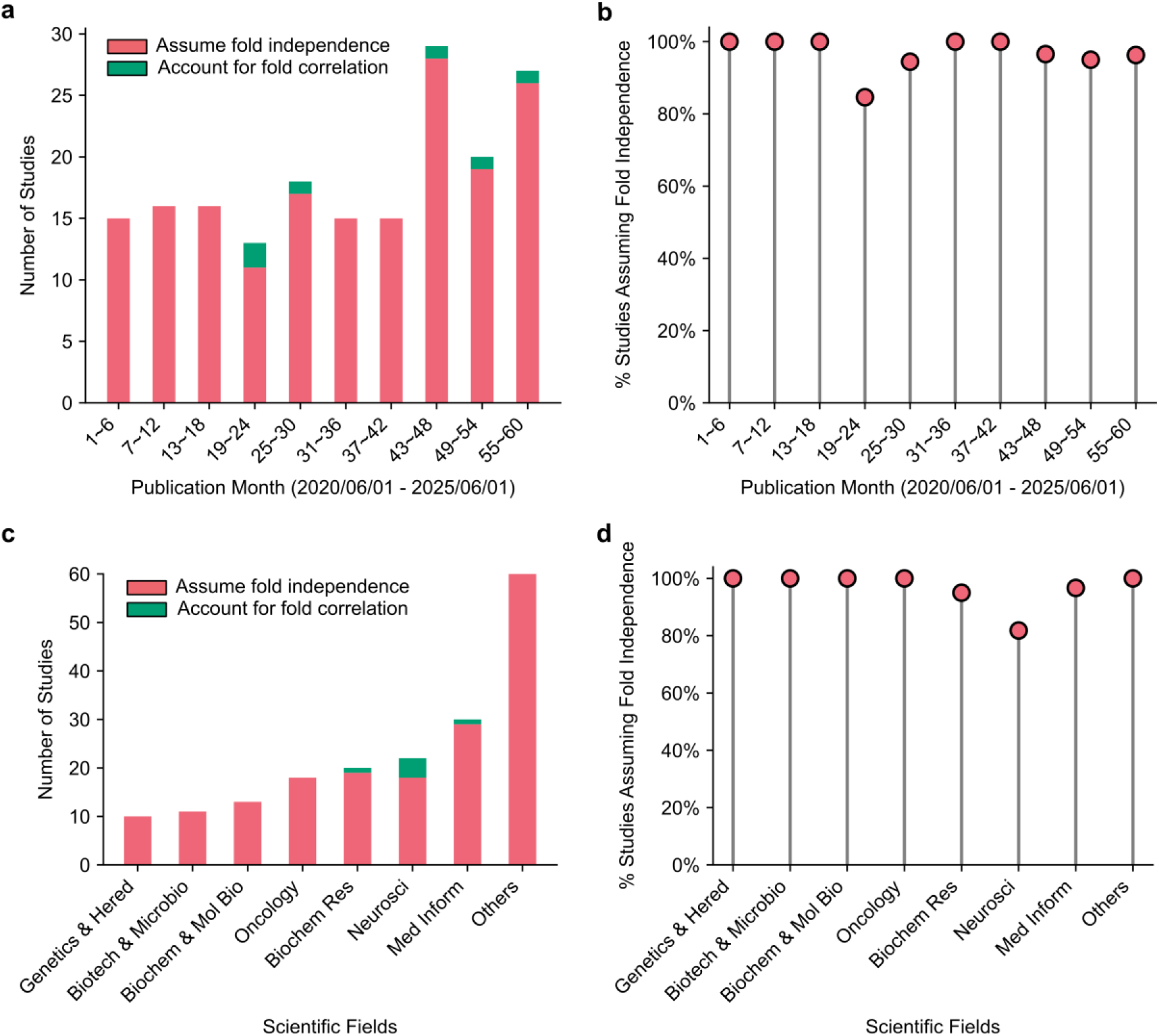
Invalid testing persists over time and across scientific fields. **a.** Number of studies in 6-month intervals from 1 June 2020 to 1 June 2025 (N = 184). The number increased by 14.5% per year (Poisson regression, *p* = 0.009). **b.** Proportion of studies assuming fold independence (i.e., ignoring between-fold correlation) over time. No temporal change was detected (permutation test, *p* = 0.63). **c.** Number of studies across 30 scientific fields. Seven fields with at least 10 studies are shown separately. The remaining 23 fields are grouped as “Others”. **d.** Proportion of studies assuming fold independence by scientific field. No difference was detected across fields (permutation test, *p* = 0.07); “Others” was excluded from this comparison. Statistical modeling is detailed in Methods. Supplementary File 1 provides summary information for each study, while Supplementary File 2 provides detailed results for each study.

Across scientific fields, reliance on invalid tests was ubiquitous (Figs. 3c,d). Among the seven fields with at least 10 studies, the proportion ignoring between-fold correlation ranged from 82% in “Neuroscience” to 100% in “Genetics & Heredity”, “Biotechnology & Applied Microbiology”, and “Biochemistry & Molecular Biology” (Fig. 3d). No difference was detected across fields (permutation test *p* = 0.07).

There was no association between ignoring between-fold correlation and journal impact factor (permutation test *p* = 0.58; Supplementary Fig. 1a,b), journal rigor score (*p* = 0.89; Supplementary Fig. 1c,d), or study-level data and code availability (*p* = 1.00 for both; Supplementary Fig. 1e,f). Prevalence remained near-universal even in journals with stringent policies and among studies sharing data, code, or both.

The persistence of invalid tests across years, fields, and publishing safeguards indicates a systemic failure to account for correlation across folds.

### Most studies contain a spurious comparison

A test with an elevated FPR can still lead to the same decision as a valid test, so its effect on published findings must be measured directly. Of the 184 studies, six used valid tests and five ignored between-fold correlation but reported no significant comparisons, leaving 173 at risk. For 105 studies, the original analyses or the reporting, data, and code provided did not permit valid re-analysis without additional assumptions. The remaining 68 contributed at least one comparison that could be re-analyzed with a valid test (see Methods).

For comparisons within a single dataset, we used the corrected resampled t-test to account for between-fold correlation (Nadeau & Bengio, 2003). Some comparisons spanned independent datasets. We averaged performance differences across folds within each dataset and compared models across datasets (see Methods). SHARP cannot be applied retrospectively because it requires a modified cross-validation scheme.

Throughout this study, we call an originally significant comparison spurious if it does not survive valid re-analysis (corrected *p* ≥ 0.05). Across all 68 studies, 47 (69%) contained a spurious comparison (Fig. 4a). Of the 68 studies, 37 (54%) made a claim in the abstract supported by an invalid test. In 22 of these 37 studies (59%), a supporting comparison was spurious (Fig. 4b).

**Figure 4.**
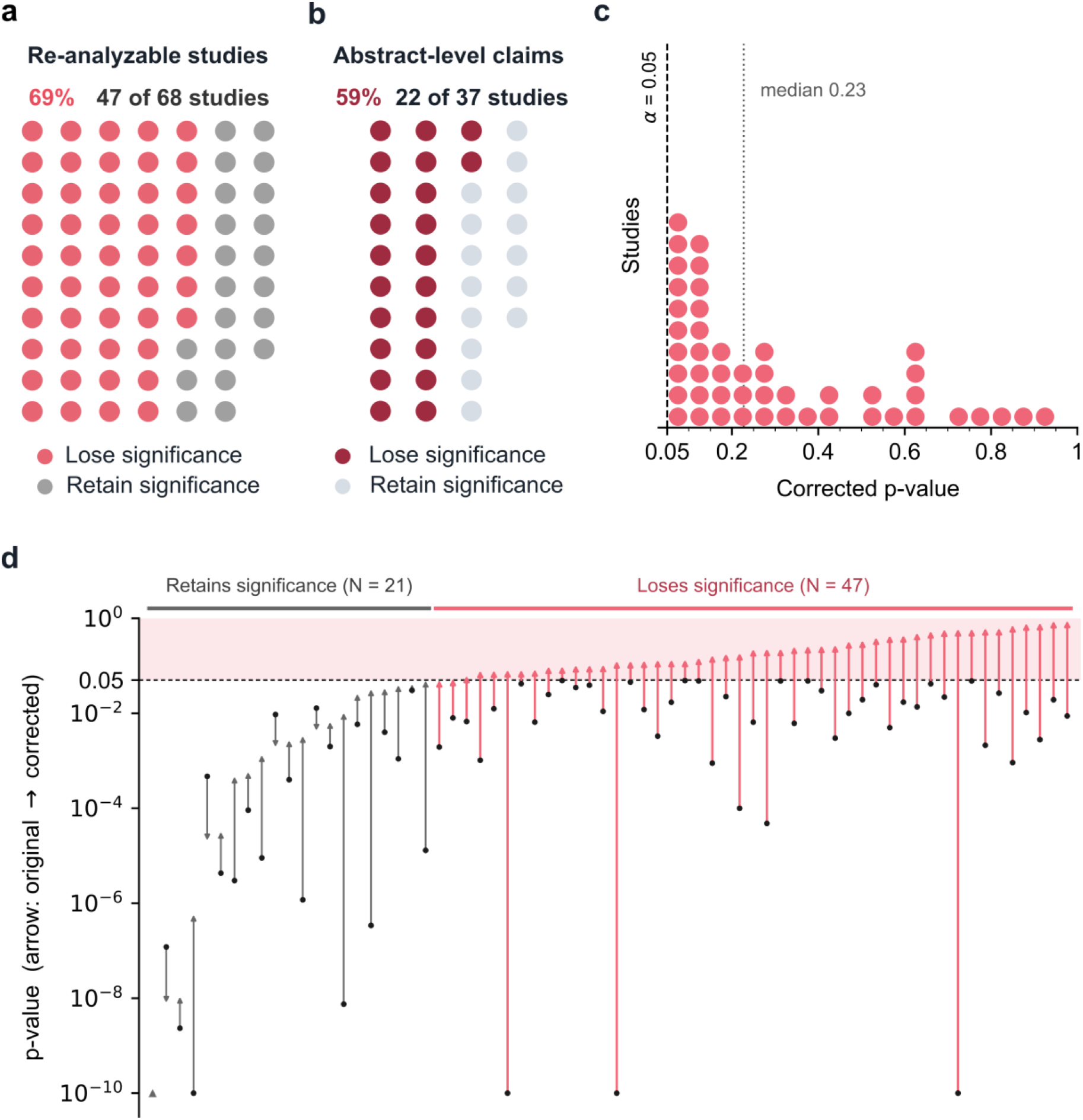
Most studies contain a spurious comparison. **a.** Study-level outcomes across 68 studies contributing at least one re-analyzable comparison. Each circle represents one study. Of these, 47 (69%; red) contained a comparison that proved to be spurious: it became non-significant under valid re-analysis. Comparisons retained significance in the remaining 21 studies (31%; grey). **b.** Study-level outcomes for the 37 studies with a claim in the abstract supported by an invalid test. In 22 studies (59%; dark red), a supporting comparison was spurious. Supporting comparisons retained significance in the remaining 15 studies (41%; grey). **c.** Largest corrected p-value in each of the 47 studies containing a spurious comparison. In the median study, the largest corrected p-value is 0.23 (dashed line). **d.** Original and corrected p-values for the comparison with the largest corrected p-value in each of the 68 studies, sorted by corrected p-value. Each arrow runs from the original p-value (black circle) to the corrected p-value. When only an upper bound was reported (e.g., ‘*p* < 0.001’), the black circle instead shows the reconstructed uncorrected p-value. The shaded region marks *p* ≥ 0.05. Triangles denote arrows extending below the axis limit. Summary information for each study is provided in Supplementary File 1. Detailed results for each study are found in Supplementary File 2.

The spurious comparisons were often not borderline. Across the 47 studies containing a spurious comparison, the median of each study’s largest corrected p-value was 0.23 (Fig. 4c). Taking the median corrected p-value among spurious comparisons within each of the 47 studies and then the median across studies yielded 0.15. For the 22 studies with spurious comparisons supporting abstract-level claims, the same calculation yielded 0.17.

Using this abstract-level summary, *p* = 0.17 corresponds to an absolute z-statistic of 1.37 under a two-sided standard-normal reference, compared with 1.96 at *p* = 0.05. On average, regaining significance would require a 43% larger effect size. As an illustration, a conventional analysis of independent observations would require approximately twice as many observations to close this gap, assuming the same population-level effect and variance. This illustration is not a sample size calculation for cross-validation studies (Supplementary Results). In many studies, the largest corrected p-value exceeded the original p-value by more than an order of magnitude (Fig. 4d).

Several features suggest that the 69% estimate is conservative. First, some comparisons in the 68 studies could not be re-analyzed, including those evaluated using tests even more vulnerable to FPR inflation, such as the DeLong test (Supplementary Fig. 2). Second, as we show later, the corrected resampled *t*-test does not always control the FPR. Third, we did not include multiple-comparison corrections unless the original study performed a multiple-comparison correction and we could reproduce the resulting adjusted p-values. Broader correction could only cause further significance losses.

Eligibility for re-analysis required sufficiently complete reporting, data, or code, giving no reason to regard the remaining studies as methodologically safer. We therefore extrapolated the 69% rate to all 173 at-risk studies, estimating that nearly two in three studies in our meta-analysis – (173/184) × (47/68) = 65% – contain a spurious comparison. Because the use of tests ignoring between-fold correlation was not associated with journal impact factor or journal rigor score, we expect a comparable proportion in the broader literature.

To calculate a similar prevalence for abstract-level claims, we note that 100 of the 184 studies had an abstract-level claim relying on an invalid test. Therefore, we estimate that (100/184) × (22/37) = 32% (or almost one-third) of studies in the broader literature contain an abstract-level claim supported by a spurious comparison.

### Tests assuming fold independence poorly control the false positive rate (FPR)

Published studies often contain a spurious comparison, but we cannot estimate the FPR because it is unknown whether the null is true for each individual study. We therefore turned to simulations in which the null hypothesis holds by construction, using four datasets spanning image recognition (EMNIST handwritten digits; Cohen et al., 2017), neuroimaging (UK Biobank; Alfaro-Almagro et al., 2018), ecological classification (Covertype; Blackard, 1998), and systems biology (KEGG Metabolic Pathway; Naeem & Asghar, 2011). Further details are provided in Methods.

To estimate FPR, we constructed pairs of trained models with identical expected predictive performance and compared them using a statistical test (Fig. 5a). First, we randomly sampled a dataset of a given sample size (e.g., *N* = 1,000) from one of the four datasets. For each sampled dataset, we generated two noisy copies by independently permuting the same fraction of target labels, then applied the same machine-learning algorithm and cross-validation scheme to both copies. The independent noise ensured that the two trained models were distinct, while the matched noise level ensured equal expected performance under the null. At a significance threshold of α = 0.05, a well-calibrated test should reject the null hypothesis no more than 5% of the time (Fig. 5a).

**Figure 5.**
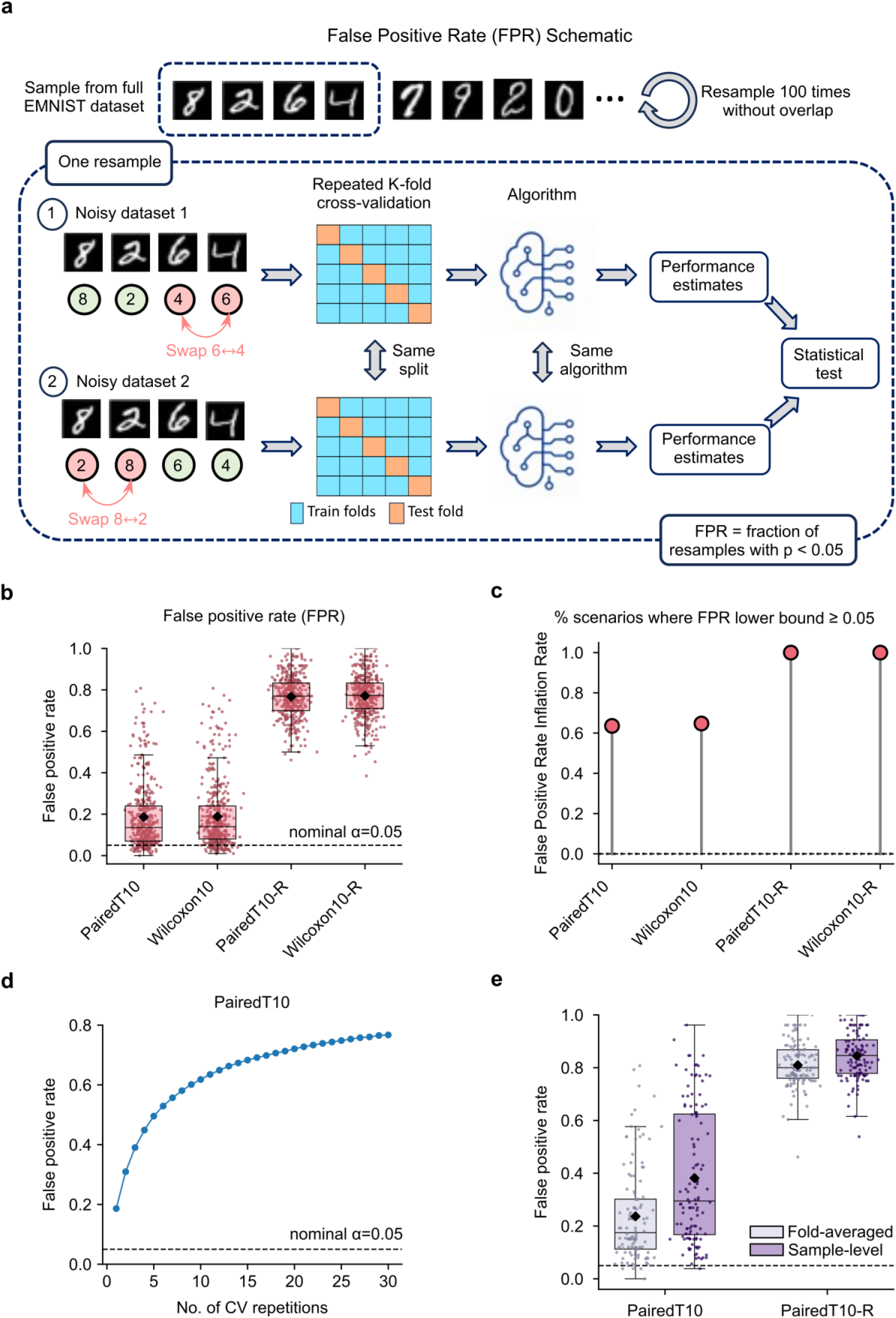
Tests that assume fold independence poorly control false positive rates. **a.** False positive rate (FPR) simulation illustrated for the EMNIST handwritten digit dataset. We randomly sampled a dataset (e.g., *N* = 1,000) from a full dataset, then generated two noisy copies by independently permuting the target labels for the same proportion of samples (e.g., 10%). For example, noisy dataset 1 (top) has the labels for digits 6 and 4 swapped, while noisy dataset 2 (bottom) has the labels for digits 8 and 2 swapped. Applying the same algorithm and identical cross-validation splits to both copies yielded paired performance estimates that were compared using a statistical test. The independent permutations produced distinct trained models, whereas the matched noise level ensured equal expected performance. At α = 0.05, a well-calibrated test should reject no more than 5% of the time. This procedure was repeated up to 100 times using non-overlapping subsamples. **b.** FPR across four datasets and four sample sizes. Each boxplot contains 420 data points, one per scenario. Black dots indicate the mean FPR. The dashed line indicates the nominal FPR of 0.05. PairedT10 and Wilcoxon10 denote the paired t-test and Wilcoxon signed-rank test under 10-fold cross-validation; the “-R” suffix denotes the same tests with 30 repetitions of 10-fold cross-validation. **c.** FPR inflation rate: percentage of the 420 scenarios in which the 95% confidence interval for FPR lay entirely above 0.05. **d.** FPR of the paired t-test as a function of the number of 10-fold cross-validation repetitions, averaged over 420 scenarios. The dashed line indicates the nominal FPR of 0.05. The Wilcoxon signed-rank test yields the same conclusions (not shown). **e.** FPR of the paired t-test using sample-level versus fold-averaged statistics, across 120 scenarios in the KEGG Metabolic Pathway dataset (one data point per scenario). The black dot indicates the mean FPR. The dashed line indicates the nominal FPR of 0.05.

We evaluated 140 combinations of dataset, sample size, and noise level under three hyperparameter-selection strategies – fixed hyperparameters, a single training-validation split, and nested cross-validation – yielding 420 scenarios. Repetition on non-overlapping subsamples, typically 100 per scenario, provided independent p-values from which we estimated FPR and its 95% confidence interval.

Across the 420 scenarios, the two most widely used tests (the paired t-test and Wilcoxon signed-rank test; Fig. 2c) exhibited an elevated FPR of ∼19% when 10-fold cross-validation was performed once (Fig. 5b). In ∼64% of scenarios, the 95% CI for the estimated FPR lay entirely above 0.05; we refer to this percentage as the “FPR inflation rate” (Fig. 5c). When 10-fold cross-validation was repeated 30 times – as is often recommended for stability (Bouckaert & Frank, 2004; Varoquaux et al., 2017) – the FPR rose to 77% (Fig. 5b) and FPR inflation was detected in 100% of scenarios. FPR rose monotonically with the number of repetitions (Fig. 5d), eventually reaching 100%.

The problem lies not with repeated cross-validation itself, which has well-established benefits, but with its interaction with tests that assume fold independence. Even when two models have the same performance at the population-level (i.e., null hypothesis is true), a finite dataset will always exhibit some non-zero difference between the two models. Tests that ignore between-fold correlation treat each fold-level estimate as independent. Consequently, the apparent sample size grows with each cross-validation repetition, so the estimated uncertainty artificially shrinks until the observed difference is declared significant, yielding 100% FPR.

### Sample-level statistics compound false positive rate inflation

So far we have focused on “fold-averaged” statistics, but our meta-analysis also identified studies using “sample-level” statistics. For example, 10-fold cross-validation on 1,000 samples yields 10 paired performance values after averaging within folds (i.e., fold-averaged statistics), whereas a sample-level analysis enters all 1,000 paired performance values directly into the statistical test. Because performance values within a fold share the same trained model, sample-level analysis introduces additional within-fold correlation.

In the KEGG Metabolic Pathway dataset, switching from fold-averaged to sample-level statistics increased FPR from 24% to 38% for one run of 10-fold cross-validation and from 81% to 84% for 30 repetitions (Fig. 5e). The DeLong test, another sample-level test, showed similarly inflated FPRs in the Covertype classification dataset (Supplementary Fig. 2). Together, these results show that sample-level statistics compound the FPR inflation caused by assuming fold independence.

### SHARP redesigns cross-validation to sidestep a fundamental statistical ambiguity

Having shown the practical consequences of ignoring between-fold correlation, we next considered correlation-aware tests. These tests face a fundamental statistical ambiguity. In a single run of K-fold cross-validation, the distribution of the *K* fold-level differences is governed by three unknowns: true mean μ, true fold-level variance σ², and true between-fold correlation ρ. From the *K* differences, we can compute the sample mean and sample variance. The sample mean is an unbiased estimate of μ, but the expected sample variance equals σ²(1 − ρ). A low sample variance could therefore reflect low fold-level variance σ² or high between-fold correlation ρ, or a combination of the two. Thus, σ² and ρ cannot be disentangled, yet both are needed to quantify uncertainty in the sample mean for valid statistical testing (Nadeau & Bengio, 2003; Supplementary Methods S6). This non-identifiability persists under repeated cross-validation, and no universal distribution-independent unbiased estimator of cross-validation variance exists (Bengio & Grandvalet, 2004).

Existing correlation-aware tests address this limitation through different compromises. Of the six studies in our meta-analysis that used such tests (Fig. 2c), four used the corrected resampled *t*-test (Nadeau & Bengio, 2003), one used the 5×2 paired *t*-test (Dietterich, 1998), and one used a variant of the empirical test of differences (Parkes et al., 2021). The corrected resampled *t*-test pre-defines the between-fold correlation to be the fraction of data in each test fold. If the true correlation is higher than the predefined correlation, FPR is inflated; if it is lower, power is reduced (Supplementary Methods S6). The 5×2 paired t-test and empirical test instead favor FPR control over power: the former discards information (Supplementary Methods S9), whereas the latter implicitly assumes a between-fold correlation of 0.5 (Supplementary Methods S10). As will be seen, our simulations confirmed these tradeoffs: the corrected resampled *t*-test did not consistently control the FPR, whereas the 5×2 paired *t*-test and empirical test consistently controlled the FPR at the cost of substantially lower power. Although far superior to tests that assume fold independence, existing correlation-aware tests retain a trade-off between reliable FPR control and statistical power.

This trade-off motivates SHARP, which re-designs cross-validation so that the quantities governing uncertainty can be estimated rather than approximated. In each of *J* repetitions, SHARP randomly partitions the dataset into two disjoint halves, and performs K-fold cross-validation independently within each half (Fig. 6a). Fold-level performance differences are averaged within each half, yielding one statistic per half. Because the two halves share no observations, the two statistics in a given repetition are independent, while statistics across repetitions remain correlated. This structure enables direct estimation of the split-half variance and across-repetition correlation, enabling inference without fixing either quantity *a priori* (see Methods). We will now evaluate SHARP’s false positive rate, statistical power and confidence interval coverage against existing tests.

**Figure 6.**
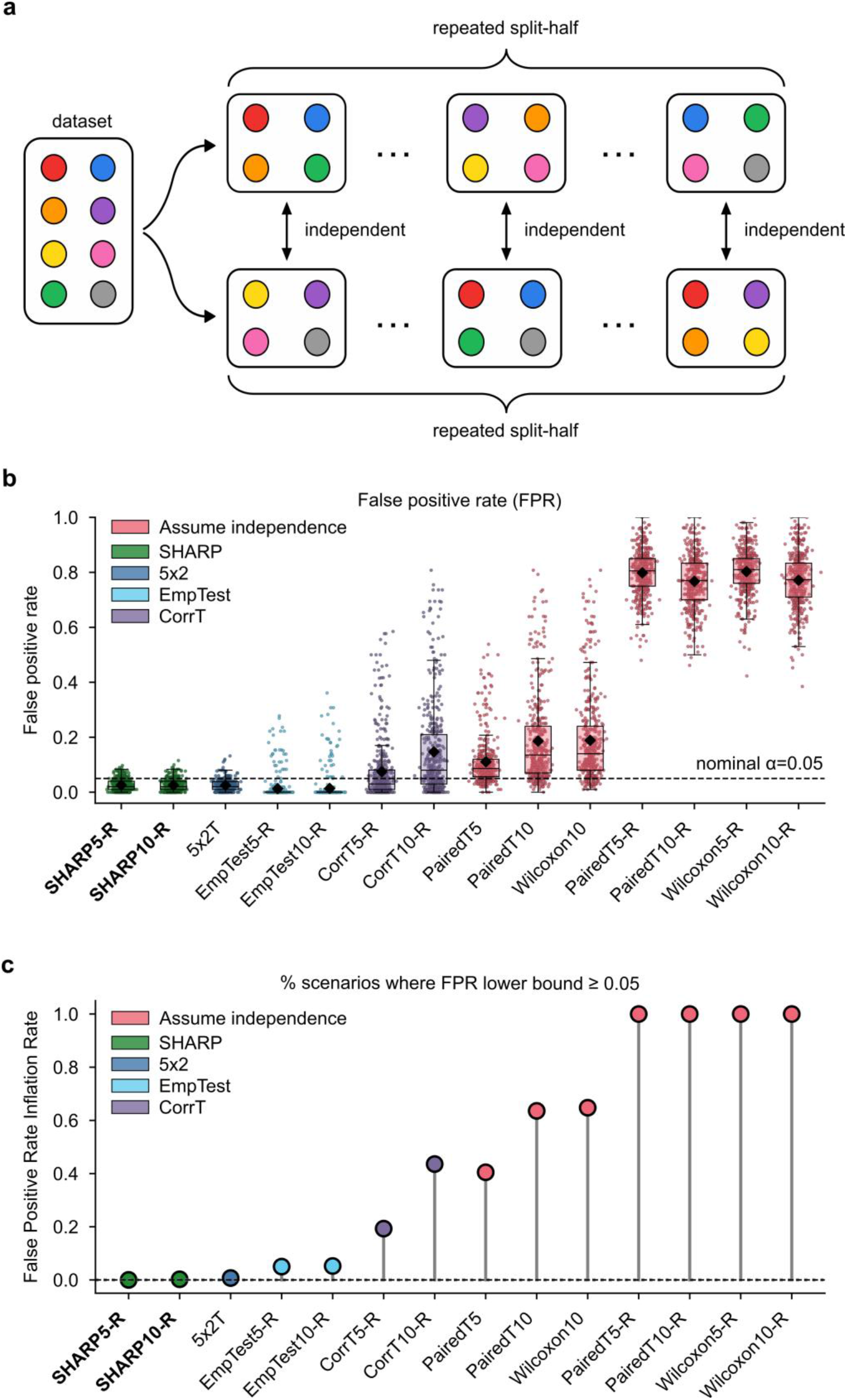
SHARP design and false positive rate control across 420 simulation scenarios. **a.** Schematic of SHARP. In each of *J* repetitions, the dataset is randomly split into disjoint halves, and K-fold cross-validation is performed independently within each half. Averaging fold-level differences within each half yields two independent split-half statistics per repetition; statistics across repetitions remain correlated because of data reuse. This structure permits estimation of the variance of the split-half statistic and the across-repetition correlation (see Methods). **b.** FPR for each statistical test, with one value per scenario (n = 420). Black dots mark mean FPR; dashed line marks the nominal level of 0.05. **c.** FPR inflation rate: percentage of the 420 scenarios in which the lower bound of the 95% confidence interval for FPR exceeded 0.05, indicating inadequate type I error control. **Test naming conventions.** Numeric suffixes “5” and “10” denote 5-fold and 10-fold cross-validation. The “-R” suffix indicates repeated cross-validation: 5-fold repeated 60 times or 10-fold repeated 30 times, yielding 300 fold-level statistics. For SHARP, each repetition yields one pair of split-half statistics, so SHARP5-R yields 60 pairs and SHARP10-R yields 30 pairs. Repetition number was chosen to ensure stabilization of the FPR of tests that account for between-fold correlation; tests assuming fold independence do not stabilize (Fig. 5d). “CorrT” and “EmpTest” denote the corrected resampled paired t-test (Nadeau & Bengio, 2003) and empirical test of differences (Parkes et al., 2021), respectively.

### SHARP: reliable FPR control through direct variance estimation

Across the same 420 simulation scenarios as the previous sections, the corrected resampled paired t-test, 5×2 paired t-test, and empirical test of differences markedly reduced FPR relative to tests assuming fold independence (Fig. 6b). The 5×2 paired t-test and empirical test controlled FPR at the nominal 5% level. The corrected resampled paired t-test was borderline under 5-fold cross-validation and failed under 10-fold, with FPR significantly exceeding 5% in 19% and 44% of scenarios, respectively (Fig. 6c). These conclusions held across datasets (Supplementary Fig. 3) and sample sizes (Supplementary Fig. 4). Under an alternative simulation scheme (Supplementary Methods S11), FPRs were lower, but tests assuming fold independence still failed to control FPR. The main qualitative difference was that the corrected resampled paired t-test controlled FPR under this scheme, indicating that its FPR control is scheme-dependent (Supplementary Fig. 5).

Across the same 420 scenarios, SHARP controlled FPR reliably (Fig. 6b), with CI lower bounds below 5% in nearly all scenarios (Fig. 6c), thereby matching the 5×2 paired t-test and the empirical test in FPR control. Conclusions were consistent across datasets (Supplementary Fig. 3), sample sizes (Supplementary Fig. 4), and the alternative simulation scheme (Supplementary Fig. 5). A broader comparison across additional tests confirmed the same pattern: tests that effectively ignored between-fold correlation had inflated FPR, while correlation-aware tests controlled FPR reliably (Supplementary Fig. 6).

### SHARP matches or exceeds the power of existing correlation-aware tests

To benchmark statistical power, we adapted the FPR simulation procedure, but instead of corrupting both copies of the sampled dataset (Fig. 5a), we corrupted only one copy (Supplementary Fig. 7). The model trained on the uncorrupted copy should have higher expected predictive performance by construction, so a sensitive test should reject the null hypothesis of equivalent performance. Within each scenario, power was estimated as the fraction of non-overlapping subsamples for which the null was rejected; the estimates were then averaged across the 420 scenarios.

SHARP5-R had the highest power, outperforming the 5×2 paired t-test by 27.8 percentage points and winning every pairwise comparison (purple row in Fig. 7a). Across a wider set of tests, SHARP remained the most sensitive, followed by the corrected resampled paired t-test (Supplementary Fig. 8). An alternative simulation scheme (Supplementary Fig. 9; Supplementary Methods S11) yielded the same overall pattern, except that SHARP5-R and CorrT5-R achieved similar power.

**Figure 7.**
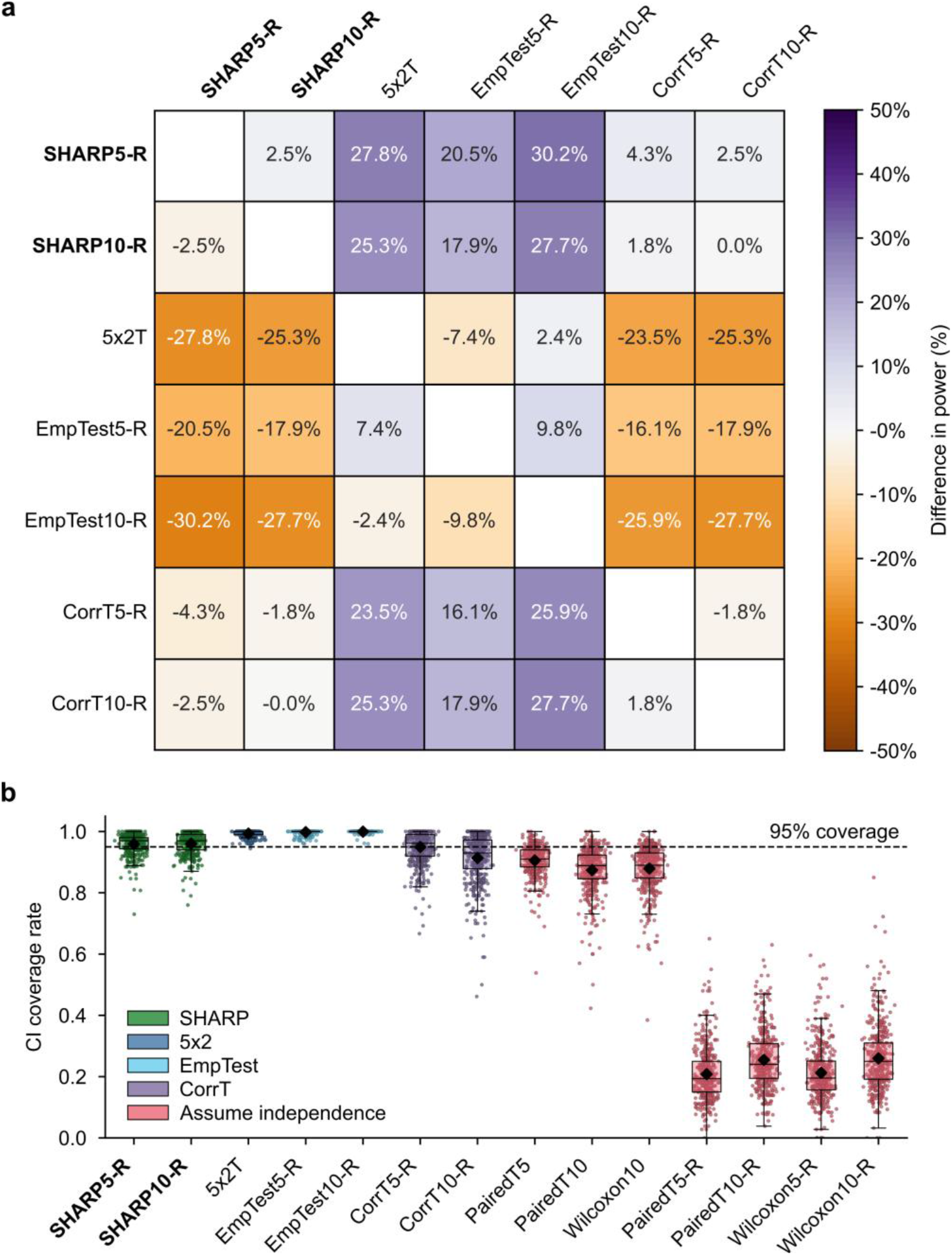
Statistical power and confidence interval coverage across 420 scenarios. **a**. Pairwise differences in power, computed as the power of the test (shown in the row) minus the power of the test (shown in the column), averaged across 420 scenarios. Purple indicates that the test in the row had higher power; orange indicates the reverse. For example, the purple cell in the first row and third column indicates that SHARP5-R has 27.8 percentage points higher power than the 5×2 paired t-test. Panel a shows only the correlation-aware tests from Fig. 6. SHARP5-R won every pairwise comparison, as reflected in its entirely purple row. **b.** 95% confidence interval (CI) coverage rate across 420 scenarios. For each scenario, the coverage rate was the percentage of non-overlapping subsamples in which the 95% CI contained the (estimated) true performance difference between models. For a well-calibrated test, the coverage rate should be ∼95% (black dashed line). Each boxplot contains 420 values: one coverage rate for each of 420 scenarios; black dots indicate means. Tests that assumed fold independence (red boxplots) had overly narrow CIs and coverage rates much lower than 95%.

### SHARP provides well-calibrated confidence intervals

Beyond binary significance decisions, confidence intervals quantify effect magnitude and precision: a statistically significant difference may be too small to matter practically, while a wide interval around a non-significant result may indicate insufficient power rather than equivalence.

For a well-calibrated test, a 95% confidence interval should contain the true performance difference between the two models ∼95% of the time (see Methods). Across the 420 power simulation scenarios, tests that assumed fold independence produced overly narrow CIs, with coverage well below 95% (Fig. 7b). The empirical test of differences and 5×2 paired t-test, which had low power (Fig. 7a), produced overly wide CIs with coverage approaching 100%.

The corrected resampled paired t-test achieved a mean coverage of 95% under 5-fold cross-validation but coverage varied widely across scenarios; this is consistent with its assumed between-fold correlation, which is fixed by the cross-validation design rather than estimated from the data. SHARP5-R and SHARP10-R both achieved mean coverage of 96%. Additional tests are shown in Supplementary Fig. 10. These conclusions also held under an alternative simulation scheme (Supplementary Fig. 11; Supplementary Methods S11), except that the corrected resampled paired t-test achieved stable coverage, mirroring its scheme-dependence FPR control.

Across FPR, power, and CI coverage, SHARP5-R provided the best balance among the tests examined. The corrected resampled paired t-test was a close second in power, but was less reliable in FPR control and CI coverage because it fixes rather than estimates fold-level correlation – an assumption that SHARP avoids. Fig. 8a consolidates FPR control, statistical power, and confidence-interval coverage across all tests.

**Figure 8.**
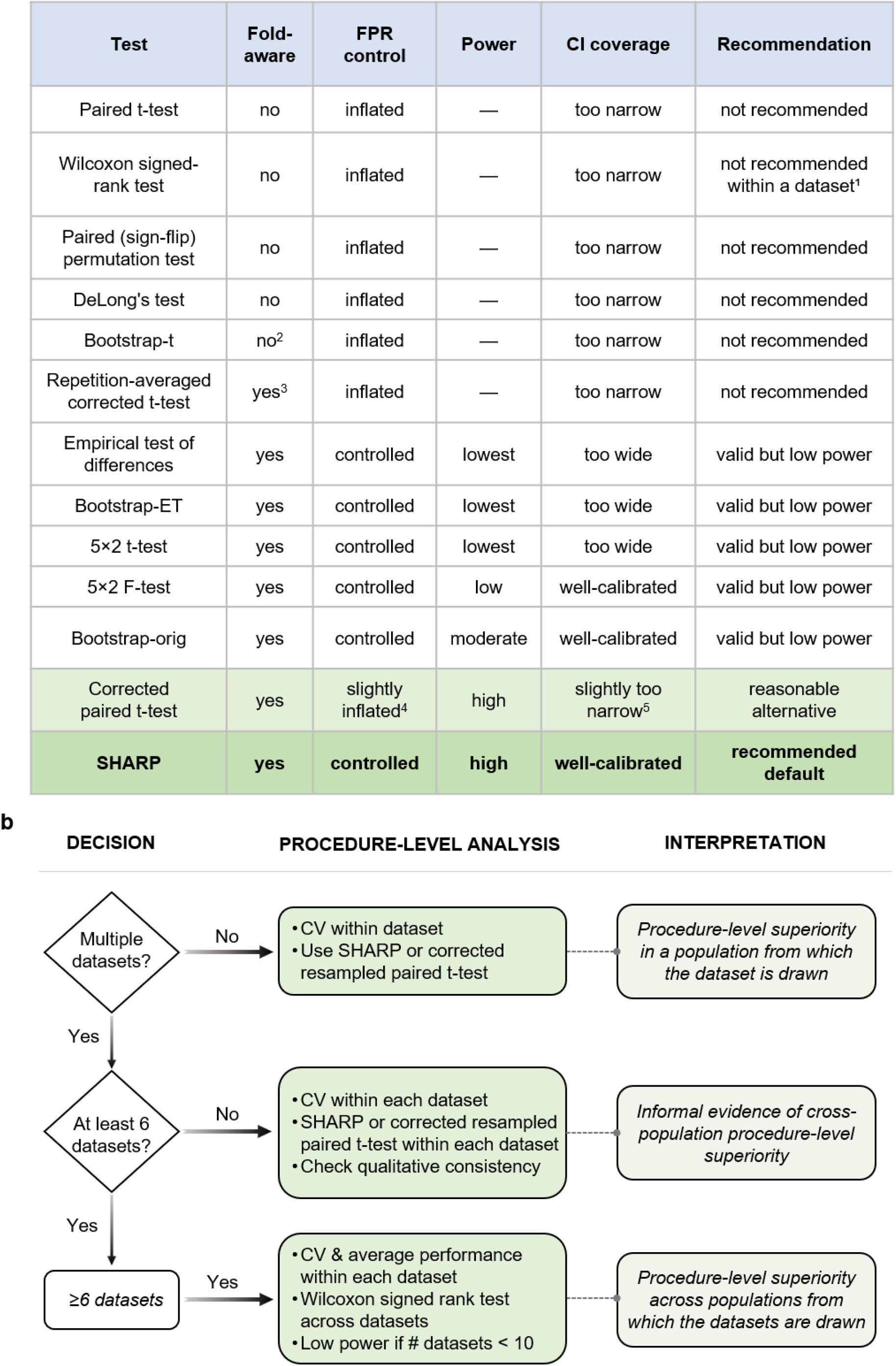
Guidelines for valid model comparison. **a.** Statistical properties of tests for comparing models in cross-validation. SHARP (highlighted) is recommended as the default; the corrected resampled paired t-test is a reasonable alternative, particularly when sample size makes SHARP’s split-half design less attractive. **Footnotes:** ^1^The Wilcoxon signed-rank test is invalid when applied to dependent fold-level performance differences within a dataset but valid for comparing independent dataset-level performance differences across datasets (Demšar, 2006; see panel b). ^2^Bootstrap-t resamples fold-level differences but wrongly treats the resulting bootstrap samples as independent, making the test invalid (see Methods). ^3^The repetition-averaged corrected resampled t-test uses an overly liberal assumption about between-fold correlation, leading to high FPR (see Supplementary Methods S8). ^4^FPR for the corrected resampled paired t-test is controlled in some regimes but mildly elevated in others. ^5^95% confidence interval for the corrected resampled paired t-test is well-calibrated in some regimes but slightly too narrow in others. Power is reported as “—” for tests that do not control FPR at the nominal level because power is not meaningful in this case. **b.** Decision framework for procedure-level inference based on the number of available independent datasets. The flowchart guides users through sequential decisions (left) to a recommended procedure-level analysis (middle) and the corresponding scope of inference (right). Although panel b depicts procedure-level inference, its dataset-count logic also applies to fixed-model external validation, where differences come from external evaluation rather than cross-validation (see main text).

## Discussion

### A serious threat to replicability in biomedical artificial intelligence (AI) research

The near-universal use of tests that assume fold independence is not merely a technicality. It occurred against a backdrop of pervasive deficiencies in statistical reporting: of 1,875 studies that compared models, 1,008 (54%) did not clearly apply a statistical test or report confidence intervals. Only 68 of the 173 at-risk studies permitted valid re-analysis without additional assumptions, yet 69% of this comparatively transparent subset contained a spurious comparison. By extrapolation, we estimate that spurious comparisons occur in nearly two in three studies and support abstract-level claims in nearly one in three studies. Thus, assuming fold independence has led to widespread overstatement of statistical evidence in biomedical AI research.

Importantly, between-fold correlation affects uncertainty quantification, not point estimation. Absent leakage or other circularity, cross-validated performance estimates remain valid estimates of generalization to the sampled population. This correlation instead invalidates independence-based estimates of their variance – and hence p-values, confidence intervals, and test statistics. Accordingly, our re-analysis does not establish that the null hypothesis is true for spurious comparisons or that the studies’ broader conclusions are false. Rather, it shows that the statistical evidence was substantially weaker than reported. Among studies with an abstract-level claim supported by a spurious comparison, the median corrected p-value was 0.17. Thus, many comparisons were not merely shifted just beyond the significance threshold.

The Open Science Collaboration provides a useful empirical benchmark: stronger original evidence predicted replication success, yet only 41% of studies with original p-values below 0.02 yielded a significant replication, compared with 26% with p-values between 0.02 and 0.04 (Open Science Collaboration, 2015). These figures do not estimate replication rates in biomedical AI, but show that findings in psychology supported by considerably stronger evidence frequently failed independent testing. The substantially weaker evidence in our re-analysis therefore raises serious concerns about replicability.

Our prevalence estimates require extrapolation from the 68 re-analyzed studies to the remaining 105 at-risk studies and beyond our corpus. Three considerations inform how this extrapolation should be interpreted. First, some comparisons could not be re-analyzed, including those evaluated with tests even more vulnerable to FPR inflation; the corrected resampled t-test does not always control the FPR; and we did not add multiple-comparison corrections retrospectively. Second, re-analysis required sufficiently complete reporting, data, or code, giving no reason to regard the remaining studies as methodologically safer. Third, although our corpus was restricted to journals with an impact factor ≥15, the use of tests assuming fold independence was not associated with impact factor, journal rigor score, or data and code availability. Together, these considerations suggest our estimates are likely conservative and existing safeguards are insufficient.

### Practical guidance for valid model comparison

Model comparison should be tailored to the scientific question. “Procedure-level inference” asks whether one algorithm, pipeline or biomarker would outperform another if model training were repeated with new samples from the population or populations of interest (Dietterich, 1998). By accounting for training-sample variation, procedure-level inference guides the choice among development procedures. “Fixed-model transportability” asks whether the advantage of an already-trained model persists across populations, sites or time periods, thereby informing deployment (Dietterich, 1998; Justice et al., 1999). The inferential question determines the analysis, while the number of available datasets determines whether inference is limited to the sampled population or extends across populations.

Cross-validation is commonly used for procedure-level inference, both within individual datasets and across multiple data sources. The appropriate inference strategy depends on the number of independent datasets available (Fig. 8). When only one dataset is available, we recommend SHARP or the corrected resampled paired t-test (Fig. 8b). A single dataset supports inference only about the population it represents. With two to five datasets, within-dataset testing with SHARP or corrected t-test, combined with qualitative consistency provides informal cross-population evidence (Fig. 8b). With six or more datasets, a Wilcoxon signed-rank test across independent dataset-level performance differences supports formal cross-population inference (Demšar, 2006). However, power is low with fewer than ten datasets, and access to these many comparable datasets is often infeasible in biomedical research.

Procedure-level performance under population shift can be examined using multiple-source cross-validation (Geras & Sutton, 2013), including leave-one-site-out (Takada et al., 2021) and leave-some-sites-out (Chen et al., 2022) cross-validation. Because models are refitted as held-out data sources change, these designs evaluate procedures rather than fixed-model transportability. Performance differences from multiple-source cross-validation remain statistically dependent through data reuse (Fig. 1b), unlike independent dataset-level differences (Fig. 8b). The corrected resampled t-test has been used in this setting (Chen et al., 2022). With enough data sources, SHARP can be extended by partitioning them into two disjoint groups and performing leave-one-source-out cross-validation within each group.

In contrast, fixed-model transportability is usually evaluated through independent external validation. Models are trained using one or more development datasets and evaluated without refitting in external datasets (Steyerberg & Harrell, 2016; Collins et al., 2024; Rosenblatt et al., 2026). Because models are not repeatedly fitted across overlapping training sets, cross-validation-induced correlation is absent and standard tests appropriate to the performance metric can be used. A single external validation dataset supports inference only for the population represented and does not by itself establish general transportability (Riley et al., 2016; la Roi-Teeuw et al., 2024). With multiple independent external datasets, the logic in Fig. 8b applies, except that performance differences come from fixed-model evaluation rather than cross-validation. With two to five datasets, within-dataset tests and qualitative consistency provide informal cross-population evidence. With six or more datasets, a Wilcoxon signed-rank test across dataset-level differences supports formal cross-population inference.

Researchers can address both questions sequentially: use procedure-level inference to select an algorithm, pipeline or biomarker, then fit the selected model and evaluate it in independent external populations. SHARP supports the first stage but not the second.

Finally, cross-population inference is often not the goal (Dietterich, 1998). Many domain-specific studies aim to identify the procedure that will generalize best to new observations drawn from the same population as those in a particular dataset. In such settings, a valid within-dataset test is not a compromise: it provides the required scope of inference. It should not, however, be interpreted as evidence that a fitted model will transport to other populations or settings.

### Minimum reporting standards for model comparison

Valid model comparison requires sufficient reporting. Of 1,875 studies (Fig. 2a), 54% did not clearly apply a statistical test or report confidence intervals: in some, no formal inference was conducted; in others, reporting was too vague to tell. Many remaining studies lacked enough detail to assess validity. This constrained our re-analysis: despite consulting shared data and code, only 68 of 173 at-risk studies could be re-analyzed. Even basic details sometimes differed between manuscripts and the shared code, such as whether cross-validation used five or ten folds.

Studies comparing models with cross-validation should report: (1) the number of folds, (2) the number of repetitions, (3) whether the models were evaluated using corresponding cross-validation splits, (4) the specific statistical test used, including whether it is one-or two-sided, and (5) how fold-level statistics are aggregated, including the dimensions of the input to the statistical test.

The unit of analysis is particularly important. Repeating 5-fold cross-validation 100 times yields 500 fold-averaged differences, or 100 repetition-averaged differences if fold differences are averaged within each repetition. These choices are not interchangeable: applying the corrected resampled t-test to repetition-averaged differences massively inflates the FPR (Supplementary Fig. 6; Supplementary Methods S8).

For bootstrap procedures, authors should report whether bootstrapping generates training and test sets or only resamples cross-validation outputs, then explain how the resulting samples are used to compute p-values or confidence intervals. Treating bootstrap samples as independent observations massively inflates the FPR, whereas using their empirical distribution yields valid inference (Supplementary Fig. 6), albeit with low power (Supplementary Fig. 8).

Confidence intervals require similar specificity. We excluded all studies reporting only confidence intervals because too few explained how they were computed. An interval based on the standard error of fold-averaged statistics incorrectly assumes independence and is too narrow, while one based on their 2.5th and 97.5th percentiles is overly conservative. Authors should therefore report the underlying variance estimator or resampling scheme.

### Practical trade-offs between SHARP and the corrected resampled t-test

Across the regimes examined, SHARP provided the best overall balance of FPR control, power, and CI coverage. Its split-half step produces pairs of independent statistics, allowing variance and correlation to be estimated rather than assumed. The cost is that cross-validation is performed separately within each half, so relative model performance may differ from full-dataset estimates, particularly when algorithms or biomarkers scale differently with training size.

The corrected resampled *t*-test retains the full dataset and uses standard cross-validation outputs but fixes rather than estimates the between-fold correlation. Its power was similar to or slightly below SHARP, but its FPR control and CI coverage were simulation-scheme-dependent.

SHARP is available as an open-source, scikit-learn-compatible implementation requiring minimal changes to existing workflows. We therefore recommend SHARP when sample size supports its split-half design and the corrected resampled *t*-test as a reasonable alternative otherwise.

### A tractable threat to replicability

Ignoring correlation across folds threatens replicability. The threat stems from a structural property of cross-validation, whereas p-hacking and selective reporting arise from researcher incentives or analytical flexibility. Unlike underpowered studies, however, this problem is unusually tractable: adopting appropriate statistical tests or evaluation procedures requires no changes to data collection or study design. Given its near-universal prevalence and substantial consequences for many published claims, accounting for between-fold correlation offers an immediate opportunity to strengthen biomedical artificial intelligence.

## Methods

### Cross-validation primer

In this section, we provide an overview of different variants of cross-validation. In K-fold cross-validation, a dataset is divided into *K* non-overlapping partitions of data samples (folds). For each of *K* iterations, one fold is treated as the test set, while training is performed on the remaining folds. The procedure is repeated *K* times, so each fold has its turn to be the test. When comparing two models, the entire K-fold cross-validation procedure can be applied to each model separately. The *K* partitions are set up to be the same across the two runs of K-fold cross-validation, so that there is fold-level correspondence between the two models.

Within each fold, the performance of each model can be evaluated on the test samples, yielding *K* pairs of performance estimates for the two models. Taking the difference within each pair produces a vector ***D*** of *J* performance differences, where *J = K*. We note that the *J* performance differences are not independent because training sets overlap across iterations (for *K* > 2) and the test set from one iteration contributes to the training set of all other iterations. Many studies apply a statistical test to the vector ***D*** of *J* performance differences to evaluate the null hypothesis of equivalent prediction performance between the two models.

For stability (Bouckaert & Frank, 2004; Varoquaux et al., 2017), many studies also repeat K-fold cross-validation *R* times. In this scenario, we still obtain a vector ***D*** of *J* performance differences, but *J = K* × *R*. Fold-averaged statistics are correlated both within and across repetitions. Within each repetition, training sets overlap across folds. Across repetitions, both training and test sets can overlap, inducing a potentially different correlation. Once again, many studies apply a statistical test to the vector ***D*** of *J* performance differences to evaluate the null hypothesis of equivalent performance between the two models.

In Monte Carlo cross-validation, the data points that make up a dataset are repeatedly divided into training and test sets randomly. If the performance metric is averaged across samples in the test set and this procedure is repeated *J* times, we obtain a vector ***D*** of *J* performance differences. In this setup, there is overlap between both training and test sets across the *J* repeats, so the fold-level statistics are non-independent. Similar to the previous setups, many studies apply a statistical test to the vector ***D*** of *J* performance differences to evaluate the null hypothesis of equivalent performance between the two models.

In the above cross-validation schemes, model performance is computed separately within each test fold, yielding one performance-difference statistic per fold for the statistical test. We refer to these as fold-averaged statistics, including when a metric such as AUC is computed over, rather than literally averaged across, the test samples. During our meta-analysis, we observed some studies using “sample-level” instead of “fold-averaged” statistics. To illustrate the distinction, consider a study performing a single instance of 10-fold cross-validation on a dataset of 1,000 independent samples. For a performance metric defined at the sample level, a fold-averaged approach averages performance differences within each fold, yielding 10 performance differences for the statistical test, i.e., the vector ***D*** has 10 values. A sample-level approach forgoes this aggregation and instead enters all 1,000 performance differences directly into the statistical test, i.e., the vector ***D*** has 1000 values. Correlations among performance differences in the same fold might be stronger than correlations between folds (since predictions within each fold are generated by the same trained model), so sample-level statistics might further inflate the false positive rate.

A final variant aggregates the fold-averaged statistics in the opposite direction: rather than disaggregating to individual samples, performance differences are averaged across folds within each repetition of the cross-validation, so that each repetition contributes a single value to the statistical test. We refer to these as “repetition-averaged” statistics, in parallel to the “fold-averaged” and “sample-level” statistics above. For example, if 10-fold cross-validation is repeated 30 times, a fold-averaged approach yields a vector ***D*** of length 300, whereas a repetition-averaged approach first averages the 10 fold-averaged performance differences within each repeat and then enters the resulting 30 values into the statistical test, i.e., the vector ***D*** has 30 values. The correlation among the 30 repetition-averaged statistics is larger than the correlation among the 300 fold-averaged statistics. Feeding this repetition-averaged vector into the corrected resampled t-test (Nadeau & Bengio, 2003; Supplementary Methods S5.5) – which is calibrated for between-fold correlation, not the larger between-repetition correlation – leads to a high false positive rate (Supplementary Methods S8).

### Meta-analysis

#### Journal selection and search strategy

We conducted a literature search in PubMed on 18 September 2025 to identify original research articles that compared machine learning models using cross-validation. The search was restricted to articles published between 1 June 2020 and 1 June 2025. Eligible journals were defined using Journal Citation Reports (JCR) and were required to have a Journal Impact Factor (JIF) ≥ 15 in either 2023 or 2024. Journals classified as review journals were excluded.

For each candidate journal, ISSN and eISSN information was obtained from JCR and matched to the corresponding National Library of Medicine (NLM) journal abbreviation in the NLM Catalog. Journals not indexed in the NLM Catalog were excluded, because PubMed searches by journal required this identifier. For each eligible journal, the following PubMed query was then executed: (“outperform” OR “outperforms” OR “state-of-the-art” OR “benchmark” OR “benchmarking” OR “algorithm comparison” OR “model comparison” OR “performance evaluation” OR “method comparison”) AND (“cross validation” OR “cross-validation” OR “crossvalidation”) AND (“2020/06/01”[Publication Date]: “2025/06/01”[Publication Date]) AND (Journal Abbreviation [TA])

This search yielded 2,400 studies for screening (Fig. 2a).

#### PRISMA workflow and criteria

Screening of the 2,400 studies followed PRISMA guidelines (Page et al., 2021). A final set of 184 studies met the following four criteria.

##### 1. Direct model performance comparison

To satisfy this criterion, the study had to compare at least two distinct models – that might differ in terms of algorithms, pipelines, hyperparameter configurations, feature sets (e.g., biomarkers), or model-selection procedures – on the same prediction target. Eligible studies must compare models based on “direct” performance metrics (e.g., accuracy, AUC, MSE, MAE, correlation, etc.) computed from comparing predictions against ground truth. The models must be evaluated on the same test data. This criterion resulted in 1875 papers involving model comparisons and 525 without model comparisons (Fig. 2a).

We excluded studies that compared only downstream quantities derived from predictions, as opposed to direct performance metrics. For example, a study might develop two brain-age models from MRI and compare them based on which model’s brain age predictions better differentiate Alzheimer’s disease from healthy individuals, as opposed to comparing models’ accuracy in predicting the training target (chronological age). Although between-fold correlation would also arise in such cases, we excluded these studies because it was difficult to have a clear set of criteria for what downstream analyses to include or exclude in the meta-analysis.

##### 2. Use of statistical inference for the comparison

The study had to report at least one p-value or confidence interval for a quantitative comparison of machine learning models. Studies that only reported descriptive statistics or performed statistical tests unrelated to model comparison (e.g., correlation between model predictions and confounding variables) were considered ineligible. This criterion resulted in 867 studies using statistical inference for comparing models, while 1008 studies were eliminated.

##### 3. Fold-based statistical inference

Among the 867 studies, some did not use cross-validation, and a large fraction did not clearly describe their inference procedure. To be included in our final analyses, a study had to clearly use what we broadly refer to as “fold-level statistics” to compute a statistical test or confidence interval. This term encompasses sample-level, fold-averaged, and repetition-averaged statistics derived from cross-validation test folds (see Methods “Cross-validation primer”). For example, we included studies that (i) performed K-fold cross-validation and used *K* pairs of fold-averaged performance values for statistical inference; or (ii) conducted *J* repeated random train–test splits (i.e., Monte Carlo cross-validation) and used *J* pairs of values for statistical inference. There were also studies that repeated K-fold cross-validation multiple times. For example, a study that repeated 10-fold cross-validation 100 times and used 1,000 pairs of fold-averaged performance values for a t-test, would be included in the meta-analysis. If such studies averaged performance across the 10 folds within each repetition and used the resulting 100 pairs of repetition-averaged values in a t-test, they would also be included. For studies reporting p-values, they also had to clearly state the name of the statistical test. This criterion resulted in a set of 239 studies.

##### 4. Use of tests that ignore or account for between-fold correlation

The reported procedure had to be sufficiently clear, so we could classify it as either ignoring or accounting for between-fold correlation. Of the 239 studies, one employed a permutation test that was invalid for other reasons, so we considered it outside the scope of the current study (see Supplementary Results for details). We also excluded studies that reported only confidence intervals, as most did not explain how they were computed, a problem that has been independently documented in medical imaging (André et al., 2026). A 95% confidence interval based on the standard error of fold-level statistics would incorrectly assume fold independence, yielding an interval that is too narrow. Conversely, a 95% interval computed from the 2.5th and 97.5th percentiles of fold-level statistics approximates the empirical test of differences (Supplementary Methods S5.7) and would be overly conservative. Without further methodological details, these studies could not be reliably assessed. Applying these criteria resulted in a final set of 184 studies.

The PRISMA flow diagram describing identification, screening, eligibility assessment and inclusion is shown in Fig. 2a.

#### Scientific field classification

To explore the prevalence of invalid statistical inference across scientific fields, we assigned each study to a field defined using the Web of Science Journal Citation Reports (JCR) subject categories. JCR listed 254 categories, and each journal could belong to one or more categories. Because our PubMed-based search targeted biomedical and life-science content, not all JCR categories were represented (Supplementary Table S2). Studies in journals assigned to one scientific category inherited that category. For journals assigned to multiple categories (e.g., Nature), we used a large language model (LLM) to determine the most relevant category for each study (see next section).

#### LLM-assisted screening and manual human verification

To scale screening and data extraction, we used an LLM (Claude Opus 4.1, Anthropic) to assist human raters. For each of 2,400 studies, we provided the main text scraped from PubMed, including figure captions, but excluding figures, tables and supplementary material, together with a structured prompt. The LLM answered five questions: (Q1) scientific field classification; (Q2) whether the study directly compared model performance (PRISMA criterion 1); (Q3) whether statistical inference was used for model comparison (PRISMA criterion 2); (Q4) details of the statistical procedure; and (Q5) whether statistical inference was based on values derived from cross-validation folds (PRISMA criterion 3).

The detailed manual verification procedures and full prompt are provided in Supplementary Methods S1.1 and S1.2, respectively. Briefly, for each of Q1–Q3, two human raters (TZ and HL) independently assessed a random sample of 50 studies and resolved discrepancies by consensus. The LLM agreed with the consensus classification for all 50 studies for Q1 and Q2. Agreement for Q3 was 39 of 50 studies; the errors comprised one false negative and ten false positives. Given the false positives, two human raters (TZ, HL or SZ) independently reviewed the full text of all 1,197 studies classified as eligible by the LLM and resolved discrepancies by consensus. For Q4 and Q5, the LLM outputs served only as references; two human raters (TZ, HL or SZ) independently re-derived the information for every study and resolved discrepancies by consensus. The human-verified Q5 information was used to assess PRISMA criterion 3, while the human-verified Q4 and Q5 information together were used to assess PRISMA criterion 4.

This procedure initially identified 210 studies satisfying all four PRISMA criteria. We re-screened these studies using Claude Opus 4.8 and 5 with access to the full text and, where available, released code and supplementary files. Two raters (TZ and HL) independently re-assessed each study and resolved discrepancies by consensus. The Q4 and Q5 information was updated for some studies during this re-screening. This further excluded 26 studies, counted among the 628 exclusions under PRISMA criterion 3 in Fig. 2a, leaving 184 studies. For each included study, the human-verified Q4 and Q5 information was used to determine whether at least one test assumed independence across folds or all tests accounted for between-fold correlation.

#### Journal policy rigor scoring

To evaluate whether journal guidelines might influence the prevalence of invalid statistical inference, we defined a journal-level “policy rigor” score comprising two domains:

1. Transparency and reproducibility
  - *Code availability policy*: requirement or encouragement to share analysis code or model scripts.
  - *Data availability policy*: requirement or encouragement to share underlying or processed data.
  - *Data/code availability statement requirement*: requirement or encouragement for a formal data and/or code availability statement in the manuscript.
  - *Code for peer review*: requirement or encouragement to provide custom code to editors and reviewers during peer review.
2. Statistical rigor and methodological guidance
  - *Statistical test guidance*: explicit guidance on appropriate statistical testing (for example, paired vs unpaired tests, reporting of uncertainty, comparison under cross-validation).
  - *Statistical review*: whether dedicated statistical review is part of the editorial or peer-review process.
  - *Reporting standards or checklists*: requirement or encouragement to follow established methodological reporting guidelines (for example, TRIPOD-AI, CONSORT-AI, PRISMA).

For transparency and reproducibility, journals were scored on a three-point scale for each of the four criteria: 0 = not mentioned or no guidance; 1 = encouraged or optional; 2 = mandatory or strongly enforced. The domain score ranged from 0 to 8.

For statistical rigor and methodological guidance, journals were also scored on a three-point scale for each criterion with 0 corresponding to the least rigor and 2 corresponding with the most rigor. See Supplementary Table S1 for the scoring criteria. The domain score ranged from 0 to 6.

To derive the scores, we used an LLM (GPT-5, OpenAI) to search each journal’s official webpages (Instructions for Authors, Editorial Policies, Submission Guidelines, Reporting Standards, and related pages) on November 7, 2025, to identify the most authoritative policy statements. The LLM prompt is provided in Supplementary Methods S1.3.

Two human raters (TZ and HL) verified the LLM-generated scores. Any discrepancies between raters were resolved by discussion and consensus. Finally, scores were summed across the two domains, resulting in an overall journal rigor score ranging from 0 to 14.

### Study-level code and data availability

For each study, we used an LLM (Claude Opus 4.1, Anthropic) to determine whether the study provided data or code. Similar to the main meta-analysis, for each of 184 studies, we provided the main text scraped from PubMed, including figure captions, but excluding figures, tables and supplementary material, together with a structured prompt. The full prompt text is provided in Supplementary Methods S1.4. In a random audit of 50 studies, two human raters (TZ and HL) independently assessed each study. Any discrepancies between raters were resolved by discussion and consensus. The LLM agreed with the consensus human classification for all 50 studies. Therefore, we used the LLM classifications in the analyses presented in Supplementary Fig. 1.

### Test of temporal trend & associations with scientific field, impact factor & scientific rigor

We tested for a trend in the number of included studies over publication time. Letting *Y_i_* be the number of studies published in the *i*-th 6-month period, we fit a log-linear Poisson regression as follows:

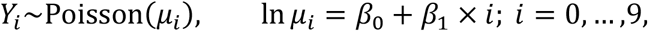

where *β*_1_ quantifies the trend over time and *β*_0_ is the intercept. If *β*_1_ = 0, then this would suggest that the number of studies stayed constant over time. If *β*_1_ is positive, then this means that volume of relevant studies is increasing. For example, in the current study, we found that *β*_1_ = 0.07 (*p* = 9e-3), so *e*^0.07^ = 1.070. This suggests that the volume of relevant studies increased by approximately 7.0% per 6-month period, or 14.5% per year. The Poisson regression model was fitted using python package statsmodels 0.14.5 with a p-value based on a Wald test.

We tested whether ignoring between-fold correlation was associated with publication period, impact-factor bin, journal rigor score, scientific field, code availability, or data availability. For ordinal factors – publication period, impact factor bin and journal rigor score – we used a permutation-based Cochran–Armitage trend test. We did not use logistic regression because the proportions of studies assuming fold independence were close to one, leading to a high prevalence of zero-and low-rate cells. The Cochran–Armitage test is sensitive to monotonic differences in the rate of studies that assumed fold independence. Specifically, we constructed a *N_rows_* × 2 contingency table in which rows indexed ordered strata (time intervals, impact-factor bins, or rigor levels) and columns counted studies that ignored versus accounted for between-fold correlation. Under the null hypothesis of no monotonic trend, we generated an empirical null distribution by repeatedly sampling 100,000 contingency tables conditional on the observed marginal totals (fixed row and column sums). For each sampled table we computed the Cochran–Armitage trend statistic. The two-sided p-value was estimated as the proportion of sampled Cochran-Armitage trend statistics whose absolute value was greater than or equal to the absolute value of the observed statistic.

For nominal factors – scientific field, study-level code availability or study-level data availability – we used a permutation-based analogue of the Fisher–Freeman–Halton exact test. We first constructed a *N_rows_* × 2 contingency table with rows representing levels of the nominal factor and the same two outcome columns as above. Under the null hypothesis of no association, we generated 100,000 tables conditional on the observed margins. For each sampled table, we computed the Pearson *χ*^2^ statistic as a measure of deviation from independence, forming an empirical null distribution. The p-value was estimated as the proportion of permuted *χ*^2^ values greater than or equal to the observed *χ*^2^.

### Re-analysis: loss of significance under a valid statistical test

To assess the practical consequences of assuming fold independence, we re-analyzed originally significant model comparisons reported by the 184 studies and determined whether they retained significance under a valid statistical test. 173 (out of the 184) studies reported at least one significant comparison based on a test that assumed fold independence, so the re-analysis focused on these 173 studies.

We assigned each originally significant comparison in the 173 studies to one of four recovery tiers (A, A′, B or C), based on whether missing information necessitated additional assumptions and whether a valid re-analysis was feasible under the study’s analytical setup.

A study could contribute multiple comparisons and comparisons from more than one tier. The primary re-analysis included the 68 studies that contributed at least one Tier A or Tier A′ comparison.

For Tier A comparisons, predictions for each test sample across all cross-validation splits (together with ground-truth labels) or fold-averaged performance values were available. For comparisons based on a single dataset, we extracted or computed fold-averaged performance values regardless of whether the original study had used sample-level or fold-averaged statistics, and applied the corrected resampled paired t-test.

Some Tier A comparisons were based on cross-validation performed separately on multiple independent datasets. The original studies had concatenated fold-averaged performance values across datasets and applied a paired test to the concatenated values, thereby ignoring between-fold correlation within each dataset. We instead averaged the performance differences across folds within each dataset and treated datasets as independent units (Fig. 8b). For comparisons with at least 6 datasets, we applied both a paired *t*-test and a Wilcoxon signed-rank test across datasets and classified the comparison as retaining significance if either test yielded *p* < 0.05. For the remaining comparisons, we applied only the paired *t*-test because a two-sided Wilcoxon signed-rank test cannot attain significance at *p* < 0.05 with fewer than six datasets.

For Tier A′ comparisons, predictions and fold-averaged performance values were unavailable, but the original analysis had clearly used a paired *t*-test or paired *z*-test on fold-averaged statistics from a single dataset and had reported an exact *p*-value and the exact cross-validation procedure. The degrees of freedom were also available for *t*-tests. We recovered the naïve statistic implied by the reported *p*-value and divided it by the Nadeau– Bengio correction factor 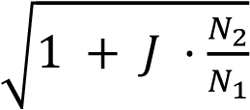, where *J* is the number of fold-averaged observations entering the test and *N*_2_/*N*_1_ is the ratio of test-set size to training-set size for each cross-validation split (Supplementary Methods S2). We then obtained the corrected p-value from the corresponding reference distribution. This is equivalent to applying the corrected resampled *t*-test or *z*-test to the unavailable fold-averaged statistics, without additional assumptions arising from missing information.

Tier B comparisons could be re-analyzed only under additional assumptions arising from missing information. Tier C comparisons could not be validly re-analyzed, either because the necessary information could not be recovered even under reasonable assumptions or because the cross-validation and testing setup did not permit a valid re-analysis. Neither Tier B nor Tier C comparisons entered the primary re-analysis, although a study included through a Tier A or Tier A′ comparison could also contribute comparisons from these tiers. Supplementary Methods S2 provides the complete tier definitions and re-analysis rules.

An originally significant comparison was classified as spurious if re-analysis yielded p ≥ 0.05 and as retaining significance if it yielded p < 0.05. A study was classified as containing a spurious comparison if any re-analyzed comparison was spurious; otherwise, it was classified as containing no spurious comparisons. Because Tier B and Tier C comparisons could not be re-analyzed, the latter classification established only that no re-analyzed comparison was spurious – not that every comparison in the study would have retained significance under valid testing. Furthermore, we generally did not adjust for multiple comparisons. In the few instances in which the original study had applied a multiple comparisons correction, we retained it only if we could reproduce the exact procedure. Omitting such corrections makes comparisons more likely to retain significance and therefore renders our estimate of significance loss conservative.

Among the 68 re-analyzed studies, we additionally examined whether claims made in each study’s abstract were supported by tests that assumed fold independence. If any supporting comparison was spurious, we classified the abstract as containing a claim supported by a spurious comparison.

No machine-extracted value entered the re-analysis without human verification. Two human raters (TZ and HL) independently confirmed key properties of the statistical test – including its reported p-value, one-sided (or two-sided), paired (or unpaired) and exact cross-validation procedure – against the original study and code where available. Discrepancies were resolved through discussion and consensus.

## Datasets

We used four datasets spanning image recognition, neuroimaging, ecological classification and systems biology. For EMNIST Digits (Cohen et al., 2017), we used 240,000 grayscale images to perform ten-class digit classification. For UK Biobank (Alfaro-Almagro et al., 2018), we used 101 regional gray-matter volumes from 36,000 participants to predict chronological age. The Covertype dataset (Blackard, 1998) comprised 581,012 samples with 54 cartographic features; we converted its seven-class outcome into a binary classification task distinguishing cover type 2 from all other types. The KEGG Metabolic Pathway dataset (Naeem & Asghar, 2011) comprised 53,414 samples with 20 graph-topological features, from which we predicted clustering coefficient. Detailed dataset descriptions, preprocessing and dataset-specific simulation settings are provided in Supplementary Methods S4.

## Simulations to evaluate FPR, statistical power and confidence intervals

### FPR simulation

We conducted analyses on four datasets — EMNIST, UKB, Covertype, and KEGG Metabolic Pathway — using four sample sizes: *N* = 100, 500, 1,000, and 2,000. For EMNIST, *N* = 100 was excluded because with 10 digit classes, it was not possible to guarantee at least one instance of each class in the test set under our proposed SHARP test, which required a split-half step within the cross-validation scheme (see Methods “SHARP test”). For UKB, *N* = 2,000 was excluded because the maximum number of non-overlapping sampled datasets that could be drawn from UKB at this sample size was fewer than 20 (i.e., 36,000 / 2,000 = 18), which we considered too few to yield stable false positive rate estimates.

We drew *m* non-overlapping sampled datasets of size *N* from the full dataset. The value of *m* was set to 100 when feasible; otherwise, it was set to the maximum number of non-overlapping sampled datasets that could be drawn. For example, for the UK Biobank dataset at *N* = 1,000, this yielded *m* = 36 (i.e., 36,000 / 1,000).

For each sampled dataset of size *N*, two noisy versions were generated by independently injecting the same level of random noise. Noise was introduced by permuting the target variable (e.g., digit labels for EMNIST) for a fixed proportion of samples. These were referred to as Noisy Dataset 1 and Noisy Dataset 2. We tested 10 permutation percentages in total: 10% to 100% with an increment of 10%.

For each noisy version of the sampled dataset of size *N*, we applied (i) a single instance of K-fold cross-validation or (ii) multiple repeats of K-fold cross-validation or (iii) Monte Carlo cross-validation. In classification tasks, stratified splitting was used to preserve label proportions across folds. All data splits were kept identical across the original sampled dataset and the two noisy datasets to ensure correspondence for paired statistical tests. We note that the SHARP test utilized a different cross-validation scheme.

A single algorithm was selected for each dataset based on initial exploration using the PyCaret Python package (Ali, 2020): logistic regression classifier for EMNIST, ridge regression for UK Biobank, Extra Trees classifier for Covertype, and Extra Trees regression for KEGG Metabolic Pathway (see Methods “Simulation settings”). For each noisy version of the sampled dataset, the algorithm was trained on the training folds and evaluated on the test fold of the original (unpermuted) sampled dataset. Evaluating both models on the same unpermuted test data ensured fair comparison between the two noisy-trained models.

For every training-test split, a pair of fold-level performance metrics were computed for each pair of noisy datasets, and the model performance differences between the two noisy datasets were computed, resulting in a vector ***D*** comprising *J* fold-level differences (except for SHARP; see Methods “SHARP test”). For example, if K-fold cross-validation was repeated *R* times, then vector ***D*** will be of length *J = K×R*. For a given two-sided statistical test (e.g., paired t-test), the vector ***D*** was used to evaluate the null hypothesis *μ* = 0, where *μ* is the true expected difference between the pair of trained models.

Since the noise level and algorithm were held constant across the pair of noisy datasets, the two trained models had identical expected predictive performance. Any rejection of the null hypothesis constituted a false positive. The false positive rate (FPR) was estimated as the fraction of *m* sampled datasets in which the null hypothesis was rejected at a significance threshold of 0.05. A well-calibrated test should reject the null hypothesis no more than 5% of the time. Recall that the *m* datasets were non-overlapping, so traditional binomial methods can be used to compute confidence intervals for the FPR. Specifically, the 95% confidence interval for the FPR was computed using the Wilson score interval with continuity correction (Newcombe, 1998); the formula is provided in Supplementary Methods S3.

Depending on the analysis, K-fold cross-validation was either performed once (*R* = 1) or repeated *R* times. When repeated, we targeted a total of 300 folds (*K × R* = 300) to ensure convergence of the statistical tests — for example, *R* = 30 for 10-fold and *R* = 60 for 5-fold cross-validation, so *J* was 300. For Monte Carlo cross-validation, we always used an 80/20 train-test split repeated 300 times, so *J* was again 300. The exception was the 5×2 statistical tests, which by design required 2-fold cross-validation repeated 5 times, so *J* = 10.

We note that given 10 noise levels (10% to 100%) and sample size (*N* = 100 to 2,000) across four datasets, there were 140 conditions. For each condition, we considered three hyperparameter-selection strategies: (1) fixed hyperparameters, (2) hyperparameter selection via a single training-validation split of the training data, and (3) nested cross-validation, in which hyperparameters were selected using cross-validation within the training data (see Methods “Simulation settings”). This resulted in 140 × 3 = 420 simulation scenarios in total. The FPR results are reported in Figs 5 and 6.

### Power simulation

The simulations to benchmark statistical power followed the same procedure as those for FPR (previous section), with one key difference. For FPR, the same algorithm was trained on two equivalently noisy datasets, so that the resulting pair of models was expected to have equivalent predictive performance. To assess statistical power, the same algorithm was instead trained on one noisy dataset and one original (unpermuted) sampled dataset. In this case, the model trained on the original (clean) sampled dataset was expected to outperform the model trained on the noisy dataset. A sensitive test should reject the null hypothesis *H*_0_: *μ* = 0 in a two-tailed test, where *μ* is the true expected difference in predictive performance between the two trained models.

As in the FPR case, power was estimated as the fraction of *m* datasets in which the null hypothesis was rejected at a significance threshold of 0.05. Crucially, a rejection was counted as a true positive only when the original-dataset model was found to significantly outperform the noisy-dataset model; cases in which the test rejected the null in the opposite direction were not counted as successes. Power was then averaged across 420 scenarios, yielding an average power for each statistical test.

### Confidence Intervals for Performance Difference

While there are different methods to compute a confidence interval (CI), here we used test-inversion which defines the CI by the set of null-hypothesis parameter values that cannot be rejected.

Recall that there were 420 simulation scenarios with each scenario corresponding to a given dataset (e.g., EMNIST), sample size (e.g., 2000), noise permutation percentage (e.g., 20%) and hyperparameter setting (e.g., fixed hyperparameters). For each scenario, there were *m* (non-overlapping) sampled datasets. For each sampled dataset, we injected noise to it, resulting in a noisy dataset. In the power simulations, the same algorithm was trained on one noisy dataset and one original (unpermuted) sampled dataset. The model trained on the original sampled dataset was expected to outperform the model trained on the noisy data.

Following the power simulations (previous section), for a given pair of original and noisy datasets, we performed cross-validation to obtain a vector ***D*** of *J* fold-level prediction performance differences, with mean *D^-^*. We applied various statistical tests to the vector ***D*** to evaluate the null hypothesis *μ* = 0, where *μ* is the true expected difference between the pair of trained models.

To obtain the 95% confidence interval for *μ* for a given statistical test (e.g., paired t-test), we tested null hypotheses *μ* = *μ*_0_ for different values of *μ*_0_ to discover the range of values such that *p* ≥ 0.05. This search was achieved by starting with *μ*_0_ = *D^-^*, and performing a binary search in the direction larger than *D^-^* and in the direction smaller than *D^-^*. The set of non-significant parameter values, [*μ_L_*, *μ_U_*], is the 95% confidence interval for the performance difference between the two models.

Ideally, we would evaluate the accuracy of the computed 95% confidence intervals by comparing with the true difference *μ*_true_, but this is not observable. Instead, we averaged *D^-^* across all *m* non-overlapping sets, resulting in the estimate μ^_true_. Across the *m* confidence intervals, we counted the fraction of times μ^_true_ fell within the confidence intervals, which we referred to as the overall coverage rate. A well-calibrated 95% confidence interval will cover *μ*_true_ 95% of the time. The whole procedure was repeated 420 times, once for each scenario.

### Simulation settings

To select an algorithm for each dataset, we used PyCaret (Ali, 2020) to compare algorithms implemented in scikit-learn with default hyperparameters (Pedregosa et al., 2011). We preferred algorithms that performed well on the original data, ensuring room for performance to decrease when noise was introduced in the FPR and power simulations. Computationally intensive algorithms were excluded even if they performed well (e.g., CatBoost).

For EMNIST, we used multiclass logistic regression with accuracy as the test-set evaluation metric. For UK Biobank, we used ridge regression with the coefficient of determination as the metric. For Covertype, we used an Extra Trees classifier with accuracy as the primary metric. For KEGG Metabolic Pathway, we used an Extra Trees regressor with mean squared error as the metric.

Each algorithm was evaluated under three hyperparameter-selection schemes: fixed hyperparameters, selection using an internal 80%/20% training-validation split within the training data, and selection using 2-fold cross-validation within the training data. For the two tuning schemes, the inner splits were identical across candidate hyperparameters, and the selected configuration was refitted on the full training set before evaluation on the test set. Hyperparameter ranges and further implementation details are provided in Supplementary Methods S4.

We evaluated four sample sizes per dataset, except EMNIST and UKB, for which three were considered (see Methods “FPR simulation”), yielding 14 dataset–sample-size combinations. Together with ten noise levels and three hyperparameter schemes, this produced 420 simulation scenarios. Covertype was the only binary-classification dataset and was therefore additionally evaluated using AUC to assess the DeLong test. KEGG was the only dataset for which performance was recorded at the sample level, yielding 120 scenarios for comparing sample-level and fold-averaged tests.

## Existing statistical tests for cross-validation

As explained in Methods “Cross-validation primer”, most tests considered here operate on a vector ***D*** containing *J* corresponding performance differences between two models. We evaluated commonly used independence-based tests, existing correlation-aware tests, and three bootstrap procedures. Supplementary Methods S5 provides more details about the tests.

The naïve paired *t*-test (also referred to as the resampled uncorrected paired *t*-test; Nadeau & Bengio, 2003), Wilcoxon signed-rank test and sign-flip permutation test all treat the fold-averaged differences as independent. The DeLong test, when applied to sample-level statistics generated under cross-validation, likewise assumes independence across samples. We therefore expected these tests to have elevated FPR.

The corrected resampled paired *t*-test modifies the conventional variance estimator to account for positive correlation between fold-averaged differences (Nadeau & Bengio, 2003). However, it assumes that the correlation is equal to the ratio of test-set size to the full sample size, rather than estimating it from the data. The 5×2 paired *t*-test (Dietterich, 1998) and 5×2 paired *F*-test (Alpaydın, 1999) use five repetitions of 2-fold cross-validation. Both account for between-fold correlation through their variance estimators, although the 5×2 *t*-test uses only one of the ten performance differences in its numerator and therefore discards information. The empirical test of differences computes a two-sided p-value as twice the smaller proportion of positive or negative fold-averaged differences.

We also evaluated three bootstrap procedures. Bootstrap-orig repeatedly generates a bootstrap training set from the original samples, evaluates models on the corresponding out-of-bag test set and constructs an interval from the resulting empirical distribution (Raschka, 2018). The other two bootstrap variants instead resample fold-averaged differences obtained through cross-validation. Bootstrap-ET (short for bootstrap empirical test) applies the empirical test of differences to the resampled differences, whereas bootstrap-*t* applies a paired *t*-test. Bootstrap-orig and bootstrap-ET controlled the FPR, whereas bootstrap-*t* had a high FPR because the paired t-test treats the resampled differences as independent.

The FPR, power and CI coverage of the statistical tests are summarized in Fig. 8a. Further theoretical analyses are provided in Supplementary Methods S6**–**S10. SHARP is described in the next section.

## SHARP test

The challenge of inference on performance differences comes down to estimating the standard error of the sample mean difference while accounting for the dependence between the *J* differences. Perhaps the clearest illustration of the problem is to consider a single application of K-fold cross-validation: The *J* entries in the vector ***D*** of fold-level differences are correlated, and for Gaussian data, there are two sufficient statistics (sample mean and sample variance), but three unknown parameters (mean, variance, and correlation). Thus, the model is overparameterized and, specifically, the variance and correlation cannot both be estimated without restrictive assumptions. The same fundamental statistical ambiguity applies to Monte Carlo cross-validation.

In their seminal study, Bengio and Grandvalet (2004) considered a single instance of K-fold cross-validation and worked from sample-level prediction errors instead of fold-averaged performance (see Methods “Cross-validation primer”). They showed that there is no universal, distribution-independent unbiased estimator of cross-validation variance. In particular, for Gaussian data, the maximum likelihood estimator does not exist. The covariance structure in their setting (based on sample-level prediction performance) is equivalent to the covariance structure of fold-averaged prediction performance under repeated K-fold cross-validation, so their findings generalize to repeated K-fold cross-validation.

The key idea behind the SHARP test is to create a situation where independent data are generated that will allow estimation of the variance parameters *σ*^2^ and *ρ*. To achieve this, we randomly divide the dataset into two disjoint halves A and B. For each machine learning model, we then perform K-fold cross-validation in subsets A and B separately. We then average the results across the K-fold cross-validation, resulting in a pair of model performance differences, *D_A_*_1_ and *D_B_*_1_ respectively. Alternatively, one iteration of Monte Carlo cross-validation can be done within each subset A and B, likewise producing a pair of performance *D_A_*_1_ and *D_B_*_1_ based on the held-out testing data in each half.

We note that *D_A_*_1_ and *D_B_*_1_ are independent since subsets A and B are non-overlapping. For the analyses in the current study, this process is repeated *J* times, resulting in two vectors ***D****_A_* = [*D_A_*_1_, …, *D_AJ_*] and ***D****_B_* = [*D_B_*_1_, …, *D_BJ_*]. We note that while *D_Aj_* and *D_Bj_* (from the *j*-th iteration) are independent, the entries within ***D****_A_* (and ***D****_B_*), and *D_Aj_* and *D_Bj_*′ for *j* ≠ *j*^′^ are dependent with a common correlation. Therefore, we have *J* pairs of independent observations and three unknowns: mean performance *μ* the variance *σ*^2^ of each entry in ***D****_A_* and ***D****_B_*, and correlation *ρ* among all non-paired entries in ***D****_A_* and ***D****_B_*. Given ***D****_A_* and ***D****_B_*, we can estimate all three unknowns to perform the statistical test.

We define *D^-^* to be the grand average of the two vectors ***D****_A_* and ***D****_B_*. We seek to test the null hypothesis of equal expected performance:

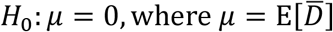

We show that the SHARP estimator of performance difference

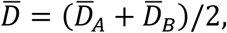

where *D^-^_A_* and *D^-^_B_* are the respective sample means, is the generalized least-squares estimator of *μ* (Supplementary Methods S12.2), with variance

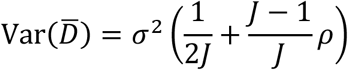

and so inference reduces to estimating *σ^2^* and *ρ*. To estimate *σ^2^* and *ρ*, we considered method-of-moments (Supplementary Methods S12.3), maximum likelihood (Supplementary Methods S12.4) or restricted maximum likelihood (Supplementary Methods S12.5), followed by a Wald test. We also considered likelihood-based tests including score test (Supplementary Methods S12.6) and likelihood ratio test (Supplementary Methods S12.7).

Based on simulated data (Supplementary Methods S12.8), we found that the score test provided the best control of FPR (Supplementary Fig. 12). Therefore, all results in the current study utilized the score test. For the score test, the Z-score is a ratio of *D^-^* to a standard error computed using Var(*D^-^*) above, but with estimates μ^^2^ and ρ^_0_ obtained by optimizing the Gaussian likelihood assuming the null hypothesis is true, i.e., *μ* = 0.

More details about the SHARP test can be found in Supplementary Methods S12. For the purpose of evaluating FPR and statistical power, we considered two settings: (1) *K* = 5, *J* = 60 and (2) *K* = 10, *J* = 30. Here, J was chosen to ensure convergence of the SHARP test.

## Ethics and data availability

Use of de-identified data from UKB datasets is approved by the National University of Singapore (NUS) Institutional Review Board (IRB). Sex and gender information was not used in this study, and race and ethnicity information was not used in this study.

This study used publicly available data from the UK Biobank (https://www.ukbiobank.ac.uk/), EMNIST (https://www.kaggle.com/datasets/crawford/emnist), Covertype (https://archive.ics.uci.edu/dataset/31/covertype) and KEGG Metabolic Pathway (https://archive.ics.uci.edu/dataset/220/kegg+metabolic+relation+network+directed).

## Code availability

Code for this study can be found here (GITHUB_LINK). Co-authors (TZ and HL) reviewed each other’s code before merging into the GitHub repository to reduce the chance of coding errors.

## Author Contributions

Conceptualization, T.Z., H.L., S.Z., Y.Q.T., C.O., K.K., D.B., T.E.N., and B.T.T.Y.; Methodology, T.Z., H.L., S.Z., C.O., O.R., T.E.N., and B.T.T.Y.; Software, T.Z., H.L., S.Z., Y.Q.T., and F.T.; Validation, T.Z., H.L., S.Z., K.P., and O.R.; Formal analysis, T.Z., H.L., and S.Z.; Investigation, T.Z., H.L., S.Z., Y.Q.T., F.T., L.A., W.C., J.C., J.S.X.C., N.D., Z.D., X.L., Z.Li, M.J.R.L., Y.L., Q.L., Z.Ling, X.Z.L., S.M.L., E.K.K.N., T.T.N., L.Q.R.O., S.P., X.Q., J.R., Z.W., Y.X., C.Z., and Y.Z.; Resources: J.H.Z., and B.T.T.Y; Data curation, T.Z., H.L., S.Z., and F.T.; Visualization, T.Z., H.L., and S.Z.; Writing – original draft, T.Z., H.L., S.Z., T.E.N., and B.T.T.Y.; Writing – review & editing, all authors; Supervision, C.O., N.U.F.D., T.E.N., and B.T.T.Y.; Project administration, B.T.T.Y.; Funding acquisition, K.P., L.P, S.C., S.E., and B.T.T.Y.

## Declaration of generative AI and AI-assisted technologies in the writing process

The authors used Claude (Anthropic; Opus 4.8 and Opus 5) and ChatGPT (OpenAI; GPT-5.5, GPT-5.6, GPT-6) to improve the readability and language of the manuscript, including checking and revising author-drafted text. Throughout this iterative process, the authors reviewed and edited the content as needed. The authors take full responsibility for the content of the published article.

## Acknowledgements

Our research is supported by the NUS Yong Loo Lin School of Medicine (NUHSRO/2020/124/TMR/LOA), the Singapore National Medical Research Council (NMRC) LCG (OFLCG19May-0035), NMRC CTG-IIT (CTGIIT23jan-0001), NMRC OF-IRG (OFIRG24jan-0006; OFIRG24jul-0049), NMRC STaR (STaR20nov-0003), Singapore Ministry of Health (MOH) Centre Grant (CG21APR1009), the United States National Institutes of Health (R01MH133334 & 2R01MH120080) and the Singapore National Research Foundation (NRF) Investigatorship (NRFI10-2024-0014). This research has been conducted using the UK Biobank Resource under application number 25163. S.C. is supported by the University of Melbourne McKenzie Fellowship and a Brain Behaviour Research Foundation - Young Investigator Grant. L.P. is supported by NIH (R00MH127296). This project/research has received funding from the European Union’s Horizon Europe Programme under the Specific Grant Agreement No. 101147319 (EBRAINS 2.0 Project) and A platform for behavioral markers of digital neuro-medicine in NRW (KP22-106A). Any opinions, findings and conclusions or recommendations expressed in this material are those of the authors and do not reflect the views of the funders.

## References

Alfaro-Almagro, F., Jenkinson, M., Bangerter, N. K., Andersson, J. L. R., Griffanti, L., Douaud, G., Sotiropoulos, S. N., Jbabdi, S., Hernandez-Fernandez, M., Vallée, E., Vidaurre, D., Webster, M., McCarthy, P., Rorden, C., Daducci, A., Alexander, D. C., Zhang, H., Dragonu, I., Matthews, P. M., … Smith, S. M. (2018). Image processing and quality control for the first 10,000 brain imaging datasets from UK Biobank. NeuroImage, 166, 400–424. 10.1016/j.neuroimage.2017.10.034

Ali, M. (2020). PyCaret: An open source, low-code machine learning library in Python (Version 1.0) [Computer software]. https://www.pycaret.org

Alpaydın, E. (1999). Combined 5 × 2 cv F test for comparing supervised classification learning algorithms. Neural Computation, 11(8), 1885–1892. 10.1162/089976699300016007

André, P., Heitz, C., Christodoulou, E., Reinke, A., Sudre, C. H., Antonelli, M., Godau, P., Cardoso, M. J., Gilson, A., Tezenas du Montcel, S., Varoquaux, G., Maier-Hein, L., & Colliot, O. (2026). *Performance uncertainty in medical image analysis: A large-scale investigation of confidence intervals* (arXiv:2601.17103). arXiv. 10.48550/arXiv.2601.17103

Bates, S., Hastie, T., & Tibshirani, R. (2024). Cross-validation: What does it estimate and how well does it do it? Journal of the American Statistical Association, 119(546), 1434–1445. 10.1080/01621459.2023.2197686

Bengio, Y., & Grandvalet, Y. (2004). No unbiased estimator of the variance of K-fold cross-validation. Journal of Machine Learning Research, 5, 1089–1105.

Blackard, J. (1998). *Covertype* [Data set]. UCI Machine Learning Repository. 10.24432/C50K5N

Bouckaert, R. R., & Frank, E. (2004). Evaluating the replicability of significance tests for comparing learning algorithms. In H. Dai, R. Srikant, & C. Zhang (Eds.), Advances in knowledge discovery and data mining (Lecture Notes in Computer Science Vol. 3056, pp. 3–12). Springer. 10.1007/978-3-540-24775-3_3

Bzdok, D., Thieme, A., Levkovskyy, O., Wren, P., Ray, T., & Reddy, S. (2024). Data science opportunities of large language models for neuroscience and biomedicine. Neuron, 112(5), 698–717. 10.1016/j.neuron.2024.01.016

Carrasco-Zanini, J., Pietzner, M., Davitte, J., Surendran, P., Croteau-Chonka, D. C., Robins, C., Torralbo, A., Tomlinson, C., Grünschläger, F., Fitzpatrick, N., Ytsma, C., Kanno, T., Gade, S., Freitag, D., Ziebell, F., Haas, S., Denaxas, S., Betts, J. C., Wareham, N. J., … Langenberg, C. (2024). Proteomic signatures improve risk prediction for common and rare diseases. Nature Medicine, 30(9), 2489–2498. 10.1038/s41591-024-03142-z

Chen, J., Tam, A., Kebets, V., Orban, C., Ooi, L. Q. R., Asplund, C. L., Marek, S., Dosenbach, N. U. F., Eickhoff, S. B., Bzdok, D., Holmes, A. J., & Yeo, B. T. T. (2022). Shared and unique brain network features predict cognitive, personality, and mental health scores in the ABCD study. Nature Communications, 13(1), 2217. 10.1038/s41467-022-29766-8

Chopra, S., Dhamala, E., Lawhead, C., Ricard, J. A., Orchard, E. R., An, L., Chen, P., Wulan, N., Kumar, P., Rubenstein, A., Moses, J., Chen, L., Levi, P., Holmes, A., Aquino, K., Fornito, A., Harpaz-Rotem, I., Germine, L. T., Baker, J. T., … Holmes, A. J. (2024). Generalizable and replicable brain-based predictions of cognitive functioning across common psychiatric illness. Science Advances, 10(45), eadn1862. 10.1126/sciadv.adn1862

Cohen, G., Afshar, S., Tapson, J., & van Schaik, A. (2017). EMNIST: Extending MNIST to handwritten letters. 2017 International Joint Conference on Neural Networks (IJCNN), 2921–2926. 10.1109/IJCNN.2017.7966217

Collins, G. S., Dhiman, P., Ma, J., Schlussel, M. M., Archer, L., Van Calster, B., Harrell, F. E., Jr., Martin, G. P., Moons, K. G. M., van Smeden, M., Sperrin, M., Bullock, G. S., & Riley, R. D. (2024). Evaluation of clinical prediction models (part 1): From development to external validation. BMJ, 384, e074819. 10.1136/bmj-2023-074819

DeLong, E. R., DeLong, D. M., & Clarke-Pearson, D. L. (1988). Comparing the areas under two or more correlated receiver operating characteristic curves: A nonparametric approach. Biometrics, 44(3), 837–845. 10.2307/2531595

Demšar, J. (2006). Statistical comparisons of classifiers over multiple data sets. Journal of Machine Learning Research, 7, 1–30.

Dietterich, T. G. (1998). Approximate statistical tests for comparing supervised classification learning algorithms. Neural Computation, 10(7), 1895–1923. 10.1162/089976698300017197

Edgington, E. S., & Onghena, P. (2007). *Randomization tests* (4th ed.). Chapman and Hall/CRC. 10.1201/9781420011814

Geras, K. J., & Sutton, C. (2013). Multiple-source cross-validation. Proceedings of the 30th International Conference on Machine Learning, 28(3), 1292–1300. https://proceedings.mlr.press/v28/geras13.html

Greene, A. S., Gao, S., Scheinost, D., & Constable, R. T. (2018). Task-induced brain state manipulation improves prediction of individual traits. Nature Communications, 9(1), 2807. 10.1038/s41467-018-04920-3

Jafrasteh, B., Adeli, E., Pohl, K. M., Kuceyeski, A., Sabuncu, M. R., & Zhao, Q. (2025). Statistical variability in comparing accuracy of neuroimaging based classification models via cross validation. Scientific Reports, 15(1), 28745. 10.1038/s41598-025-12026-2

Justice, A. C., Covinsky, K. E., & Berlin, J. A. (1999). Assessing the generalizability of prognostic information. Annals of Internal Medicine, 130(6), 515–524. 10.7326/0003-4819-130-6-199903160-00016

la Roi-Teeuw, H. M., van Royen, F. S., de Hond, A., Zahra, A., de Vries, S., Bartels, R., Carriero, A. J., van Doorn, S., Dunias, Z. S., Kant, I., Leeuwenberg, T., Peters, R., Veerhoek, L., van Smeden, M., & Luijken, K. (2024). Don’t be misled: 3 misconceptions about external validation of clinical prediction models. Journal of Clinical Epidemiology, 172, 111387. 10.1016/j.jclinepi.2024.111387

Mann, H. B., & Whitney, D. R. (1947). On a test of whether one of two random variables is stochastically larger than the other. The Annals of Mathematical Statistics, 18(1), 50– 60. 10.1214/aoms/1177730491

Mansour L., S., Tian, Y., Yeo, B. T. T., Cropley, V., & Zalesky, A. (2021). High-resolution connectomic fingerprints: Mapping neural identity and behavior. NeuroImage, 229, 117695. 10.1016/j.neuroimage.2020.117695

Moor, M., Banerjee, O., Abad, Z. S. H., Krumholz, H. M., Leskovec, J., Topol, E. J., & Rajpurkar, P. (2023). Foundation models for generalist medical artificial intelligence. Nature, 616(7956), 259–265. 10.1038/s41586-023-05881-4

Nadeau, C., & Bengio, Y. (2003). Inference for the generalization error. Machine Learning, 52(3), 239–281. 10.1023/A:1024068626366

Naeem, M., & Asghar, S. (2011). *KEGG Metabolic Relation Network (Directed)* [Data set]. UCI Machine Learning Repository. 10.24432/C5CK52

Newcombe, R. G. (1998). Two-sided confidence intervals for the single proportion: Comparison of seven methods. Statistics in Medicine, 17(8), 857–872. 10.1002/(SICI)1097-0258(19980430)17:8%3C857::AID-SIM777%3E3.0.CO;2-E

Open Science Collaboration. (2015). Estimating the reproducibility of psychological science. Science, 349(6251), aac4716. 10.1126/science.aac4716

Page, M. J., McKenzie, J. E., Bossuyt, P. M., Boutron, I., Hoffmann, T. C., Mulrow, C. D., Shamseer, L., Tetzlaff, J. M., Akl, E. A., Brennan, S. E., Chou, R., Glanville, J., Grimshaw, J. M., Hróbjartsson, A., Lalu, M. M., Li, T., Loder, E. W., Mayo-Wilson, E., McDonald, S., … Moher, D. (2021). The PRISMA 2020 statement: An updated guideline for reporting systematic reviews. BMJ, 372, n71. 10.1136/bmj.n71

Parkes, L., Moore, T. M., Calkins, M. E., Cieslak, M., Roalf, D. R., Wolf, D. H., Gur, R. C., Gur, R. E., Satterthwaite, T. D., & Bassett, D. S. (2021). Network controllability in transmodal cortex predicts positive psychosis spectrum symptoms. Biological Psychiatry, 90(6), 409–418. 10.1016/j.biopsych.2021.03.016

Pedregosa, F., Varoquaux, G., Gramfort, A., Michel, V., Thirion, B., Grisel, O., Blondel, M., Prettenhofer, P., Weiss, R., Dubourg, V., Vanderplas, J., Passos, A., Cournapeau, D., Brucher, M., Perrot, M., & Duchesnay, É. (2011). Scikit-learn: Machine learning in Python. Journal of Machine Learning Research, 12, 2825–2830.

Perez-Lopez, R., Ghaffari Laleh, N., Mahmood, F., & Kather, J. N. (2024). A guide to artificial intelligence for cancer researchers. Nature Reviews Cancer, 24(6), 427–441. 10.1038/s41568-024-00694-7

Rajpurkar, P., Chen, E., Banerjee, O., & Topol, E. J. (2022). AI in health and medicine. Nature Medicine, 28(1), 31–38. 10.1038/s41591-021-01614-0

Raschka, S. (2018). *Model evaluation, model selection, and algorithm selection in machine learning* (arXiv:1811.12808). arXiv. 10.48550/arXiv.1811.12808

Riley, R. D., Ensor, J., Snell, K. I. E., Debray, T. P. A., Altman, D. G., Moons, K. G. M., & Collins, G. S. (2016). External validation of clinical prediction models using big datasets from e-health records or IPD meta-analysis: Opportunities and challenges. BMJ, 353, i3140. 10.1136/bmj.i3140

Rosenblatt, M., Foster, M. L., Adkinson, B. D., Tejavibulya, L., Khaitova, M., Ye, J., Sun, H., Rodriguez, R. X., Camp, C. C., Chinta, A., McCusker, M. C., Han, L., Fields, C. T., Mehta, S., & Scheinost, D. (2026). External validation improves generalizability, replicability and reproducibility in predictive models for neuroimaging. Nature Methods. Advance online publication. 10.1038/s41592-026-03115-9

Steyerberg, E. W., & Harrell, F. E., Jr. (2016). Prediction models need appropriate internal, internal–external, and external validation. Journal of Clinical Epidemiology, 69, 245– 247. 10.1016/j.jclinepi.2015.04.005

Takada, T., Nijman, S., Denaxas, S., Snell, K. I. E., Uijl, A., Nguyen, T.-L., Asselbergs, F. W., & Debray, T. P. A. (2021). Internal-external cross-validation helped to evaluate the generalizability of prediction models in large clustered datasets. Journal of Clinical Epidemiology, 137, 83–91. 10.1016/j.jclinepi.2021.03.025

Varoquaux, G., Raamana, P. R., Engemann, D. A., Hoyos-Idrobo, A., Schwartz, Y., & Thirion, B. (2017). Assessing and tuning brain decoders: Cross-validation, caveats, and guidelines. NeuroImage, 145, 166–179. 10.1016/j.neuroimage.2016.10.038

Wagner, S. J., Reisenbüchler, D., West, N. P., Niehues, J. M., Zhu, J., Foersch, S., Veldhuizen, G. P., Quirke, P., Grabsch, H. I., van den Brandt, P. A., Hutchins, G. G. A., Richman, S. D., Yuan, T., Langer, R., Jenniskens, J. C. A., Offermans, K., Mueller, W., Gray, R., Gruber, S. B., … Kather, J. N. (2023). Transformer-based biomarker prediction from colorectal cancer histology: A large-scale multicentric study. Cancer Cell, 41(9), 1650–1661.e4. 10.1016/j.ccell.2023.08.002

Wilcoxon, F. (1945). Individual comparisons by ranking methods. Biometrics Bulletin, 1(6), 80–83. 10.2307/3001968

Yeo, B. T. T., Sabuncu, M. R., Vercauteren, T., Holt, D. J., Amunts, K., Zilles, K., Golland, P., & Fischl, B. (2010). Learning task-optimal registration cost functions for localizing cytoarchitecture and function in the cerebral cortex. IEEE Transactions on Medical Imaging, 29(7), 1424–1441. 10.1109/TMI.2010.2049497

Yoo, S.-K., Fitzgerald, C. W., Cho, B. A., Fitzgerald, B. G., Han, C., Koh, E. S., Pandey, A., Sfreddo, H., Crowley, F., Korostin, M. R., Debnath, N., Leyfman, Y., Valero, C., Lee, M., Vos, J. L., Lee, A. S., Zhao, K., Lam, S., Olumuyide, E., … Chowell, D. (2025). Prediction of checkpoint inhibitor immunotherapy efficacy for cancer using routine blood tests and clinical data. Nature Medicine, 31(3), 869–880. 10.1038/s41591-024-03398-5

