## Supplementary Material for "Spurious model comparisons are widespread in biomedical artificial intelligence"

#### Supplemental Material

This supplemental material consists of Supplemental Methods, Results, Tables and Figures to complement the main text.

#### Supplemental Methods

#### S1. Large Language Model (LLM) settings

##### S1.1 LLM screening and manual verification

As mentioned in the main text, we used an LLM (Claude Opus 4.1, Anthropic) to scale screening and data extraction. The LLM prompts can be found in Supplementary Methods S1.2 to S1.4. Here we provide details about the application of different manual verification procedures to the five LLM questions.

1. **Scientific field classification (Q1).** If a study was published in a multi-field journal, the LLM was instructed to assign the study to a single scientific subfield using a predefined set of categories, or "OTHERS" if none applied.

   *Manual verification*: In a random audit of 50 studies, two human raters (TZ and HL) independently assessed each study. Any discrepancy between raters was resolved by discussion and consensus. The LLM agreed with the consensus human classification for all 50 studies. Therefore, we used the LLM results directly.

2. **Model comparison evaluation (Q2).** The LLM was asked to determine whether the study satisfied PRISMA criterion 1 (Methods " PRISMA workflow and criteria").

   *Manual verification*: In a random audit of 50 studies, two human raters (TZ and HL) independently assessed each study. Any discrepancy between raters was resolved by discussion and consensus. The LLM agreed with the consensus human classification for all 50 studies. Therefore, we used the LLM results directly.

3. **Statistical inference (Q3).** If the LLM's answer to Q2 was "Yes", the LLM was instructed to identify every instance in which a p-value or confidence interval was reported to compare model performance. Studies with at least one instance were deemed eligible under PRISMA criterion 2 (Methods "PRISMA workflow and criteria").

   *Manual verification:* In a random audit of 50 studies, two human raters (TZ and HL) independently assessed each study. Any discrepancy between raters was resolved by discussion and consensus. The LLM agreed with consensus human classification for 39 studies (78%). For the remaining 11 studies, there was only one false negative, i.e., LLM counted a study as being ineligible under PRISMA criterion 2, disagreeing with the human consensus. We considered this level of false negatives to be acceptable. The remaining 10 errors were false positives, i.e., the LLM counted the studies as being eligible under PRISMA criterion 2, disagreeing with the human consensus. Given the relatively high false positive rate, human raters read the full text (including reporting summary) of all 1197 studies that the LLM considered to satisfy PRISMA criterion 2.

More specifically, each study was independently assessed by two human raters (TZ, HL or SZ), and discrepancies were resolved by discussion and consensus.

4. **Statistical test details (Q4).** For each instance identified in Q3, the LLM was instructed to extract the name of the test or confidence interval, details about compared models, performance metric(s) used for the comparison, and the cross-validation procedure used. The LLM was asked to support its conclusions with direct quotations from the study.

   *Manual verification*: The extracted quotations served only as references: two human raters (TZ, HL or SZ) independently re-derived this information for every study. Discrepancies were resolved by discussion and consensus.

5. **Data classification (Q5).** For each instance identified in Q3, the LLM was also asked to classify the values entering the test as either derived from cross-validation folds (referred to as "resampling units" in our prompt) or as "Other/Unclear", corresponding to PRISMA criterion 3 (Methods "PRISMA workflow and criteria").

   *Manual verification:* The LLM classification served as a reference: two human raters (TZ, HL or SZ) independently examined every study to determine whether PRISMA criterion 3 was satisfied. Discrepancies were resolved by discussion and consensus.

The procedure above yielded an initial set of 210 studies that satisfied all four PRISMA criteria. Before finalizing the corpus, we re-screened these 210 studies using newer LLMs (Claude Opus 4.8 and Opus 5, Anthropic) with access to the full text, as well as all available code, data and supplementary files. Based on this information, two raters (TZ and HL) independently re-assessed each study, and discrepancies were again resolved by discussion and consensus. The Q4 and Q5 information was updated for some studies during this re-screening. This additional review further excluded 26 studies, which are included among the 628 exclusions under PRISMA criterion 3 (in Fig. 2a), leaving the final set of 184 studies.

For each of the 184 included studies, the human-verified information from Q4 and Q5 was used to classify the study into one of two groups: (i) at least one statistical test ignored between-fold correlation, or (ii) every statistical test accounted for between-fold correlation.

#### S1.2 Prompt for screening papers

We developed an automated pipeline for systematically evaluating academic papers using a large language model (LLM) to identify and classify methodological characteristics related to statistical inference in predictive modeling studies. The implementation leverages the Claude API (claude-opus-4-1-20250805) to perform structured content analysis on full-text scientific articles. We set LLM temperature at 0 for deterministic outputs. No system-level prompt was employed; all instructions were contained within the user message. The maximum output token limit was set to 15,000. The full prompt is shown below:

You are tasked with evaluating a scientific paper by answering questions Q1 through Q5. Follow these instructions carefully and adhere to the specified output formats.

First, read the full text of the paper:

<paper_text>
{{PAPER_TEXT}}
</paper_text>

You will also use the category list below for Q1, the categories are separated by ";"

<category_list>
{{CATEGORY_LIST}}
</category_list>

Now, answer the following questions in order:

**Q1: Paper Classification**
Classify the paper into one of the categories from the category list based on its main content and purpose. Select the most appropriate category that best represents the paper's main goal, method, or contribution. If no category is clearly applicable, answer "OTHERS".

Output your answer in this format:
<answer>
Q1: [The most appropriate category/OTHERS]
Evidence_or_Reason: [One to three sentences justification]
</answer>

**Q2: Method Comparison Evaluation**
Evaluate whether the paper compares at least two distinct alternatives, where each alternative can represent a trainable method or pipeline that undergoes model training and evaluation on data.

Alternatives can differ in algorithms or architectures, hyperparameter configurations, training strategies, feature sets/biomarkers, data modalities/preprocessing pipelines, or can be model selection procedures applied to the development dataset. The comparison must be supported by reported metrics computed directly from predictions against ground truth (e.g., accuracy, AUC, MSE, Pearson correlation).

Answer "Yes" if the condition is met. Answer "No" if the condition is not met.
If your answer is "Yes", provide one to five verbatim sentences from the paper that directly support this.
If your answer is "No", provide a short reasoning explaining why the condition is not met.

Output your answer in this format:
<answer>
Q2: [Yes/No]
Evidence_or_Reason: [One to five sentences justification or verbatim quotes]
</answer>

If you answered "Yes" to Q2, create an internal scratchpad (do not output it) containing all verbatim sentences or short passages in the paper that report quantitative comparisons between methods. You will use this scratchpad for the remaining questions.

**Q3: Statistical Inference Count**

Only evaluate Q3 if Q2 = "Yes". Otherwise, answer "N/A".

Re-read your internal scratchpad from Q2 and the full paper. Identify every instance where the paper performs a statistical inference in a quantitative comparison by reporting either a p-value or a confidence interval. Include cases where p-values or confidence intervals are reported during model selection in the development dataset.

Count the total number of statistical test instances found.

Output your answer in this format:
<answer>
Q3: [Integer/N/A]
</answer>

If your answer to Q3 is greater than 0, create a new internal scratchpad (do not output it) to include the verbatim sentences documenting each statistical test and confidence interval, prefixed with index numbers (1, 2, 3, …).

**Q4: Statistical Test Details**
Only evaluate Q4 if Q3 returns a number greater than zero. Otherwise, answer "N/A".

For each indexed passage in your internal scratchpad created in Q3 that documents a statistical test or confidence interval, provide:
1. Test name: the exact name of the statistical test or interval the paper used (e.g., the DeLong test, bootstrap confidence interval, paired t-test, 95% CI)
2. Data description:
   a. What two alternatives are compared
   b. What prediction performance metric(s) entered the test or confidence interval (e.g., AUCs, accuracies, MSE values)
   c. How those performance data were obtained from model training and evaluation (train/test splits, cross-validation folds, bootstrap or permutation resampling, etc.)

Answer in a numbered list corresponding to each indexed statistical test passage from Q3.

Output your answer in this format, you don't need to include the evidence in your answer for Q4:
<answer>
Q4:
1. [Test name]
   a: [alternatives compared]
   b: [performance metrics used]
   c: [how performance data were obtained]

2. [Test name]
   a: [alternatives compared]
   b: [performance metrics used]
   c: [how performance data were obtained]
...
</answer>

After answering Q4, update your internal scratchpad by adding to each indexed test the verbatim sentences that support: (1) the test name, and (2) the description of the performance data used and how those data were obtained.

**Q5: Data Classification**
Only evaluate Q5 if Q3 returns a number greater than zero. Otherwise, answer "N/A".

For each indexed statistical test, classify the values that entered the test or built the confidence intervals into one of:
a. Cross-validation folds - every value is derived from a "resampling unit", which may correspond to: (i) one of the K partitions in K-fold cross-validation, (ii) the test set from one of the K repeated random train-test splits, (iii) one of the N averages obtained from N repetitions of K-fold cross-validation
b. Other / Unclear

Follow these rules:
1. Cross-validation includes K-fold CV, Monte-Carlo CV, Leave-one-out CV, and leave-one-dataset-out CV
2. Apply categories in order: if (a) is satisfied, do not consider (b)
3. Always provide your reasoning after your classification

Output your answer in this format:
<answer>
Q5 Answer:
1. [a/b]
   Reason: [your justification]
2. [a/b]
   Reason: [your justification]
...
</answer>

If your answer is "a" for any test, add to that indexed test in your scratchpad the verbatim sentences focusing on how the cross-validations are carried out for this test.

**Final Output**
After completing all questions, output your final scratchpad:

<final_scratchpad>
[Your complete scratchpad content with all indexed statistical tests and supporting verbatim sentences]
</final_scratchpad>

**General Rules:**
1. For Q4, always quote verbatim sentences describing the data that entered the test, not just the test name
2. Preserve the exact wording from the paper for all quoted text
3. Do not include any information that cannot be traced back directly to the paper text
4. Follow the conditional logic: Q3-Q5 depend on your answers to previous questions
5. Use the exact output formats specified for each question

#### S1.3 Prompt for evaluating journal rigor

We implemented an automated policy-auditing pipeline using the OpenAI API (gpt-5-2025-08-07) with integrated web-search capability. The model was configured with high reasoning effort, a temperature of 0.2, and a maximum output token limit of 20,000 to prevent truncation. The built-in web_search tool was enabled to allow retrieval of authoritative first-party journal webpages. All task instructions and scoring criteria were provided within the user prompt. The full prompt is shown below:

You are auditing journal policies for: {journal}.

Task:
1) For each criterion below, find the most authoritative public policy page(s) (Instructions for Authors, Editorial Policies, Submission Checklist, Reporting Guidelines).
2) Give exact quotes from webpages of the relevant policy concisely (maximum 3 sentences). If the criterion is not mentioned, say so.
3) Assign a score (0=not mentioned/none, 1=encouraged/optional, 2=required/mandatory).
4) Provide the 1 or 2 best source URLs as evidence for each criterion (prefer first-party journal/publisher pages).

Criteria:
{json.dumps(CRITERIA, ensure_ascii=False, indent=2)}

Description of criteria:
- Code availability policy: Whether the journal encourages or requires authors to share analysis code or model scripts.
- Data availability policy: Whether the journal encourages or requires sharing of underlying datasets or processed data.
- Availability statement requirement: Whether a formal data or code availability statement is encouraged or required in the manuscript.
- Code required for peer review: Whether the journal encourages or requires authors to provide the custom code or scripts used for analysis to editors and reviewers during peer review.
- Statistical test review policy: Whether review of statistical tests is an optional or mandatory part of the editorial or peer review process.
- Reporting standards or checklist: Whether the journal encourages or mandates adherence to recognized methodological reporting standards (e.g., TRIPOD-AI, CONSORT-AI, PRISMA).

Return STRICT JSON matching the provided schema.
Scoring must be conservative; if unclear, give 1 (encouraged) not 2.

Schema:
{json.dumps(schema, ensure_ascii=False)}

#### S1.4 Prompt for extracting study-level code and data availability

We implemented an automated pipeline using the Anthropic API (claude-opus-4-1-20250805) to classify whether each study reported data and/or code availability. The model was configured with a temperature of 0 to ensure deterministic outputs and a maximum output token limit of 15,000 to prevent truncation. No system-level prompt was used; all instructions were contained within the user message. The full prompt is shown below:

```
You are an expert research auditor evaluating whether a scientific paper shares its data or
code publicly.
Below is the full text of a paper (as scraped from PubMed Central).
Please carefully read it and rate the paper according to the criteria described below.

---

<paper text begins here>
{{PAPER_TEXT}}
<paper text ends here>

---

### Your task:
For each criterion, assign a numerical score, provide a one-sentence justification.
If the score is 1, include a short direct quote (1–2 sentences or a few phrases) from the paper
text that supports your judgment.

---

#### Scoring guidelines

**1. Data availability (0-1):**
   - 0 = No mention of data availability, or data not publicly available.
   - 1 = Data partially available or available upon request or publicly available in a repository
(e.g., Zenodo, Dryad, GEO, OSF) with a clear link or accession number.

**2. Code availability (0-1):**
   - 0 = No mention of code availability or code not available.
   - 1 = Code partially available or available upon request or code is publicly available in a
repository (e.g., GitHub, Zenodo, OSF) with a clear link.

---

### Output format (JSON only):

{
  "data_availability": {
    "score": <0-1>,
    "justification": "<one-sentence summary of reasoning>",
    "quote": "<short direct quote from paper text, empty string if score=0>",
  },
  "code_availability": {
    "score": <0-1>,
    "justification": "<one-sentence summary of reasoning>",
    "quote": "<short direct quote from paper text, empty string if score=0>",
```

```
  }
}
```

#### S2. Re-analysis of published results

S2.1 Assignment of comparisons to recovery tiers & inclusion of studies in the re-analysis

We assigned each originally significant model comparison in 173 studies to one of four recovery tiers (A, A′, B or C) based on whether missing information necessitated additional assumptions, and whether a valid re-analysis is feasible under the study's analytical setup. For Tier A comparisons, predictions for each test sample across all cross-validation splits (together with ground-truth labels) or fold-averaged performance values were available, allowing us to apply a valid test directly. For Tier A′ comparisons, predictions and fold-averaged performance values were unavailable, but sufficient information was reported to recover the naïve test statistic (from a paired t-test or paired z-test). We could therefore apply the correction for between-fold correlation without additional assumptions arising from missing information. A study could contribute comparisons from more than one tier. We re-analyzed all studies that contributed at least one Tier A or Tier A′ comparison (N = 68), as reported in the main text.

Throughout, an originally significant comparison was classified as spurious if re-analysis yielded $p \geq 0.05$ and as retaining significance if it yielded $p < 0.05$. We generally did not adjust for multiple comparisons. In the few cases where the original study had applied a multiple-comparison correction, we retained it only when we could reproduce the exact procedure. Omitting such corrections makes comparisons more likely to retain significance and therefore renders our estimated prevalence of spurious comparisons conservative.

A study was classified as containing a spurious comparison if any Tier A or Tier A′ comparison was spurious. Otherwise, it was classified as containing no spurious comparisons. This study-level classification did not imply that any Tier B or Tier C comparisons in the study would necessarily have retained significance.

Tier B comparisons could be re-analyzed only under additional assumptions arising from missing information. For example, some studies reported results from a Wilcoxon signed-rank test but shared neither predictions nor fold-averaged performance values. No standard correlation-aware correction exists for the Wilcoxon signed-rank test when applied to cross-validation results. Re-analysis would therefore require converting the reported Wilcoxon result to a $z$-statistic and assuming that the correlation-aware correction used for paired $z$-tests in Tier A′ remained valid for this transformed statistic.

Tier C comparisons could not be re-analyzed, either because the necessary information could not be recovered even under reasonable assumptions or because the study's cross-validation and testing setup did not permit a valid re-analysis. For example, in some studies, the cross-validation procedure was described too unclearly or was too irregular to calculate the Nadeau–Bengio correction factor (see Supplementary Methods S2.2 and S2.3). Tier C also included comparisons evaluated using sample-level tests (e.g., the DeLong test) for which the studies did not provide the predictions and ground-truth labels needed to reconstruct fold-averaged performance values and conduct a valid re-analysis.

S2.2 Re-analysis of Tier A comparisons

For Tier A comparisons based on a single dataset, we used either sample-level predictions (and corresponding ground-truth labels) or fold-averaged performance values shared by the study. Because no valid sample-level test exists for cross-validation results, we extracted or computed fold-averaged performance values regardless of whether the original study used sample-level or fold-averaged statistics, and applied the corrected resampled paired t-test (Supplementary Methods S6).

Some Tier comparisons were based on cross-validation performed separately on multiple independent datasets. The original studies concatenated the fold-averaged performance values across datasets and applied a paired test to the concatenated values. This approach failed to account for between-fold correlation within each dataset. We instead averaged the performance differences across folds within each dataset, yielding a single mean performance difference for each dataset. Because the datasets were independent, we could apply standard tests across these dataset-level differences.

For multi-dataset comparisons involving fewer than six independent datasets, we applied the paired t-test because a two-sided Wilcoxon signed-rank test requires at least six observations to attain significance ($p < 0.05$).

For the remaining multi-dataset comparisons, we applied both a paired t-test and a Wilcoxon signed-rank test across datasets. A comparison was classified as spurious only if both tests gave $p \geq 0.05$. If either test was significant ($p < 0.05$), we classified the comparison as retaining significance. This rule was deliberately lenient. These studies had only a small number of independent datasets, a setting in which both the paired t-test and the Wilcoxon signed-rank test have low power. Taking the smaller of the two p-values reduced the risk of classifying a comparison as spurious solely because of low power.

An alternative for these multi-dataset comparisons would be to apply a corrected resampled t-test separately within each dataset (Fig. 8b). Although the dataset-specific results could be assessed for qualitative consistency, they might yield significance in some datasets but not others and therefore would not by themselves provide the single, reproducible significance verdict required for our meta-analysis. We therefore treated datasets as independent units and directly tested the dataset-level mean performance differences.

Some comparisons provisionally assigned to Tier A had been evaluated using a two-sample (unpaired) t-test, treating the two sets of fold-averaged performance values as independent, equal-sized samples. This procedure assumed fold independence and, where applicable, ignored pairing between corresponding folds. In some studies, the folds did not correspond because the authors had not fixed the random seed when performing cross-validation. In others, the research question required a cross-validation design without corresponding folds. For others, we could not determine whether the original test had been paired or unpaired.

The Nadeau-Bengio correction cannot be applied to the two-sample case. Accordingly, for comparisons evaluated using a two-sample test, or for which the test type was unclear, we carefully examined the study and its code to determine whether the folds corresponded. If they did, we applied the corrected resampled paired *t*-test. If the folds did not correspond or we could not determine whether they did, we re-assigned the comparisons to Tier B.

Some Tier A comparisons were explicitly evaluated using one-sided tests. For these comparisons, we applied the corresponding valid one-sided test: the corrected resampled paired *t*-test for comparisons based on one dataset, or both the paired *t*-test and the Wilcoxon signed-rank test for comparisons based on multiple datasets. We did not retrospectively assess whether the one-sided test was scientifically justified, but retained the directionality specified by the original studies.

For some Tier A comparisons, we could not determine whether the original test was one-sided or two-sided. We therefore performed both one- and two-sided corrections. If both corrected p-values were $\geq 0.05$, the comparison was classified as spurious. If both p-values were $< 0.05$, the comparison was classified as retaining significance. If the two corrected p-values yielded different significance verdict, the comparison was re-assigned to Tier B, because determining whether to apply a one- or two-sided tests would require an additional assumption.

Finally, we reassigned to Tier B any comparisons for which we could not reproduce the reported *p* value, because retaining them in Tier A would require assuming that the discrepancy arose from an error in the reported *p* value rather than in the shared data.

##### S2.3 Re-analysis of Tier A′ comparisons

For Tier A′ comparisons, prediction data and fold-averaged performance values were unavailable. However, for each comparison, the original analysis had clearly used a paired t-test or a paired z-test on fold-averaged statistics from a single dataset and had reported an exact p-value and the exact cross-validation scheme. For *t*-tests, the degrees of freedom were also available.

From the reported p-value, we recovered the naïve (uncorrected) t-statistic (or z-statistic) and divided it by the Nadeau–Bengio correction factor $\sqrt{1 + J \cdot \frac{N_2}{N_1}}$, where $J$ is the number of fold-averaged statistics, and $N_2/N_1$ is the ratio of test-set size to training-set size for each cross-validation split (Supplementary Methods S6). We obtained the corrected p-value from the corresponding reference distribution, using the reported degrees of freedom for t-tests and the standard normal distribution for z-tests. This correction is equivalent to applying the corrected resampled t-test (or z-test) to the original fold-averaged statistics (which were not available to us) without additional assumptions arising from missing information.

By construction, Tier A′ included only comparisons for which a paired test had clearly been used. Comparisons analyzed with an unpaired test were re-assigned to Tier C. Without the underlying fold-averaged performance values, we could not apply the corrected resampled paired t-test even if there was fold correspondence. Comparisons for which it was unclear whether the test was paired were assigned to Tier C for the same reason.

Some Tier-A′ comparisons had been explicitly evaluated using one-sided paired tests. We did not retrospectively assess whether a one-sided test was scientifically justified, but retained the directionality specified by the original studies. For these comparisons, we recovered the naïve statistic from the reported one-sided p-value, divided it by the Nadeau–Bengio correction factor, and obtained the corrected one-sided p-value.

Some Tier A′ comparisons were clearly based on paired tests but we could not determine whether the tests were one- or two-sided. We therefore performed both one- and two-sided corrections. If both corrected p-values were $\geq 0.05$, the comparison was classified as spurious. If both p-values were $< 0.05$, the comparison was classified as retaining significance. If the two corrected p-values yielded different significance verdict, the comparison was re-assigned to Tier B, because determining whether to apply a one- or two-sided tests would require an additional assumption.

#### S3. Confidence interval calculation for false positive rate

In the main text, Methods "FPR simulation" describes the simulations to estimate the false positive rate (FPR) of various statistical tests across $m$ independent scenarios. To compute confidence intervals for the FPR, we treated the FPR as a binomial proportion, representing the number of significant detections ($p < 0.05$) out of $m$ independent tests.

Since binomial proportions, especially at the extremes or with limited sample sizes, often deviate from the normal approximation, we used the Wilson score interval with continuity correction, as recommended by a previous study (Newcombe, 1998). Given a total of $m$ independent realizations and $n_s$ observed rejections of the null hypothesis, the sample proportion is $\hat{p} = \frac{n_s}{m}$. For a 95% confidence level, the critical value from the standard normal distribution is $z_{.025} = 1.96$. The continuity-corrected Wilson interval was then computed as (Newcombe, 1998):

$$w_{cc}^{-} = \max\left\{0, \frac{2m\hat{p}+z_{.025}^{2}-\left[z_{.025}\sqrt{z_{.025}^{2}-\frac{1}{m}+4m\hat{p}(1-\hat{p})+(4\hat{p}-2)}+1\right]}{2\left(m+z_{.025}^{2}\right)}\right\} \quad \text{(S3.1)}$$

$$w_{cc}^{+} = \min\left\{1, \frac{2m\hat{p}+z_{.025}^{2}+\left[z_{.025}\sqrt{z_{.025}^{2}-\frac{1}{m}+4m\hat{p}(1-\hat{p})-(4\hat{p}-2)}+1\right]}{2\left(m+z_{.025}^{2}\right)}\right\} \quad \text{(S3.2)}$$

This method provides more accurate interval bounds than the standard Wald interval, particularly when the proportion is near 0 or 1, or when $m$ is small (Newcombe, 1998).

#### S4. Datasets, algorithms & hyperparameters for the main simulation scheme

##### S4.1 Hyperparameter selection strategies

We considered three hyperparameter selection strategies. The first strategy used a fixed, predetermined set of hyperparameters for each algorithm based on scikit-learn defaults. The second strategy used an internal 80%/20% train-validation split within the training set. For each candidate hyperparameter configuration, the model was trained on the 80% subset and evaluated on the 20% validation set. The training-validation split was fixed across candidates, so validation performance was comparable. The hyperparameter configuration with the best validation performance was then used to train a final model on the full training set and evaluated in the test set.

The third strategy used nested cross-validation. For each candidate hyperparameter configuration, 2-fold cross-validation was performed on the training set. The cross-validation split was fixed across candidates, so performance was comparable. The hyperparameter configuration with the best cross-validation performance was then used to train a final model on the full training set and evaluated in the test set.

##### S4.2 EMNIST

We used the "digits" subset of the Extended MNIST (EMNIST) dataset (Cohen et al., 2017), which comprised 240,000 grayscale images (28×28 pixels) in the training set and 40,000 grayscale images (28×28 pixels) in the test set. For the current study, we only utilized the training set. There were 24,000 images per digit class (0–9). The machine learning task was to classify each image into one of the 10-digit classes.

For the prediction task, we used multi-class logistic regression classifier and the performance evaluation metric was classification accuracy, defined as the fraction of samples in the test set that was classified correctly. The hyperparameter of interest was $C$, the inverse regularization strength. In the fixed hyperparameter setting, $C$ was set to 1.0, while in the hyperparameter tuning settings, $C$ was searched across the following values: [0.0001, 0.0005, 0.001, 0.005, 0.01, 0.05, 0.1, 0.5, 1, 5, 10, 50, 100, 500, 1000, 5000, 10000].

##### S4.3 UK Biobank

We analyzed data from 36,454 UK Biobank participants (Alfaro-Almagro et al., 2018), consistent with our previous studies (He et al., 2022; Wulan et al., 2024). The prediction task was regression, with age (defined as MRI scan date minus birth year and month) as the target variable. The volumes of 101 cortical and subcortical gray-matter regions generated by the FreeSurfer software (Fischl, 2012) were features used for prediction.

Raw volumetric measures were first normalized by dividing by the intracranial volume of each participant. Feature z-normalization was then performed using the mean and standard deviation computed from a held-out subset of 454 randomly sampled participants, which were subsequently applied to the remaining 36,000 participants. This yielded a final dataset of 36,000 individuals, each represented by 101 standardized features.

For the prediction task, we used linear ridge regression and the performance evaluation metric was the coefficient of determination in the test set (Wright, 1921). The hyperparameter of interest was the regularization parameter $\alpha$. In the fixed hyperparameter setting, $\alpha$ was set to 10, and for hyperparameter tuning, $\alpha$ was searched across the following values: [0.0001, 0.0005, 0.001, 0.005, 0.01, 0.05, 0.1, 0.5, 1, 1.5, 2, 3, 4, 5, 10, 15, 20, 30, 40, 50, 100, 150, 200, 300, 400, 500, 1000, 2000, 5000, 10000].

##### S4.4 Covertype

The Covertype dataset enabled the benchmarking of forest cover type classification based on cartographic variables derived from remote sensing and US Forest Service data. Each instance corresponded to a 30×30 meter cell in one of four wilderness areas within the Roosevelt National Forest, Colorado (Blackard, 1998).

The dataset contained 581,012 instances and 54 features, including both continuous variables (elevation, slope, aspect, hillshade values, distance to hydrology/roads) and binary variables encoding soil types and wilderness area indicators. Labels represented one of seven forest cover types, derived from the USFS Region 2 Resource Information System (RIS).

Following previous studies (Collobert et al., 2001), we constructed a binary classification task: class 1 included all cover types except for type 2, while class 2 included only cover type 2 (commonly associated with lodgepole pine in Rawah and Comanche Peak). This transformation resulted in a more balanced distribution: 297,711 samples in class 1 and 283,301 samples in class 2.

For the prediction task, we used the Extra Trees classifier and the performance evaluation metric was classification accuracy, defined as the fraction of samples in the test set that was classified correctly. Because Covertype was the only binary-classification dataset, we additionally computed AUC to evaluate the DeLong test.

In the fixed hyperparameter setting, we used scikit-learn defaults with 100 estimators. For hyperparameter tuning, ranges were chosen following prior studies (Komer et al., 2014; Grinsztajn et al., 2022) and optimized using the Tree-structured Parzen Estimator (TPE; Bergstra et al., 2011, 2013). Candidate configurations were evaluated using either the internal training-validation split or 2-fold cross-validation described in Supplementary Methods S4.1.

##### S4.5 KEGG

The KEGG (Kyoto Encyclopedia of Genes and Genomes) Metabolic Pathway database describes interactions between biochemical compounds, enzymes, and genes. These pathways can be represented as graphs using two models: (i) the reaction network, where nodes correspond to substrates and products, and edges correspond to catalyzing genes or enzymes; and (ii) the relation network, where compounds form edges between gene/enzyme nodes (Naeem & Asghar, 2011). A large set of pathway entries were parsed and transformed into graphs using both representations.

The dataset provided graph-based features extracted using Cytoscape (Shannon et al., 2003), a software platform for biological network visualization and analysis. There were a total of 53,414 samples, each described by 20 topological features (e.g., degree centrality, betweenness centrality, and clustering coefficient). In the current study, the learning objective was to predict the clustering coefficient of each network instance, making this a regression task.

For the prediction task, we used the Extra Trees regressor and the performance evaluation metric was the mean squared error, computed by averaging the squared error across all test samples.

In the fixed hyperparameter setting, we used scikit-learn defaults with 100 estimators. For hyperparameter tuning, ranges were chosen following prior studies (Komer et al., 2014; Grinsztajn et al., 2022) and optimized using the Tree-structured Parzen Estimator (TPE; Bergstra et al., 2011, 2013). Candidate configurations were evaluated using either the internal training-validation split or 2-fold cross-validation described in Supplementary Methods S4.1.

The KEGG Metabolic Pathway dataset was the only dataset in which performance was recorded at the individual sample level (squared error per test sample), rather than aggregated across the entire test set. It was therefore used to compare sample-level and fold-averaged statistical tests.

#### S5. Definitions and implementation of statistical tests evaluated

We consider the problem of testing whether one machine learning model statistically outperforms another machine learning model in a given dataset based on cross-validation. For example, in K-fold cross-validation, the dataset of $N$ samples is partitioned into $K$ mutually exclusive folds. Each fold serves once as a test set, while the remaining $K-1$ folds form the training set. The training set is used to train each model, and model performance is evaluated in the test set. This yields $K$ pairs of performance metrics on which we want to perform a statistical test.

If we repeat K-fold cross-validation $R$ times, then we have $J = K \times R$ pairs of performance metrics. In general, we assume the statistical test operates on a vector $\boldsymbol{D}$ (of length $J$), where each element of vector $\boldsymbol{D}$ corresponds to a fold-level performance difference between the two models for a particular test fold.

For example, in the case of performing 10-fold cross-validation once, the vector $\boldsymbol{D}$ will be of length $J = 10$. If we perform 10-fold cross-validation 30 times, vector $\boldsymbol{D}$ will be of length $J = 300$. In the case of 80-20 Monte Carlo cross-validation where the dataset is repeatedly split into 80% training set and 20% test set 300 times, the vector $\boldsymbol{D}$ will again be of length $J = 300$.

In the following subsections, we briefly summarize various statistical tests used in the literature. We note that the resampled paired t-test (Supplementary Methods S5.1), Wilcoxon signed-rank test (Supplementary Methods S5.2), permutation test (Supplementary Methods S5.3) and the DeLong test (Supplementary Methods S5.4) are expected to have elevated FPR. The corrected resampled t-test (Supplementary Methods S5.5), the 5×2 tests (Supplementary Methods S5.6) and the empirical test of differences (Supplementary Methods S5.7) implicitly or explicitly try to account for between-fold correlation. The bootstrap may or may not be valid depending on how it was implemented (Supplementary Methods S5.8).

##### S5.1 Resampled paired t-test

As demonstrated in our meta-analysis, the most common statistical test used in comparing models in cross-validation is the paired t-test on the fold-level accuracy differences, referred to as the “resampled paired t-test” (Nadeau & Bengio, 2003). Given a vector $\boldsymbol{D}$ of $J$ fold-level performance differences between two models, we test the null hypothesis $\mu = 0$, where $\mu$ is the true expected performance difference between the two models. The paired t-test utilizes the following statistic:

$$T = \frac{\bar{D}}{\sqrt{\frac{1}{J} S^2}} \quad \text{(S5.1)}$$

where $\bar{D}$ is the mean of vector $\boldsymbol{D}$, $S^2 = \frac{1}{J-1} \sum_j \left(D_j - \bar{D}\right)^2$ is the sample variance of vector $\boldsymbol{D}$.

The paired t-test assumes independence among the elements of vector $\boldsymbol{D}$, an assumption that is violated under cross-validation. In K-fold cross-validation, test folds are mutually disjoint across iterations, but the training sets overlap substantially (for $K > 2$), and the test fold in one iteration contributes to the training set in another iteration. In repeated K-fold cross-validation, the folds are redefined, but between-fold correlation remains. Similarly, in Monte Carlo cross-validation, where the dataset is repeatedly split into training and test sets, there will be overlap of both training and test sets across random dataset splits.

Because the correlations among fold-level statistics are positive, the paired t-test is expected to have elevated false positive rates, as demonstrated empirically in Fig. 5. A more detailed discussion of this issue can be found in Supplementary Methods S6.

##### S5.2 Wilcoxon signed-rank test

As demonstrated in our meta-analysis, the second most common statistical test used in comparing models in cross-validation is the Wilcoxon signed-rank or Wilcoxon rank-sum test. The Wilcoxon rank-sum test is also referred to as the Mann–Whitney U test, and is the unpaired version of the Wilcoxon signed-rank test. The Wilcoxon signed-rank test is a nonparametric alternative to the paired t-test, designed to test whether the median of paired differences is zero (Wilcoxon, 1945; Siegel, 1956).

Similar to the paired t-test, the Wilcoxon signed-rank test takes in a vector of paired differences and ranks the absolute differences between the $J$ paired observations. The test then computes a test statistic based on the sum of these ranks, weighted by the sign of the original differences. The resulting test statistic is then compared against a reference distribution to obtain a p-value (Japkowicz & Shah, 2011).

Like the paired t-test, the Wilcoxon signed-rank test assumes independence among the paired differences. Because the fold-level statistics are positively correlated, the Wilcoxon signed-rank test is expected to have elevated false positive rates, as demonstrated empirically in Fig. 5.

##### S5.3 Sign-flip permutation test

We found that there are two different permutation tests used in literature, but both are invalid. One permutation test is invalid not because of ignoring between-fold correlation but because it permuted the wrong variables. We excluded this permutation test from our meta-analysis. For details, see Supplementary Results. We are not aware of a valid permutation test for comparing predictive performance under cross-validation.

We will describe the second permutation test here. Once again, the null hypothesis is $\mu = 0$, where $\mu$ is the true expected difference between the two models. Given a vector $\boldsymbol{D}$ of $J$ fold-level performance differences between two models, let $\overline{D}$ be the mean of vector $\boldsymbol{D}$. A permutation of the data generates a null distribution: For each iteration, the sign of each element of vector $\boldsymbol{D}$ is flipped independently with probability 0.5, thus creating a new vector $\boldsymbol{D}_0$. The mean of $\boldsymbol{D}_0$ is then computed, contributing a single null value to the null distribution. After many iterations (e.g., 10,000), the original statistic $\overline{D}$ is compared against the null distribution to generate a p-value.

This permutation test wrongly assumes the elements of vector $\boldsymbol{D}$ are independent. Therefore, similar to the naïve t-test, we expect the permutation test to yield higher FPR, as demonstrated empirically in Supplementary Fig. 6. A more detailed discussion of this issue can be found in Supplementary Methods S7.

##### S5.4 DeLong test

The DeLong test is a statistical procedure specific to model comparison based on a Receiver Operating Characteristic (ROC) curve (DeLong et al., 1988). The test only applies to binary classification tasks, in which the model predicts one of two possible outcomes (e.g., spam vs. non-spam, disease vs. no disease). In such tasks, classifier performance can be summarized by an ROC curve, which plots the true positive rate (TPR) against the false positive rate (FPR) across different decision thresholds. To compare two ROC curves, the Area Under the ROC Curve (AUC) is used as a scalar performance metric, with larger AUC values indicating better performance.

The DeLong test evaluates whether the difference in AUCs between two algorithms is statistically significant. Specifically, the test statistic is constructed as the difference between the two estimated AUCs divided by an estimate of its standard error, yielding an asymptotic z-statistic. The AUC estimator is a function of the pairwise comparisons between positive and negative samples and can be expressed as a normalized Mann–Whitney U-statistic. The

DeLong test estimates the 2×2 covariance matrix of the two AUC estimators using an influence-function-based approach for vectors of U-statistics (DeLong et al., 1988). The joint asymptotic normality of the AUC estimators justifies the use of a normal approximation to compute p-values.

Although the variance estimation in the DeLong test is mathematically more involved than that of simpler tests, the U-statistic variance derivation underlying the DeLong test assumes that predictions are independent across data samples, which is violated under cross-validation. We also note that the DeLong test is based on sample-level statistics, as opposed to fold-averaged statistics (see Methods "Cross-validation primer"). As such, we expect the DeLong test to have elevated FPR, as empirically shown in Supplementary Fig. 2.

##### S5.5 Corrected resampled paired t-test

The corrected resampled paired t-test was proposed to adjust the variance estimator to reflect the positive correlation between cross-validation folds (Nadeau & Bengio, 2003). Although originally developed for Monte Carlo cross-validation, the corrected paired t-test has also been applied to repeated K-fold cross-validation (Bouckaert & Frank, 2004).

Similar to previous tests, the corrected resampled t-test operates on the vector $\boldsymbol{D}$ of $J$ fold-level performance differences between two models. Again, we wanted to test the null hypothesis $\mu = 0$, where $\mu$ is the true expected difference between the two models. The corrected resampled t-test assumes that the correlation between folds is equal to the fraction of the full dataset used in the test fold, resulting in the following statistic:

$$T = \frac{\bar{D}}{\sqrt{\left(\frac{1}{J}+\frac{N_2}{N_1}\right)S^2}} \qquad \text{(S5.2)}$$

where $\bar{D}$ is the mean of vector $\boldsymbol{D}$, $S^2 = \frac{1}{J-1}\sum_j \left(D_j - \bar{D}\right)^2$ is the sample variance of vector $\boldsymbol{D}$, and $D_j$ refers to the $j$-th element of vector $\boldsymbol{D}$; $N_1$ and $N_2$ are the number of training and test samples, respectively, in a particular split of the dataset into training and test sets. The statistic $T$ is assumed to follow a Student's t distribution with $J-1$ degrees of freedom, from which a p-value can be computed.

A more detailed discussion of the corrected resampled t-test can be found in Supplementary Methods S6. One problem with the corrected resampled t-test is the assumption that the correlation between cross-validation folds is solely due to the overlap in the test set. In reality, the correlation likely depends on the complex interaction between the machine learning algorithms and the dataset being analyzed. We also note that some studies applied the corrected resampled t-test wrongly, resulting in a high FPR (Supplementary Fig. 6; Supplementary Methods S8).

##### S5.6 5×2 paired t-test and 5×2 paired F-test

The 5×2 t-test (Dietterich, 1998) and 5×2 F-test (Alpaydın, 1999) involve 5 repetitions of 2-fold cross-validation, hence the name 5×2. For each repetition $r \in \{1,2,3,4,5\}$, the data is randomly split into two halves A and B. Both models are trained on set A and evaluated on set B, resulting in performance difference $D_{Br}$; likewise, both models are trained on set B and evaluated on set A, resulting in performance difference $D_{Ar}$.

The 5×2 t-test utilizes the following statistic (Dietterich, 1998):

$$T = \frac{D_{A1}}{\sqrt{\frac{1}{5}\sum_{r=1}^{5} S_r^2}} \tag{S5.3}$$

where $S_r^2 = (D_{Ar} - D_{Br})^2/2$. The statistic is assumed to follow a Student's t distribution with 5 degrees of freedom, from which a p-value can be computed. Supplementary Methods S9.1 explains why the 5×2 t-test will have a much lower false positive rate than the naïve t-test. However, the numerator in the T statistic $D_{A1}$ only uses a single accuracy difference from the first fold of the first 2-fold cross-validation, thus discarding information from the other folds.

To address the inefficiency of the 5×2 t-test, Alpaydın (1999) proposed the 5×2 paired F-test, which uses the following statistic:

$$F = \frac{\sum_{r=1}^{5}(D_{Ar}^2 + D_{Br}^2)}{2\sum_{r=1}^{5} S_r^2} \tag{S5.4}$$

where $S_r^2 = (D_{Ar} - D_{Br})^2/2$ (same as the 5×2 t-test). The statistic is assumed to follow the F-distribution with 10 and 5 degrees of freedom in the numerator and denominator respectively, from which a p-value can be defined. Supplementary Methods S9.2 explains why the 5×2 F-test will have a much lower false positive rate than the naïve t-test. Intuitively, since the numerator uses fold-level differences from all 5 repetitions of 2-fold cross-validation (as opposed to just a single fold in the 5×2 t-test), the 5×2 F-test might be a more sensitive test than the 5×2 t-test (Alpaydın, 1999).

Indeed, our results suggest that both the 5×2 t-test and 5×2 F-test reliably control FPR, and that the 5×2 F-test exhibits higher power than the 5×2 t-test. However, both tests had lower power than SHARP.

S5.7 Empirical test of differences

Given a vector $\boldsymbol{D}$ of $J$ fold-level performance differences between two models, suppose the number of positive entries is greater than the number of negative entries. The empirical test of differences then computes the p-value as the fraction of negative entries multiplied by two.

On the other hand, suppose the number of positive entries is less than the number of negative entries. The empirical test of differences then computes the p-value as the fraction of positive entries multiplied by two.

The empirical test of differences has been utilized by a few brain imaging studies (Dhamala et al., 2021; Parkes et al., 2021a, 2021b), which referred to it as the exact test of differences. Our results suggest that the empirical test of differences reliably controls false positives (Fig. 6) while having the lowest power among statistical tests that account for between-fold correlation (Fig. 7). Supplementary Methods S10 provides a theoretical explanation for these findings.

S5.8 Three bootstrap variants

Bootstrap is a common technique for estimating confidence intervals (Efron & Tibshirani, 1994; DiCiccio & Efron, 1996). There are different bootstrapping variants for cross-validation (Raschka, 2018; Cai et al., 2025). Our current meta-analysis excluded all studies that only reported a confidence interval (PRISMA criterion 4; Methods "Meta-analysis"). As such, all studies using bootstrap were also excluded. However, we note that only one study described their bootstrapping procedure in sufficient details for us to evaluate its validity (Peneder et al., 2021).

Below we outlined three broad bootstrap strategies that we evaluated in the current study. The three strategies were not meant to be exhaustive but were aimed to cover the good and bad variants of bootstrap. First, we implemented the bootstrap procedure from Raschka (2018), since it coincided with the exemplary study in our meta-analysis that clearly described the procedure (Peneder et al., 2021). More specifically, given a dataset of $N$ samples, we bootstrapped (sampled with replacement) $N$ samples from the dataset, which served as the training set. We note that on average, 63% of the training samples will be unique. The test set comprised samples that were not included in the training set.

To compare two models, the bootstrapped training set was used to train each model and the trained models were evaluated on the test set, yielding a single bootstrapped difference value. The entire bootstrapping procedure was repeated $M$ times, and the 95% confidence interval was constructed from the $M$ performance difference estimates based on the 2.5 and 97.5 percentiles. If the 95% confidence interval did not cover zero, we considered the difference to be statistically significant. We referred to this bootstrapping procedure as "bootstrap-orig". For the evaluation of FPR and power, we followed the procedure in Methods "Simulations to evaluate FPR, statistical power and confidence intervals" with $M$ being set to 300.

The other two bootstrapping procedures performed bootstraps after normal cross-validation. For both approaches, given a vector $\boldsymbol{D}$ of $J$ fold-level performance differences, bootstrapping (sampling with replacement) was performed on the entries of vector $\boldsymbol{D}$, yielding a new vector $\boldsymbol{D}^*$ of 1000 bootstrapped fold-level differences. An empirical test of differences (Supplementary Methods S5.7) was applied to $\boldsymbol{D}^*$, which we referred to as bootstrap-ET. Alternatively, the naïve paired t-test (Supplementary Methods S5.1) was applied to $\boldsymbol{D}^*$, which we referred to as bootstrap-t.

Our results suggest that bootstrap-orig and bootstrap-ET reliably control FPR, but bootstrap-t has high FPR (Supplementary Fig. 6). However, both bootstrap-orig and bootstrap-ET had markedly worse power than SHARP, with bootstrap-ET being the least powerful (Supplementary Fig. 8).

#### S6. Correlation induced by cross-validation & limitations of naïve & corrected t-tests

We begin by characterizing the correlation structure of fold-level performance differences from a single run of K-fold cross-validation. The data split for the K-fold cross-validation is identical for both models, so there is fold-level correspondence between the two models. In each of $K$ iterations we train the two models on $K{-}1$ folds and test the two resulting models on the remaining fold. Therefore, each test fold yields a performance metric difference, and any pair of performance differences is correlated due to overlap of training data (for $K > 2$) and overlap between testing data in one fold and the training data for a different fold. Throughout, we denote $\boldsymbol{D}$ as the length-$J$ vector of performance metric differences; here $J = K$.

Now consider Monte Carlo cross-validation, where the dataset is repeatedly divided into training and test sets $J$ times, again leading to $J$ fold-level differences. In this case, there is overlap in both training and testing data between folds. The performance metric differences $\boldsymbol{D}$ are equally correlated, just like a single run of K-fold cross-validation.

Assuming the performance metric differences follows a Gaussian distribution, we can write

$$\boldsymbol{D} \sim N(\mu \mathbf{1}, \boldsymbol{\Sigma}) \tag{S6.1}$$

where $\mu$ is a scalar representing the true population average performance metric difference. $\mathbf{1}$ is a length-$J$ column vector of ones, and

$$\boldsymbol{\Sigma} = \sigma^2 \begin{bmatrix} 1 & \rho & \rho \\ \rho & \ddots & \rho \\ \rho & \rho & 1 \end{bmatrix} \tag{S6.2}$$

is the $J \times J$ covariance matrix, $\sigma^2$ is the variance of a given fold-level statistic and $\rho$ is the correlation between the fold-level statistics. The diagonals of $\boldsymbol{\Sigma}$ are $\sigma^2$, while all off-diagonal entries are $\sigma^2\rho$. By symmetry of the cross-validation procedure, $\boldsymbol{\Sigma}$ has a compound symmetric correlation structure, where each fold is correlated to a different fold with correlation $\rho$.

To perform a statistical test of whether the mean of a Gaussian variable is statistically different from zero, we need an estimate of the mean and the variance of the estimator of the mean. In the current cross-validation setup, the mean accuracy difference is $\bar{D} = \frac{1}{J}\mathbf{1}^\top \boldsymbol{D}$, which is an unbiased estimator of $\mu$, i.e., $\mathrm{E}(\bar{D}) = \mu$.

The variance of the estimator $\bar{D}$ is not easy to estimate (Nadeau & Bengio, 2003). Based on Equations S6.1 and S6.2 (above), the variance of the estimator $\bar{D}$ is given by

$$\mathrm{Var}(\bar{D}) = \frac{1}{J^2}\mathbf{1}^\top \boldsymbol{\Sigma} \mathbf{1} = \frac{\sigma^2}{J^2}(J + J(J-1)\rho) = \sigma^2\left(\frac{1}{J} + \frac{J-1}{J}\rho\right). \tag{S6.3}$$

However, we have two unknown variance parameters $\sigma^2$ and $\rho$ and a single sufficient statistic for the variance, the sample variance $S^2$. The sample variance of $\boldsymbol{D}$ can be written

$$S^2 = \frac{1}{J-1}\sum_j \left(D_j - \bar{D}\right)^2 = \frac{1}{J-1}\boldsymbol{D}^\top\left(\mathbf{I} - \frac{1}{J}\mathbf{J}\right)\boldsymbol{D}, \tag{S6.4}$$

where $D_j$ is the $j$-th entry of $\boldsymbol{D}$, $\mathbf{I} - \frac{1}{J}\mathbf{J}$ is the centering matrix, $\mathbf{I}$ is the identity and $\mathbf{J}$ is a square matrix of 1's. Unfortunately, $S^2$ is a biased estimator of $\sigma^2$:

$$\begin{aligned} \mathrm{E}(S^2) &= \frac{1}{J-1}\mathrm{tr}\left(\boldsymbol{\Sigma}\left(\mathbf{I} - \frac{1}{J}\mathbf{J}\right)\right) \\ &= \frac{1}{J-1}\left(J\sigma^2 - \frac{1}{J}\left(J\sigma^2(1 + (J-1)\rho)\right)\right) \\ &= \sigma^2(1-\rho) \end{aligned} \tag{S6.5}$$

where we have used the following result: for a given random vector $\boldsymbol{x}$ with mean $\boldsymbol{\mu}$ and covariance $\boldsymbol{\Sigma}$ and a constant matrix $\mathbf{A}$, $\mathrm{E}(\boldsymbol{x}^\top \mathbf{A}\boldsymbol{x}) = \mathrm{tr}(\mathbf{A}\boldsymbol{\Sigma}) + \boldsymbol{\mu}^\top \mathbf{A}\boldsymbol{\mu}$.

Since we expect the correlation $\rho$ to be positive between cross-validation folds, $1 - \rho$ is smaller than one and so the sample variance $S^2$ underestimates $\sigma^2$. The relationship between $\mathrm{E}(S^2)$ and $\mathrm{Var}(\bar{D})$ is thus given by (Nadeau & Bengio, 2003):

$$\mathrm{Var}(\bar{D}) = \left(\frac{1}{J} + \frac{\rho}{1-\rho}\right)\mathrm{E}(S^2) \tag{S6.6}$$

$$= \frac{1}{J}\mathrm{E}(S^2) + \frac{\rho}{1-\rho}\mathrm{E}(S^2) \tag{S6.7}$$

Since the correlation $\rho$ is expected to be positive between cross-validation folds, the second term in Equation S6.7 is positive. A conventional paired t-test applied in this setting is called a resampled paired t-test, and has the form:

$$T = \frac{\bar{D}}{\sqrt{\frac{1}{J}S^2}}. \tag{S6.8}$$

Therefore, when studies directly apply the paired-sample t-test to compare model performance, they are using only the first term in Equation S6.7, which is thus an underestimate of the variance of the mean estimator $\mathrm{Var}(\bar{D})$. Consequently, the resampled paired-sample t-test is expected to yield an inflated false positive rate.

Equation S6.6 provides the form of a bias correction needed to be used in the corrected resampled t-test (Nadeau & Bengio, 2003), which is to multiply the sample variance $S^2$ by $\frac{1}{J} + \frac{\rho}{1-\rho}$ (instead of $\frac{1}{J}$ as in Equation S6.8). Nadeau and Bengio assume that the correlation between folds $\rho = \frac{N_2}{N_1+N_2}$, where $N_1$ and $N_2$ are the number of training and test samples, respectively. Therefore, $\frac{\rho}{1-\rho}$ becomes $\frac{N_2}{N_1}$, leading to the following statistic:

$$T = \frac{\bar{D}}{\sqrt{\left(\frac{1}{J}+\frac{N_2}{N_1}\right)S^2}}, \tag{S6.9}$$

which Nadeau & Bengio (2003) call the "corrected resampled t-test".

Many studies perform repeated K-fold cross-validation, where the K-fold procedure is repeated $R$ times, resulting in $J = K \times R$ fold-level differences. While the corrected paired t-test has also been applied to repeated K-fold cross-validation (Bouckaert & Frank, 2004), the theoretical justification is weaker. The repeated K-fold cross-validation produces a $\boldsymbol{D}$ with a nested correlation structure, where the correlation between folds within a particular instance of K-fold cross-validation is different from the correlation between folds from two different repeats of K-fold cross-validation. Nevertheless, in practice we find there is no difference in FPR between the application of the corrected resampled t-test in Monte Carlo cross-validation and repeated K-fold cross-validation (Supplementary Fig. 6).

The strongest assumption of the corrected resampled t-test is that the correlation between folds $\rho$ is $\frac{N_2}{N_1+N_2}$. In reality, the correlation magnitude likely depends on the complex interaction between the machine learning algorithm and the dataset being analyzed. As shown in the main results, the corrected resampled t-test can have a slightly elevated FPR (Fig. 6), and worse power than the SHARP test (Fig. 7).

#### S7. How between-fold correlation invalidates the paired permutation test

There are two different permutation tests used in the literature, but both are invalid. The first permutation test is discussed in Supplementary Results. Here we discuss the second permutation test.

Given a vector $\boldsymbol{D}$ of $J$ performance differences between two models, let $\bar{D}$ be the mean of $\boldsymbol{D}$. To generate a null distribution, for each iteration, the sign of each entry of $\boldsymbol{D}$ is flipped with probability 0.5, thus creating a new vector $\boldsymbol{D}_u$. The mean of $\boldsymbol{D}_u$ is then computed, which we denote as $\bar{D}_u$. By repeating the procedure $U$ times, we obtain $U$ null values. The final two-sided p-value is computed as

$$p = \frac{1}{U}\sum_{u=1}^{U} \mathbb{I}(|\bar{D}_u| \geq |\bar{D}|) \quad \text{(S7.1)}$$

where $\mathbb{I}(|\bar{D}_u| \geq |\bar{D}|)$ is an indicator variable that is equal to one if $|\bar{D}_u| \geq |\bar{D}|$, and zero otherwise. Note that to avoid a p-value of 0, the permutation test p-value is typically computed by adding one to the numerator and denominator of Equation S7.1, i.e., $p = \frac{1}{U+1}(1 + \sum_{u=1}^{U} \mathbb{I}(|\bar{D}_u| \geq |\bar{D}|))$. However, for the purpose of the analysis below, we will use Equation S7.1.

This permutation test wrongly assumes the elements of $\boldsymbol{D}$ are independent. To show this more explicitly, we first note that based on Supplementary Methods S6, we can write:

$$\mathrm{E}(\bar{D}) = \mu \quad \text{(S7.2)}$$

$$\mathrm{Var}(\bar{D}) = \sigma^2\left(\frac{1}{J} + \frac{J-1}{J}\rho\right) \quad \text{(S7.3)}$$

For the null permutation distribution to be valid, we note that under the null hypothesis, we require the following conditions to be true:

$$\mathrm{E}(\bar{D}_u) = 0 \quad \text{(S7.4)}$$

$$\mathrm{Var}(\bar{D}_u) = \sigma^2\left(\frac{1}{J} + \frac{J-1}{J}\rho\right) \quad \text{(S7.5)}$$

To evaluate if the above conditions are met, let $D_j$ be the $j$-th element of $\boldsymbol{D}$. Then we can write

$$\bar{D}_u = \frac{1}{J}\sum_{j=1}^{J} s_j D_j \quad \text{(S7.6)}$$

where each $s_j$ independently takes values from $\{+1, -1\}$ with probability $\frac{1}{2}$. Therefore,

$$\begin{aligned}
\mathrm{E}(\bar{D}_u) &= \frac{1}{J}\sum_{j=1}^{J}\mathrm{E}(s_j)\mathrm{E}(D_j) \\
&= \frac{1}{J}\sum_{j=1}^{J} 0 \times \mu = 0
\end{aligned} \tag{S7.7}$$

$$\begin{aligned}
\mathrm{Var}(\bar{D}_u) &= \frac{1}{J^2}\mathrm{E}\left(\mathrm{Var}\left(\sum_{j=1}^{J} s_j D_j \,|D_j\right)\right) + \\
&\quad \frac{1}{J^2}\mathrm{Var}\left(\mathrm{E}\left(\sum_{j=1}^{J} s_j D_j \,|D_j\right)\right) \\
&= \frac{1}{J^2}\mathrm{E}\left(\sum_{j=1}^{J} D_j^2\right) + \frac{1}{J^2}\mathrm{Var}(0) \\
&= \frac{1}{J^2}\mathrm{E}(\boldsymbol{D}^\top \boldsymbol{D}) \\
&= \frac{1}{J^2}(\mathrm{tr}(\boldsymbol{\Sigma}) + \mu^2 J) = \frac{\sigma^2+\mu^2}{J},
\end{aligned} \tag{S7.8}$$

where we have used the following result: for a given random vector $\boldsymbol{x}$ with mean $\boldsymbol{\mu}$ and covariance $\boldsymbol{\Sigma}$, $\mathrm{E}(\boldsymbol{x}^\top \boldsymbol{x}) = \mathrm{tr}(\boldsymbol{\Sigma}) + \boldsymbol{\mu}^\top \boldsymbol{\mu}$.

Under the null hypothesis $\mu = 0$,

$$\mathrm{Var}(\bar{D}_u) = \sigma^2 \frac{1}{J} \leq \sigma^2 \left(\frac{1}{J} + \frac{J-1}{J}\rho\right) = \mathrm{Var}(\bar{D}) \tag{S7.9}$$

Therefore, the first condition above (Equation S7.4) is met, but the second condition (Equation S7.5) is not met. In particular, since we expect the elements of $\boldsymbol{D}$ to be positively correlated, then $\mathrm{Var}(\bar{D}_u) < \mathrm{Var}(\bar{D})$, suggesting that the null distribution is overly narrow. Therefore, the permutation is expected to have an elevated FPR, which we observe empirically (Supplementary Fig. 6).

#### S8. Incorrectly using the corrected resampled paired t-test leads to high FPR

In repeated K-fold cross-validation (CV), the dataset is partitioned into $K$ folds, and this procedure is repeated $R$ times using different random splits. This yields a vector $\boldsymbol{D}$ of $J$ (= $K$×$R$) fold-level differences. As described in Supplementary Methods S5.5, the corrected resampled paired t-test was proposed to account for between-fold correlation. The corrected paired t-test utilizes the following statistic:

$$T = \frac{\bar{D}}{\sqrt{\left(\frac{1}{J}+\frac{N_2}{N_1}\right)S^2}} \quad , \tag{S8.1}$$

where $\bar{D}$ is the average of the vector $\boldsymbol{D}$. $N_1$ and $N_2$ are the number of training and test samples respectively in a particular split of the dataset into training and test sets. The corrected paired t-test assumes that the correlation between folds is equal to the fraction of the full dataset used in the test fold. For example, if 10-fold cross-validation was repeated 30 times, then $\boldsymbol{D}$ is a vector of length 300, so $J$ = 300 and $N_2/N_1$=1/9. In this setup, $\rho = 1/10$ (implicitly). As shown in Fig. 6, the corrected t-test has slightly elevated FPR.

However, we have observed certain studies implementing an incorrect version of the corrected resampled t-test. For example, if 10-fold cross-validation was repeated 30 times, then instead of creating a vector $\boldsymbol{D}$ (of length 300), some studies took the matrix of fold-level differences of size 10×30, and averaged each column of the matrix, so they ended up with a vector $\boldsymbol{D}^*$ (of length 30). They then utilized the above T statistic (Equation S8.1) using the mean of vector $\boldsymbol{D}^*$ in the numerator (which is actually the same as $\bar{D}$) with $J$ = 30 and $N_2/N_1$=1/9.

In other words, the authors implicitly assumed the entries in $\boldsymbol{D}^*$ were correlated with $\rho = 1/10$. However, the entries in $\boldsymbol{D}^*$ (vector of length 30) were likely to be much more correlated than the entries in $\boldsymbol{D}$ (vector of length 300). As such, we expected this incorrect version of the corrected t-test to have significantly higher FPR than the correctly-implemented corrected t-test, which was confirmed in Supplementary Fig. 6.

#### S9. Conservativeness of 5×2 t-test and 5×2 F-test

Our results suggest that both 5×2 t-test (Dietterich, 1998) and 5×2 F-test (Alpaydın, 1999) reliably control FPR (Fig. 6 & Supplementary Fig. 6) and both tests had lower statistical power than the SHARP test (Fig. 7 & Supplementary Fig. 8). In this section, we provide some insights on why the 5×2 tests are more conservative than the resampled paired t-test.

Recall that the 5×2 t-test tests involve 5 repetitions of 2-fold cross-validation. For each repetition $r \in \{1,2,3,4,5\}$, the data is randomly split into two halves A and B. Both models are trained on set A and evaluated on test set B, resulting in performance difference $D_{Br}$. Both models are also trained on set B and evaluated on test set A, resulting in performance difference $D_{Ar}$.

##### S9.1 5×2 paired t-test

The 5×2 t-test utilizes the following statistic (Dietterich, 1998):

$$T = \frac{D_{A1}}{\sqrt{\frac{1}{5}\sum_{r=1}^{5} S_r^2}} \quad \text{(S9.1)}$$

where $S_r^2 = (D_{Ar} - D_{Br})^2/2$. The statistic is assumed to follow a Student's t distribution with 5 degrees of freedom, from which a p-value can be computed.

Following the notation in Supplementary Methods S6, we note that $\mathrm{E}(D_{Ar}) = \mathrm{E}(D_{Br}) = \mu$, the fold-level statistics have a variance of $\sigma^2$ and the correlation between fold-level statistics within a single run of two-fold cross-validation is $\rho$. For convenience, we can write $D_{Ar} = \mu + \epsilon_{Ar}$, and $D_{Br} = \mu + \epsilon_{Br}$, where $\epsilon_{Ar}$ and $\epsilon_{Br}$ are zero mean random variables with variance $\sigma^2$. Furthermore, $\mathrm{Cov}(\epsilon_{Ar}, \epsilon_{Br}) = \sigma^2\rho$. Note that there is also a correlation $\rho'$ across the 5 repeats of the 2-fold cross-validation, but this $\rho'$ is not necessary for the analysis below.

The numerator $D_{A1}$ has mean $\mu$ and variance $\sigma^2$. Ideally, we would like the denominator $\frac{1}{5}\sum_{i=1}^{5} \mathrm{S}_\mathrm{r}^2$ to be an unbiased estimate of the variance of the numerator. While Dietterich assumes that the $D_{Ar}$ and $D_{Br}$ are independent, we find that

$$\begin{aligned}
\mathrm{E}\left(\frac{1}{5}\sum_{r=1}^{5} S_r^2\right) &= \frac{1}{5}\sum_{r=1}^{5} \mathrm{E}\left(\frac{(D_{Ar}-D_{Br})^2}{2}\right) \\
&= \frac{1}{5}\sum_{r=1}^{5} \mathrm{E}\left(\frac{(\epsilon_{Ar}-\epsilon_{Br})^2}{2}\right) \\
&= \frac{1}{5}\sum_{r=1}^{5} \frac{1}{2}(\sigma^2 + \sigma^2 - 2\rho\sigma^2) \\
&= \sigma^2(1-\rho). \quad \text{(S9.2)}
\end{aligned}$$

Therefore, ideally the 5×2 t-statistic should be multiplied by $\sqrt{1-\rho}$ (but $\rho$ is unknown). When the 5×2 t-test was first introduced, it was demonstrated that correlation $\rho$ between two folds can be positive, which might cause the test to be liberal (Dietterich, 1998).

Let us contrast this with the paired t-test applied to a single run of K-fold cross-validation. Based on Supplementary Methods S6, the correction factor required for the paired t-test is $\sqrt{\frac{1}{K}/\left(\frac{1}{K}+\frac{\rho}{1-\rho}\right)} = \sqrt{\frac{1-\rho}{1+(K-1)\rho}}$ (based on Equation S6.6), which is a much stronger correction factor than the 5×2 test ($\sqrt{1-\rho}$). Therefore, the 5×2 t-test is more conservative than the paired t-test applied to a single run of K-fold cross-validation.

However, we note that although the 5×2 t-test is theoretically slightly liberal, in practice, the 5×2 t-test reliably controls FPR (Fig. 6), but its power is significantly worse than the SHARP test (Fig. 7).

S9.2 5×2 paired F-test

To address the inefficiency of the 5×2 t-test, Alpaydın (1999) proposed the 5×2 paired F-test, which uses the following statistic:

$$F = \frac{\sum_{r=1}^{5}(D_{Ar}^2 + D_{Br}^2)}{2\sum_{r=1}^{5} S_r^2} \quad \text{(S9.3)}$$

where $S_r^2 = (D_{Ar} - D_{Br})^2/2$ (same as the 5×2 t-test). The statistic is assumed to follow the F-distribution with 10 and 5 degrees of freedom in the numerator and denominator, from which a p-value can be defined.

We can check the validity of this F-test by comparing the expectation of numerator to that of denominator under the null hypothesis. Ideally, these two expectations should be equal. Under null hypothesis, $\mu = 0$, we have:

$$\mathrm{E}\left(\sum_{r=1}^{5}(D_{Ar}^2 + D_{Br}^2)\right) = 10\sigma^2 \quad \text{(S9.4)}$$

$$\mathrm{E}\left(2\sum_{r=1}^{5} S_r^2\right) = 10\sigma^2(1-\rho) \quad \text{(S9.5)}$$

The expectation of the denominator is smaller than the expectation of the numerator, suggesting that the 5×2 F-test might be slightly liberal, similar to the 5×2 t-test. However, in practice, the 5×2 F-test reliably controls FPR (Supplementary Fig. 6), but its power is significantly worse than the SHARP test (Supplementary Fig. 8).

#### S10. Conservativeness of the empirical test of differences

As explained in Supplementary Methods S5.7, given a vector $\boldsymbol{D}$ of $J$ fold-level differences between two models, the empirical test of differences constructs an empirical histogram from the entries in the vector $\boldsymbol{D}$ and computes the fraction of entries in which the (overall) worse model outperforms the (overall) better model. To derive a two-sided test p-value, the empirical test of differences then multiplies the fraction by two.

We explore why the empirical test of differences is conservative through the following framework. Assume the $J$ fold-level differences follow a Gaussian distribution. Given a single instance of K-fold cross-validation or Monte Carlo cross validation, the empirical distribution of the accuracy differences in $\boldsymbol{D}$ is approximately Gaussian, centered at $\overline{D}$ with spread measured by $S^2$, where $\mathrm{E}(S^2) = \sigma^2(1-\rho)$ (Equation S6.5). The sample variance reflects only the variability within vector $\boldsymbol{D}$ and not the marginal variance $\sigma^2$. Without loss of generality, suppose $\bar{D}$ is negative, then the empirical test of differences is essentially computing 2× the area of the right positive tail of this Gaussian distribution.

On the other hand, we note that an oracle-based paired t-test that has knowledge of $\rho$ can be thought of as computing 2× the area of the right positive tail (again, assuming negative $\bar{D}$) of a Gaussian distribution with mean $\bar{D}$ and variance given by $\mathrm{Var}(\bar{D}) = \left(\frac{1}{J} + \frac{\rho}{1-\rho}\right)\mathrm{E}(S^2)$ (Equation S6.6).

Considering these two distributions that generate p-values, both centered at $\bar{D}$ but with different variances, illustrates why the empirical test of differences is less powerful. The empirical test of differences' distribution has variance expected to be $\sigma^2(1-\rho)$, while the oracle paired t-test has variance that differs by a factor of $\left(\frac{1}{J} + \frac{\rho}{1-\rho}\right)$. Therefore, if $\left(\frac{1}{J} + \frac{\rho}{1-\rho}\right) < 1$, then the empirical test is likely to be more conservative than the oracle paired t-test. This condition can be re-written as $\rho < \frac{J-1}{2J-1}$, so if J is very large, then this is equivalent to $\rho < 0.5$. Therefore, if $\rho$ is much smaller than 0.5, then the empirical test of differences will be significantly less powerful than an oracle paired t-test.

Indeed, we find that the empirical test of differences reliably controls the FPR (Fig. 6), while having the worst power (Fig. 7), suggesting that $\rho$ is positive, but much smaller than 0.5.

#### S11. Two-algorithm simulation scheme

Our main results utilized a simulation scheme for evaluating FPR, in which the same algorithm was trained on two noisy versions of the same dataset. Similarly, statistical power was evaluated by training the same algorithm on a clean dataset and a noisy version of the same dataset. As a supplementary analysis, we considered the two-algorithm simulation scheme (Dietterich, 1998; Nadeau & Bengio, 2003). The drawback of the two-algorithm simulation scheme is that we have to assume the true difference between the two algorithms is known, in order to evaluate the FPR. However, in practice, the results of the two-algorithm simulations were very similar to the noisy-model simulations from the main results.

##### S11.1 Algorithm Selection

We considered four datasets: EMNIST, Covertype, KEGG Metabolic, and UK Biobank. For each dataset, we selected two algorithms to compare. Similar to the noisy-model simulation, we used the AutoML package PyCaret (Ali, 2020) to explore a set of algorithms available in the scikit-learn package with default hyperparameters (Pedregosa et al., 2011). Because the simulations were highly computationally expensive, similar to the main results, we prioritized algorithms that were fast to run. We also prioritized pairs of algorithms that exhibited a large performance difference.

More specifically, we randomly selected 20,000, 20,000, 10,000, and 5,000 samples from EMNIST, Covertype, KEGG Metabolic, and UK Biobank datasets, respectively. Each sampled dataset was then partitioned into disjoint subsets of 1,000 samples. For each candidate algorithm, we conducted a 5-fold cross-validation on each subset. The performance metric of interest (e.g., classification accuracy) was averaged across five folds and recorded. This procedure yielded 20, 20, 10, and 5 averaged performance metrics for EMNIST,

Covertype, KEGG Metabolic, and UK Biobank, respectively, which were then further averaged to obtain a single performance estimate per algorithm on each dataset. Additionally, we recorded and averaged the runtime required to complete one 5-fold cross-validation for each algorithm.

Based on the selection criteria outlined above, we chose the following algorithm pairs:

1. EMNIST: Extra Tree Classifier and Linear Discriminant Analysis
2. Covertype: Random Forest Classifier and Support Vector Machine with Linear Kernel
3. KEGG Metabolic: Extra Tree Regressor and K-Nearest Neighbors Regressor
4. UK Biobank: Extra Tree Regressor and K-Nearest Neighbors Regressor

For each dataset, the algorithm listed first in the pair achieved numerically better prediction performance. Furthermore, the data samples used to select the two algorithms were excluded for subsequent analyses to avoid biasing the FPR and power simulations in the following sections. We will first explain how to evaluate power and then discuss how FPR can be obtained.

##### S11.2 Simulation procedure for statistical power

Consistent with the main results, we conducted analyses on four datasets — EMNIST, UKB, Covertype, and KEGG metabolic pathway — using four sample sizes: $N$ = 100, 500, 1000, and 2000. For EMNIST, $N$ = 100 was excluded because with 10 digit classes, it was not possible to guarantee at least one instance of each class in the test set under the SHARP test, which required a split-half step within the cross-validation scheme. For UKB, $N$ = 2,000 was excluded because the maximum number of non-overlapping sampled datasets that could be drawn from UKB at this sample size was fewer than 20, which we considered too few to yield stable estimates. Therefore, across the four datasets and sample sizes, there were 14 scenarios.

For each sample size $N$, we drew $m$ non-overlapping sampled datasets of size $N$ from the full dataset. The value of $m$ was set to 100 when feasible; otherwise, it was set to the maximum number of non-overlapping sampled datasets that could be drawn. For each of $m$ sampled datasets and each of the two algorithms associated with the dataset (previous section), we performed either repeated K-fold cross-validation or Monte Carlo cross-validation. In classification tasks, stratified splitting was used to preserve label proportions across folds.

All data splits were kept identical across both algorithms, so that we ended up with a vector $\boldsymbol{D}$ of $J$ performance differences. In the case of the SHARP test, each of the $m$ non-overlapping datasets was first divided into two non-overlapping halves, and cross-validation was performed within each half. We note that SHARP operates on a vector $\boldsymbol{D}$ of $2J$ performance differences.

Finally, the vector $\boldsymbol{D}$ of differences between the two algorithms was used in different statistical tests with the null hypothesis $\mu = 0$, where $\mu$ is the true expected difference between the two trained models. Since the two algorithms have been selected so that their prediction performance was different from each other, a well-calibrated test should reject the null hypothesis. Statistical power was estimated as the fraction of $m$ sampled datasets in which the null hypothesis was rejected at a significance threshold of 0.05. We then averaged the power across 14 scenarios, resulting in an average power for each test.

##### S11.3 Simulation procedure for false positive rate (FPR)

To estimate the FPR, the simulation procedure is the same as evaluating statistical power. However, since the two algorithms were selected so that their prediction performance was different from each other, the FPR was more difficult to estimate since we expected the null hypothesis $\mu = 0$ to be rejected.

If we knew the true difference between the two algorithms $\mu_{\text{true}}$, we could instead evaluate the null hypothesis $\mu = \mu_{\text{true}}$, in which case rejection of the null hypothesis constituted a false positive. However, we note that $\mu_{\text{true}}$ was unknown. The value of $\mu_{\text{true}}$ was also likely to be different across sample sizes and datasets.

Therefore, for each of the 14 scenarios, we estimated $\mu_{\text{true}}$, by averaging the entries of the vector $\boldsymbol{D}$ to obtain $\overline{D}$, and then averaged $\overline{D}$ across the *m* non-overlapping datasets, yielding $\hat{\mu}_{\text{true}}$. For each statistical test, we then evaluated the null hypothesis $\mu = \hat{\mu}_{\text{true}}$. Rejection of the null hypothesis constituted a false positive.

FPR was estimated as the fraction of *m* sampled datasets in which the null hypothesis was rejected at a significance threshold of 0.05. The 95% confidence interval for the FPR was computed using the Wilson score interval with continuity correction (Newcombe, 1998); the formula is provided in Supplementary Methods S3. Therefore, for each statistical test, we ended up with 14 FPRs and 95% confidence intervals for the FPRs.

##### S11.4 Confidence interval of prediction performance differences

To estimate the confidence interval of performance difference between the two algorithms, we followed a similar procedure as the noisy-model simulations (Methods "Confidence Intervals for Performance Difference"). Recall that we had 14 scenarios in the two-algorithm simulation scheme. For each scenario and each of *m* non-overlapping datasets, we found the range $[\mu_L, \mu_U]$ (i.e., confidence interval), such that each null hypothesis $H_0: \mu = \mu_0$, $\mu_0 \in [\mu_L, \mu_U]$, could not be rejected, i.e., $p \geq 0.05$. Since we have *m* non-overlapping datasets, we had *m* confidence intervals in total. We then counted the fraction of times $\hat{\mu}_{\text{true}}$ (Supplementary Methods S11.3) fell within the confidence interval, which we referred to as the overall coverage rate. For a well-calibrated 95% confidence interval, $\mu_{\text{true}}$ should fall into the confidence interval 95% of the times.

#### S12. SHARP (Split-Half Analysis of Repeated Performance) tests

##### S12.1 Covariance structure of SHARP cross-validation

Our goal is to compare two machine learning models. Instead of the traditional cross-validation scheme, we randomly divide the dataset into two disjoint halves A and B. For each machine learning model, we then perform K-fold cross-validation in subsets A and B separately. The data split for the K-fold cross-validation is identical for both models, so there is fold-level correspondence between the two models. We then averaged the results across the K-fold cross-validation, resulting in model performance differences $D_{A1}$ and $D_{B1}$ respectively. This process is repeated *J* times, resulting in two vectors $\boldsymbol{D}_A = [D_{A1}, \dots, D_{AJ}]^\top$ and $\boldsymbol{D}_B = [D_{B1}, \dots, D_{BJ}]^\top$.

Suppose there are model hyperparameters to be estimated, then the K-fold cross-validation can be modified to become a nested cross-validation where hyperparameters are estimated in the inner cross-validation loop. Alternatively, we can perform Monte Carlo cross-validation, where the subset A (or B) is repeatedly divided into training, validation and test sets. Regardless of the type of cross-validation scheme, we end up with two vectors $\boldsymbol{D}_A = \left[D_{A1}, \dots, D_{AJ}\right]^\top$ and $\boldsymbol{D}_B = \left[D_{B1}, \dots, D_{BJ}\right]^\top$.

Let $\boldsymbol{D} = [\boldsymbol{D}_A^\top, \boldsymbol{D}_B^\top]^\top = \left[D_{A1}, \dots, D_{AJ},\ D_{B1}, \dots, D_{BJ}\right]^\top$, so $\boldsymbol{D}$ is a column vector of length $2J$. We denote $\mathrm{Var}\left(D_{Aj}\right) = \mathrm{Var}\left(D_{Bj}\right) = \sigma^2$. Because of the disjoint subsets A and B, for any iteration *j*, $\mathrm{Corr}\left(D_{Aj}, D_{Bj}\right) = 0$. On the other hand, for two different iterations *j* and *k*, $\mathrm{Corr}\left(D_{Aj}, D_{Ak}\right) = \mathrm{Corr}\left(D_{Bj}, D_{Bk}\right) = \mathrm{Corr}\left(D_{Aj}, D_{Bk}\right) = \rho$. Therefore, the covariance matrix of $\boldsymbol{D}$ can be written as

$$\boldsymbol{\Sigma}(\sigma^2, \rho) = \sigma^2 \begin{bmatrix} \boldsymbol{M} & \boldsymbol{C} \\ \boldsymbol{C} & \boldsymbol{M} \end{bmatrix} \qquad \text{(S12.1)}$$

where $\boldsymbol{M} \in \mathbb{R}^{J \times J}$ has ones on the diagonal and $\rho$ in all off-diagonal entries, and $\boldsymbol{C} \in \mathbb{R}^{J \times J}$ has zeros on the diagonal and $\rho$ in all off-diagonal entries:

$$M_{jk} = \begin{cases} 1, j = k \\ \rho, j \neq k \end{cases} \text{ or } \boldsymbol{M} = \begin{bmatrix} 1 & \rho & \rho \\ \rho & \ddots & \rho \\ \rho & \rho & 1 \end{bmatrix} \qquad \text{(S12.2)}$$

and

$$C_{jk} = \begin{cases} 0, j = k \\ \rho, j \neq k \end{cases} \text{ or } \boldsymbol{C} = \begin{bmatrix} 0 & \rho & \rho \\ \rho & \ddots & \rho \\ \rho & \rho & 0 \end{bmatrix} \qquad \text{(S12.3)}$$

S12.2 GLS estimator of $\mu$ reduces to the sample mean

We assume that the vector $\boldsymbol{D}$ (of length 2J) follows a Gaussian distribution:

$$\boldsymbol{D} \sim N(\mu \boldsymbol{1}, \boldsymbol{\Sigma}) \qquad \text{(S12.4)}$$

The generalized least-squares (and maximum-likelihood) estimator for $\mu$ and the associated variance are as follows (Yan & Su, 2009):

$$\hat{\mu}_{GLS} = (\boldsymbol{1}^\top \boldsymbol{\Sigma}^{-1} \boldsymbol{1})^{-1} \boldsymbol{1}^\top \boldsymbol{\Sigma}^{-1} \boldsymbol{D}, \qquad \text{(S12.5)}$$

Because each row of $\boldsymbol{\Sigma}$ sums to a constant value, we note that the vector $\boldsymbol{1}$ is an eigenvector of $\boldsymbol{\Sigma}$ (and $\boldsymbol{\Sigma}^{-1}$). Therefore, $\boldsymbol{\Sigma}^{-1}\boldsymbol{1} = (1/c)\boldsymbol{1}$ for some scalar c > 0, so

$$(\boldsymbol{1}^\top \boldsymbol{\Sigma}^{-1} \boldsymbol{1})^{-1} \boldsymbol{1}^\top \boldsymbol{\Sigma}^{-1} = \frac{1}{2J} \boldsymbol{1}^\top. \qquad \text{(S12.6)}$$

Note that $1/c$ is the sum of any row of $\boldsymbol{\Sigma}^{-1}$, and $c$ is the sum of any row of $\boldsymbol{\Sigma}$. Plugging Equation S12.6 back into Equation S12.5, we get

$$\hat{\mu}_{GLS} = \frac{1}{2J} \boldsymbol{1}^\top \boldsymbol{D} = \overline{D}, \qquad \text{(S12.7)}$$

thus showing that the GLS and sample mean coincide. We also note that $\mathrm{E}(\bar{D}) = \mu$, and furthermore, we have:

$$\begin{aligned}
\mathrm{Var}(\bar{D}) &= \mathrm{Var}\left(\frac{1}{2J}\mathbf{1}^{\mathrm{T}}\boldsymbol{D}\right) \\
&= \frac{1}{4J^2}\mathbf{1}^T\boldsymbol{\Sigma}\mathbf{1} \\
&= \frac{\sigma^2}{4J^2}[2J + 4J(J-1)\rho] \\
&= \sigma^2\left(\frac{1}{2J} + \frac{J-1}{J}\rho\right), \qquad \text{(S12.8)}
\end{aligned}$$

where we used the following result: for a random vector $\boldsymbol{x}$ with covariance matrix $\boldsymbol{\Sigma}$ and constant vector $\boldsymbol{a}$, $\mathrm{Var}(\boldsymbol{a}^{\mathrm{T}}\boldsymbol{x}) = \boldsymbol{a}^T\boldsymbol{\Sigma}\boldsymbol{a}$. Therefore, if we can estimate $\sigma^2$ and $\rho$, we will be able to estimate $\mathrm{Var}(\bar{D})$, and perform a statistical test. In the following sections, we outline different approaches to performing the statistical test.

S12.3 Method-of-moments (MoM) estimates and Wald test

In this section, we use the method of moments (MoM) to estimate $\sigma^2$ and $\rho$. The Wald test is then used to perform the statistical test. We first compute the mean within subsets A and B.

$$\bar{D}_A = \frac{1}{J}\sum_{j=1}^{J} D_{Aj}, \qquad \text{(S12.9)}$$

$$\bar{D}_B = \frac{1}{J}\sum_{j=1}^{J} D_{Bj}, \qquad \text{(S12.10)}$$

$$\bar{D} = \frac{\bar{D}_A + \bar{D}_B}{2}. \qquad \text{(S12.11)}$$

We then compute sample variances within subsets A and B:

$$S_A^2 = \frac{1}{J-1}\sum_{j=1}^{J}\left(D_{Aj} - \bar{D}_A\right)^2 \qquad \text{(S12.12)}$$

$$S_B^2 = \frac{1}{J-1}\sum_{j=1}^{J}\left(D_{Bj} - \bar{D}_B\right)^2. \qquad \text{(S12.13)}$$

Similar to Equation S6.5 in Supplementary Methods S6, $\mathrm{E}(S_A^2) = \mathrm{E}(S_B^2) = \sigma^2(1-\rho)$. Let us also define the between-subset sample variance:

$$\hat{\sigma}_\Delta^2 = \frac{1}{2J}\sum_{j=1}^{J}\left(D_{Aj} - D_{Bj}\right)^2 \qquad \text{(S12.14)}$$

Using Equation S9.2 in the 5×2 t-test (Supplementary Methods S9.1), except that under SHARP cross-validation scheme, $\mathrm{Cov}\left(D_{Aj}, D_{Bj}\right) = 0$, we get $\mathrm{E}(\hat{\sigma}_\Delta^2) = \sigma^2$. We then use the following MoM estimators of $\rho$ and $\sigma^2$:

$$\hat{\rho}_{\mathrm{MoM}} = \frac{\hat{\sigma}_\Delta^2 - \frac{1}{2}(S_A^2 + S_B^2)}{\hat{\sigma}_\Delta^2}, \qquad \text{(S12.15)}$$

$$\hat{\sigma}_{\mathrm{MoM}}^2 = \hat{\sigma}_\Delta^2 \qquad \text{(S12.16)}$$

Furthermore, by substituting $\hat{\rho}_{\mathrm{MoM}}, \hat{\sigma}^2_{\mathrm{MoM}}$ into Equation S12.8, we get $\widehat{\mathrm{Var}}(\bar{D}) = \hat{\sigma}^2_{\mathrm{MoM}}\left(\frac{1}{2J} + \frac{J-1}{J}\hat{\rho}_{\mathrm{MoM}}\right)$. Finally, the MoM Wald test uses the following statistic:

$$Z_{\mathrm{MoM}} = \frac{\bar{D}}{\sqrt{\hat{\sigma}^2_{\mathrm{MoM}}\left(\frac{1}{2J}+\frac{J-1}{J}\hat{\rho}_{\mathrm{MoM}}\right)}} \tag{S12.17}$$

Under the null hypothesis of equal performance between the two algorithms, $Z_{\mathrm{MoM}}$ is approximately distributed as $N(0, 1)$, and the corresponding two-sided p-value is given by $p = 2\left(1 - \Phi(|Z_{\mathrm{MoM}}|)\right)$, where $\Phi(\cdot)$ denotes the standard normal cumulative distribution function.

S12.4 Maximum likelihood (ML) estimation and Wald test

Recall that we assume $\boldsymbol{D} \sim N\left(\mu\mathbf{1}, \boldsymbol{\Sigma}(\sigma^2, \rho)\right)$. The log likelihood is given by

$$\ell(\mu, \sigma^2, \rho) = -\frac{1}{2}\log|\boldsymbol{\Sigma}| - \frac{1}{2}(\boldsymbol{D} - \mu\mathbf{1})^{\top}\boldsymbol{\Sigma}^{-1}(\boldsymbol{D} - \mu\mathbf{1}) + \text{constant} \tag{S12.18}$$

From Supplementary Methods S12.2, the ML estimate of $\mu$ is equal to $\bar{D}$. The remaining parameters $\sigma^2$ and $\rho$ can be estimated by maximizing $\ell(\bar{D}, \sigma^2, \rho)$, yielding $\hat{\sigma}^2_{\mathrm{ML}}$ and $\hat{\rho}_{\mathrm{ML}}$. In our implementation, we utilized the "minimize" function in the scipy.optimize package, and initialized the optimization with the method of moments estimates $\hat{\rho}_{\mathrm{MoM}}, \hat{\sigma}^2_{\mathrm{MoM}}$.

The MLE-based Wald statistic is

$$Z_{\mathrm{ML}} = \frac{\bar{D}}{\sqrt{\hat{\sigma}^2_{\mathrm{ML}}\left(\frac{1}{2J}+\frac{J-1}{J}\hat{\rho}_{\mathrm{ML}}\right)}} \tag{S12.19}$$

and the corresponding two-sided p-value is given by $p = 2\left(1 - \Phi(|Z_{\mathrm{ML}}|)\right)$, where $\Phi(\cdot)$ denotes the standard normal cumulative distribution function.

S12.5 Restricted maximum likelihood (ReML) and Wald test

Variance parameters estimated with maximum likelihood are well known to have bias when estimated alongside mean parameters, particularly in small samples (Lehmann & Casella, 1998; Hogg et al., 2013). Building a likelihood based on contrasts that remove the mean parameters while retaining all other information in the data gives estimates with reduced bias, which is known as Restricted Maximum Likelihood (ReML).

In the SHARP cross-validation scheme, recall that we assume $\boldsymbol{D} \sim N(\mu\mathbf{1}, \boldsymbol{\Sigma})$. ReML can be implemented by centering the mean of the distribution to zero. We define the following residual-forming matrix

$$\boldsymbol{R} = \mathbf{I}_{2J\times 2J} - \frac{1}{2J}\mathbf{J}_{2J\times 2J} \tag{S12.20}$$

where $\mathbf{I}$ is the identity matrix and $\mathbf{J}$ is the matrix of all ones. To ensure a full-rank transformation for likelihood evaluation, we drop the last row of $\boldsymbol{R}$, resulting in $\boldsymbol{R}^* = (\boldsymbol{R})_{1:(2J-1),1:2J}$. The transformed residuals have the distribution

$$\boldsymbol{R}^{*}\boldsymbol{D} \sim N(0, \boldsymbol{R}^{*}\boldsymbol{\Sigma}\boldsymbol{R}^{*\top}) \tag{S12.21}$$

Similar to the previous section (Supplementary Methods S12.4), we can maximize the new likelihood to estimate $\hat{\sigma}^2_{\text{ReML}}, \hat{\rho}_{\text{ReML}}$, which can be used to compute the corresponding Wald statistic

$$Z_{\text{ReML}} = \frac{\bar{D}}{\sqrt{\hat{\sigma}^2_{\text{ReML}}\left(\frac{1}{2J} + \frac{J-1}{J}\hat{\rho}_{\text{ReML}}\right)}} \tag{S12.22}$$

The two-sided p-value is given by $p = 2\big(1 - \Phi(|Z_{\text{ReML}}|)\big)$, where $\Phi(\cdot)$ denotes the standard normal cumulative distribution function.

S12.6 Score test (used in the main Methods)

We also consider the use of likelihood-based score test, which evaluates the slope of the log-likelihood (known as the score) with respect to $\mu$ at $\mu = 0$ (under our null hypothesis), with nuisance parameters $(\sigma^2, \rho)$ estimated under the null.

In our current implementation, we utilize the "minimize" function in the scipy.optimize package to optimize the log likelihood $\ell(0, \sigma^2, \rho)$ (from Equation S12.18 in Supplementary Methods S12.4) to estimate $\hat{\sigma}^2_0, \hat{\rho}_0$. The optimization was initialized with the method of moments estimates $\hat{\rho}_{\text{MoM}}, \hat{\sigma}^2_{\text{MoM}}$ from Supplementary Methods S12.3.

The score under the Gaussian model is

$$U(0) = \frac{\partial}{\partial \mu}\ell(\mu, \sigma^2, \rho)|_{\mu=0} = \left(\frac{\partial \ell}{\partial(\boldsymbol{D} - \mu\boldsymbol{1})}\right)^{\top} \cdot \frac{\partial(\boldsymbol{D} - \mu\boldsymbol{1})}{\partial \mu}|_{\mu=0} \tag{S12.23}$$

$$= -\frac{1}{2}\left(2\boldsymbol{\Sigma}^{-1}(\boldsymbol{D} - \mu\boldsymbol{1})\right)^{\top} \cdot (-\boldsymbol{1})|_{\mu=0} \tag{S12.24}$$

$$= (\boldsymbol{D} - \mu\boldsymbol{1})^{\top}\boldsymbol{\Sigma}^{-1}\boldsymbol{1}|_{\mu=0} = \boldsymbol{D}^{\top}\boldsymbol{\Sigma}^{-1}\boldsymbol{1} \tag{S12.25}$$

Equation S12.23 follows from chain rule and Equation S12.24 follows from matrix calculus identity $\frac{\partial}{\partial \boldsymbol{x}}(\boldsymbol{x}^{\top}\boldsymbol{\Sigma}^{-1}\boldsymbol{x}) = \left(\boldsymbol{\Sigma}^{-1} + \boldsymbol{\Sigma}^{-1^{\top}}\right)\boldsymbol{x} = 2\boldsymbol{\Sigma}^{-1}\boldsymbol{x}$, where $\boldsymbol{x} = \boldsymbol{D} - \mu\boldsymbol{1}$. The Fisher information under the null hypothesis is

$$I(0) = -\frac{\partial^2}{\partial^2 \mu}\ell(\mu, \sigma^2, \rho)|_{\mu=0} = -\frac{\partial}{\partial \mu}\left(\frac{\partial}{\partial \mu}\ell(\mu, \sigma^2, \rho)\right)|_{\mu=0} \tag{S12.26}$$

$$= -\frac{\partial}{\partial \mu}(\boldsymbol{D} - \mu\boldsymbol{1})^{T}\boldsymbol{\Sigma}^{-1}\boldsymbol{1}|_{\mu=0} = \boldsymbol{1}^{\top}\boldsymbol{\Sigma}^{-1}\boldsymbol{1} \tag{S12.27}$$

The ratio of the squared score to the Fisher's information (again, both evaluated under the null hypothesis) converges in distribution to a chi-squared distribution with one degree of freedom (Rao, 1948). Equivalently, the standardized score $U(0)/\sqrt{I(0)}$ converges in distribution to a standard normal.

Let $\hat{\boldsymbol{\Sigma}}_0 = \boldsymbol{\Sigma}(\hat{\sigma}^2_0, \hat{\rho}_0)$. The score test statistic is therefore

$$Z_{\text{score}} = \frac{\boldsymbol{D}^{\top}\hat{\boldsymbol{\Sigma}}_0^{-1}\boldsymbol{1}}{\sqrt{\boldsymbol{1}^{\top}\hat{\boldsymbol{\Sigma}}_0^{-1}\boldsymbol{1}}} = \frac{\overline{D}}{\sqrt{\hat{\sigma}_0^2\left(\frac{1}{2J}+\frac{J-1}{J}\hat{\rho}_0\right)}} \qquad \text{(S12.28)}$$

which is asymptotically standard Gaussian under the null hypothesis. The two-sided p-value is given by $p = 2\big(1 - \Phi(|Z_{\text{score}}|)\big)$, where $\Phi(\cdot)$ denote the standard normal cumulative distribution function.

S12.7 Likelihood ratio test (LRT) and signed-root transformation
As a final statistical test, we compare the maximum log likelihood under the null hypothesis with the maximum unrestricted log likelihood:

$$X_{\text{LRT}}^2 = -2\,[\ell(0, \tilde{\sigma}_0^2, \tilde{\rho}_0) - \ell(\tilde{\mu}, \tilde{\sigma}^2, \tilde{\rho})] \qquad \text{(S12.29)}$$

where $(\tilde{\sigma}_0^2, \tilde{\rho}_0)$ maximizes the log likelihood assuming the null is true (same as $(\hat{\sigma}_0^2, \hat{\rho}_0)$ from the score test in Supplementary Methods S12.6), and $(\tilde{\mu}, \tilde{\sigma}^2, \tilde{\rho})$ maximizes the unrestricted log likelihood (same as $(\overline{D}, \hat{\sigma}_{\text{ML}}^2, \hat{\rho}_{\text{ML}})$ from the maximum likelihood test in Supplementary Methods S12.4).

Under the null hypothesis, $X_{\text{LRT}}^2$ follows the chi square distribution with one degree of freedom. The test statistic is converted to a signed-root form for interpretability:

$$Z_{\text{LRT}} = \text{sign}(\tilde{\mu})\sqrt{X_{\text{LRT}}^2}\ . \qquad \text{(S12.30)}$$

The two-sided p-value is given by $p = 2\big(1 - \Phi(|Z_{\text{LRT}}|)\big)$, where $\Phi(\cdot)$ denotes the standard normal cumulative distribution function.

S12.8 SHARP tests toy example selection
To select among the variants of SHARP, we sampled $\boldsymbol{D} \sim N(\mu\boldsymbol{1}, \boldsymbol{\Sigma}(\sigma^2, \rho))$ with $\mu = 0, \sigma^2 = 1$ while varying $\rho$ from 0.01 to 0.49. $J$ = 300, so $\boldsymbol{D}$ is a vector of length 600. The whole procedure was repeated independently 100 times to compute a FPR for each value of $\rho$. The score test showed the best control of FPR (Supplementary Fig. 12) and was therefore used for the rest of the manuscript.

S12.9 Split-HAlf (SHA) test
As a variant of the SHARP test, we can also randomly divide the dataset into two disjoint halves A and B once (as opposed to $J$ times like the SHARP test). For each machine learning model, we then perform K-fold cross-validation in subsets A and B separately. The data split for the K-fold cross-validation is identical for both models, so there is fold-level correspondence between the two models. Denote the fold-level model performance differences $D_{Ak}$ and $D_{Bk}$ respectively, for $k = 1, \dots, K$. In this version of the procedure, there is no replication. This results in two length-$K$ vectors $\boldsymbol{D}_A = [D_{A1}, \dots, D_{AK}]^{\top}$ and $\boldsymbol{D}_B = [D_{B1}, \dots, D_{BK}]^{\top}$.

Let $\boldsymbol{D} = [\boldsymbol{D}_A^{\top}, \boldsymbol{D}_B^{\top}]^{\top} = [D_{A1}, \dots, D_{AK},\ D_{B1}, \dots, D_{BK}]^{\top}$, so $\boldsymbol{D}$ is a column vector of length $2K$. We denote $\text{Var}(D_{Ak}) = \text{Var}(D_{Bk}) = \sigma^2$. Because of the disjoint subsets A and B, $\text{Corr}(D_{Ak}, D_{Bk}) = 0$ but also for any pair of folds $k\ \&\ k'$, $\text{Corr}(D_{Ak}, D_{Bk'}) = 0$. On the other hand, within subsets, $\mathrm{Corr}(D_{Ak}, D_{Ak'}) = \mathrm{Corr}(D_{Bk}, D_{Bk'}) = \rho$. Therefore, the covariance matrix of $\boldsymbol{D}$ can be written as

$$\boldsymbol{\Sigma}(\sigma^2, \rho) = \sigma^2 \begin{bmatrix} \boldsymbol{M} & \boldsymbol{0} \\ \boldsymbol{0} & \boldsymbol{M} \end{bmatrix} \tag{S12.31}$$

where $\boldsymbol{M} \in \mathbb{R}^{K \times K}$ has ones on the diagonal and $\rho$ in all off-diagonal entries,

$$M_{jk} = \begin{cases} 1, j = k \\ \rho, j \neq k \end{cases} \text{ or } \boldsymbol{M} = \begin{bmatrix} 1 & \rho & \rho \\ \rho & \ddots & \rho \\ \rho & \rho & 1 \end{bmatrix} \tag{S12.32}$$

and $\boldsymbol{0} \in \mathbb{R}^{K \times K}$ is the zeros matrix. Similar to the SHARP test, we can use various approaches to estimate $\sigma^2$ and $\rho$, followed by a Wald test. A likelihood ratio test or score test can also be utilized. The SHA test likely has worse statistical power than the SHARP test, so we did not consider the SHA test in the current study.

### Supplemental Results

#### Meta-analysis: prevalence

In our meta-analysis, we encountered a study using a permutation test that was invalid for reasons unrelated to between-fold correlation. In this study, the test statistic was the performance difference between models evaluated using 10-fold cross-validation. The labels were then permuted 1000 times. For each permutation, both models were re-evaluated using the same 10-fold cross-validation procedure, thereby constructing a null distribution of performance differences. The p-value was obtained by comparing the observed performance difference to this permutation-based null distribution. However, this procedure did not actually test the null hypothesis of equal predictive performance between models. By permuting labels, the relationship between features and outcomes was destroyed, so the resulting null distribution reflected a no-signal scenario rather than the case where two models performed equally well. We note that this is not an infrequent mistake in the literature. Because this error fell outside the scope of our meta-analysis, we excluded this study from our meta-analysis, so we ended up with 184 studies.

In Fig. 2b, we reported that six studies utilized statistical tests that accounted for between-fold correlation. Four of them used the corrected resampled t-test (Nadeau and Bengio, 2003), while one of them used the 5×2 t-test (Dietterich, 1998). The final study utilized the following heuristic: a model was declared superior to another model only if the lower bound of its reported performance range (mean − 2 × SD across the 10 cross-validation folds) exceeded the other model's mean performance. Because the variability term used was the standard deviation rather than the standard error ($SD/\sqrt{10}$), this criterion can be seen as a variant of the empirical test of differences, which implicitly accounts for between-fold correlation (Supplementary Methods S5). As such, we consider it to account for between-fold correlation by avoiding the independence-based $\sqrt{10}$ shrinkage of variability.

In Fig. 2c, we categorized 11 studies that employed statistical tests assuming fold independence under "Others." Of these, seven studies used ANOVA (Fisher, 1992), a generalization of the naïve resampled paired t-test for comparing more than two models. One study each employed the Quade test (Quade, 1979), the Kolmogorov-Smirnov test (Massey, 1951), the U-test (Harrell, 2001) and the Tukey-Kramer test (Kramer, 1956).

#### Meta-analysis: re-analysis

In the main text, we sought to contextualize how far the median corrected p-value $p = 0.17$ lies from statistical significance. Under a two-sided standard-normal reference, $p = 0.17$ corresponds to $|z| = 1.37$, compared with $|z| = 1.96$ at $p = 0.05$. On average, reaching significance would therefore require a 43% larger effect size [1.96/1.37 = 1.43]. To make this gap intuitive to readers unfamiliar with machine learning, we also provide a benchmark based on a conventional analysis of independent observations. Holding the population mean and observation-level variance constant, the variance of the sample mean scales as $1/N$. Reaching significance would then require twice as many independent observations [$(1.96/1.37)^2 \approx 2.05$].

This is an illustrative benchmark, not a sample-size calculation for cross-validation studies. Adding folds or repetitions does not create independent observations. Even if the original sample size were doubled and the mean difference remained unchanged after rerunning cross-validation, there is no general way to predict how much the variance of the mean difference would decrease – and therefore whether significance would be regained. Thus, this illustration is intended to convey the gap between p = 0.17 and p = 0.05, not to estimate the sample size increase required for the studies in our re-analysis.

In the main text, we estimated that 32% of studies in the broader literature contain an abstract-level supported by a spurious comparison. A second route through the corpus reaches a similar value: 173 of the 184 studies were at risk, 47 of the 68 studies permitting re-analysis had a spurious comparison, and in 22 of the 47 studies had an abstract-level claim supported by a spurious comparison. Therefore, a different route to computing the prevalence is (173/184) × (47/68) × (22/47) = 30%. The two routes are not independent, but they decompose the corpus differently and they agree because the proportion of studies containing a model-comparison claim in the abstract was nearly identical among the 68 re-analyzed studies (37/68 = 54%) and the remaining 105 at-risk studies (63/105 = 60%).

### Supplemental Tables

**Table S1. Criteria for evaluating journal statistical rigor & methodological guidance**

| Category | Score 0 | Score 1 | Score 2 |
|---|---|---|---|
| Code availability policy | No mention in guidelines/policies; no encouragement or requirement to share code/algorithms. | Code sharing encouraged or optional but not required for publication. | Code sharing is mandatory and a condition for publication. |
| Data availability policy | No mention in guidelines/policies; no encouragement or requirement to share data. | Data sharing (any type) encouraged or optional but not required. | Data sharing is mandatory and a condition for publication. |
| Code for peer review policy | No mention of providing code specifically for peer review. | Code submission for review encouraged but not mandatory. | Code submission required for peer review. |
| Availability statement requirement | No request or mention of a data/code/materials availability statement. | Availability statement encouraged but not mandatory. | Formal availability statement required for submission/publication. |
| Statistical test guidance rigor (11 items*) | None of the 11 statistical reporting items mentioned. | 1–3 of the 11 items mentioned. | More than 3 of the 11 items mentioned. |
| Statistical review policy | No mention or requirement of statistical review in reviewer guidance or peer review policy. | Statistical review acknowledged and optional. | Statistical review required for specified article types (e.g., clinical trials, ML, meta-analysis, high-dimensional data). |
| Reporting standards/checklists | No scientific reporting checklist mentioned; no requirement or encouragement of external reporting guidelines. | Reporting checklists encouraged but not required. | Completion of one or more scientific reporting checklists required for submission, review, or acceptance. |

*Statistical Test Guidance Rigor – 11 Items Considered:
1. Report test name; 2. Report p-value with ns/s; 3. Report sample size per group;
4. State one- or two-tailed; 5. Justify test/assumption checks; 6. State alpha level;
7. Report degrees of freedom; 8. Carry out multiple testing correction;
9. Include clinical/practical significance; 10. Report confidence intervals;
11. Explain methods for constructing confidence intervals.

**Table S2. Web of Science subject categories represented among the 2,400 screened studies**

| **Web of Science Core Collection Subject Categories** | | |
|---|---|---|
| *Behavioral Sciences* | *Gastroenterology & Hepatology* | Parasitology |
| *Biochemical Research Methods* | *Genetics & Heredity* | Pediatrics |
| *Biochemistry & Molecular Biology* | *Geriatrics & Gerontology* | Peripheral Vascular Disease |
| *Biotechnology & Applied Microbiology* | Green & Sustainable Science & Technology | *Pharmacology & Pharmacy* |
| *Cardiac & Cardiovascular Systems* | Hematology | Plant Sciences |
| *Cell & Tissue Engineering* | *Immunology* | *Psychiatry* |
| *Cell Biology* | Infectious Diseases | Psychology |
| *Chemistry, Multidisciplinary* | Materials Science, Biomaterials | Psychology, Biological |
| Clinical Neurology | Medical Ethics | Psychology, Experimental |
| *Computer Science, Artificial Intelligence* | *Medical Informatics* | Psychology, Multidisciplinary |
| *Computer Science, Interdisciplinary Applications* | Medicine, General & Internal | *Public, Environmental & Occupational Health* |
| Critical Care Medicine | *Medicine, Research & Experimental* | *Radiology, Nuclear Medicine & Medical Imaging* |
| Dentistry, Oral Surgery & Medicine | *Microbiology* | Remote Sensing |
| *Endocrinology & Metabolism* | Mycology | *Respiratory System* |
| *Engineering, Biomedical* | *Neurosciences* | Rheumatology |
| *Engineering, Electrical & Electronic* | *Oncology* | Surgery |
| Engineering, Environmental | *Ophthalmology* | Urology & Nephrology |
| *Environmental Sciences* | *Optics* | Virology |
| Food Science & Technology | | |

Categories shown in *italics* are the 30 categories represented among the 184 studies in the final set.

### Supplemental Figures

#### Prevalence of invalid tests across journal metrics & open science practice

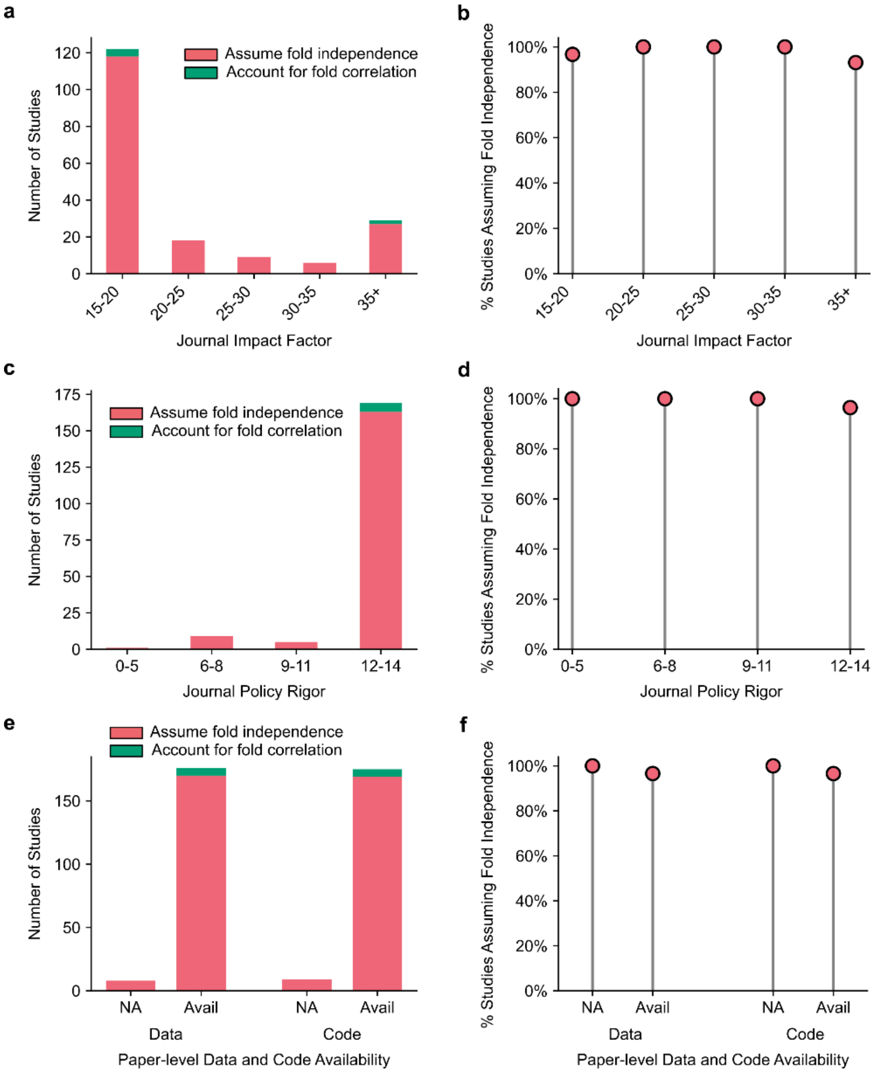


**Supplementary Fig. 1. Neglect of between-fold correlation persists across impact factor, journal policies for scientific rigor and open science practices. a.** Distribution of studies by journal impact factor. **b.** Proportion of studies assuming fold independence (i.e., ignoring between-fold correlation) by impact factor bins, calculated as the number of studies using statistical tests assuming fold independence divided by the total number of studies in each bin. No trend was detected (permutation test $p = 0.59$). **c.** Distribution of studies by journal policy rigor score, ranging from 0 (least rigorous) to 14 (most rigorous). **d.** Proportion of studies assuming fold independence by journal policy rigor score. Increasing rigor did not reduce the use of invalid statistical tests (permutation test $p = 0.89$). **e.** Number of studies assuming fold independence stratified by data and code availability. Most studies provided data, code or both. **f.** Proportion of studies assuming fold independence stratified by data and code availability. The proportion was not significantly associated with data or code availability (permutation test $p = 1.00$ for both).

**DeLong test**

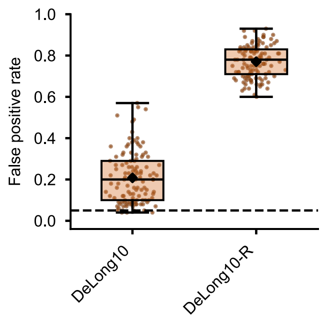


**Supplementary Fig. 2. False positive rate (FPR) for the DeLong test with 10-fold cross-validation repeated once ("DeLong10") or repeated 30 times ("DeLong10-R").** The DeLong test is only applicable to binary classification, so the current analyses were based on the Covertype dataset. Each boxplot comprised 120 data points, corresponding to 120 scenarios (4 sample sizes × 3 hyperparameter schemes × 10 noise levels). The black dot indicates the mean FPR across 120 scenarios. The DeLong test exhibited FPRs of 21% and 77% for 10-fold cross-validation with one repetition and 30 repetitions, respectively. For comparison, the paired t-test on fold-averaged statistics in the same dataset yielded FPRs of 13% and 72% (Supplementary Fig. 3c).

#### FPR across datasets

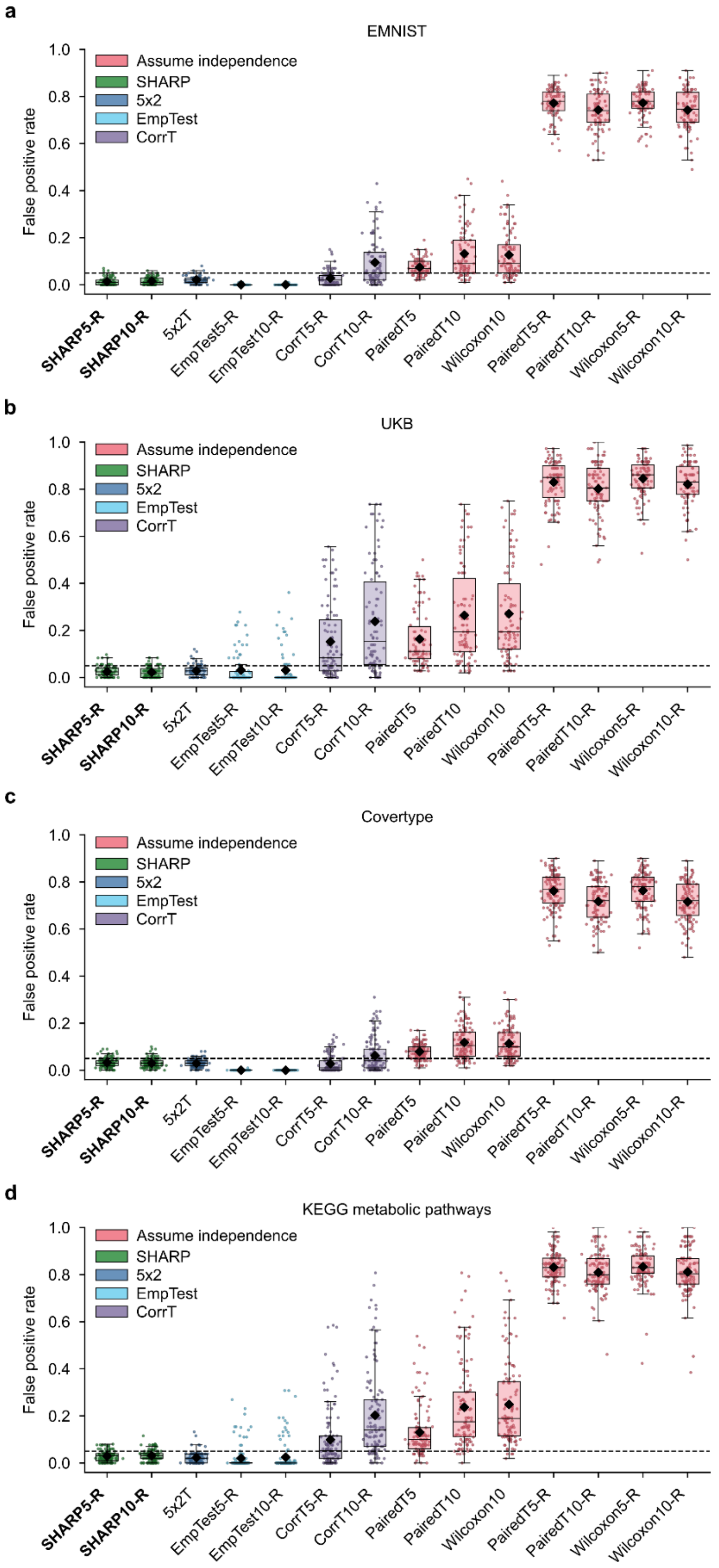


**Supplementary Fig. 3. False positive rates (FPR) of various statistical tests across 420 scenarios broken down by datasets. a.** Boxplots of FPRs in the EMNIST dataset. Each boxplot comprises 90 data points, corresponding to 90 scenarios (3 sample sizes × 10 noise levels × 3 hyperparameter schemes). **b.** Boxplots of FPRs in the UKB dataset. Each boxplot comprises 90 datapoints, corresponding to 90 scenarios (3 sample sizes × 10 noise levels × 3 hyperparameter schemes). **c.** Boxplots of FPRs in the Covertype dataset. Each boxplot comprises 120 datapoints, corresponding to 120 scenarios (4 sample sizes × 10 noise levels × 3 hyperparameter schemes). **d.** Boxplots of FPRs in the metabolic dataset. Each boxplot comprises 120 datapoints, corresponding to 120 scenarios (4 sample sizes × 10 noise levels × 3 hyperparameter schemes). **Test naming conventions.** Numeric suffixes "5" and "10" denote 5-fold and 10-fold cross-validation. The "-R" suffix indicates repeated cross-validation: 5-fold repeated 60 times or 10-fold repeated 30 times, yielding 5 × 60 = 10 × 30 = 300 fold-level statistics. For example, "CorrT5-R" denotes the corrected resampled t-test evaluated under 60 repetitions of 5-fold cross-validation, while "PairedT10" denotes the naïve paired t-test under a single run of 10-fold cross-validation. "SHARP5-R" denotes the split-half procedure repeated 60 times with 5-fold cross-validation within each half; "SHARP10-R" denotes 30 repetitions with 10-fold cross-validation within each half. The number of repetitions was chosen such that the FPR of tests accounting for between-fold correlation had stabilized; tests that assume fold independence do not stabilize, with FPR increasing toward 1 as repetitions grow (Fig. 5d). "CorrT" denotes the corrected resampled t-test (Nadeau & Bengio, 2003). “EmpTest” denotes the empirical test of differences (Parkes et al., 2021a). For details on all tests, see Methods “Existing statistical tests for cross-validation” and Methods “SHARP test”.

**FPR across sample sizes**

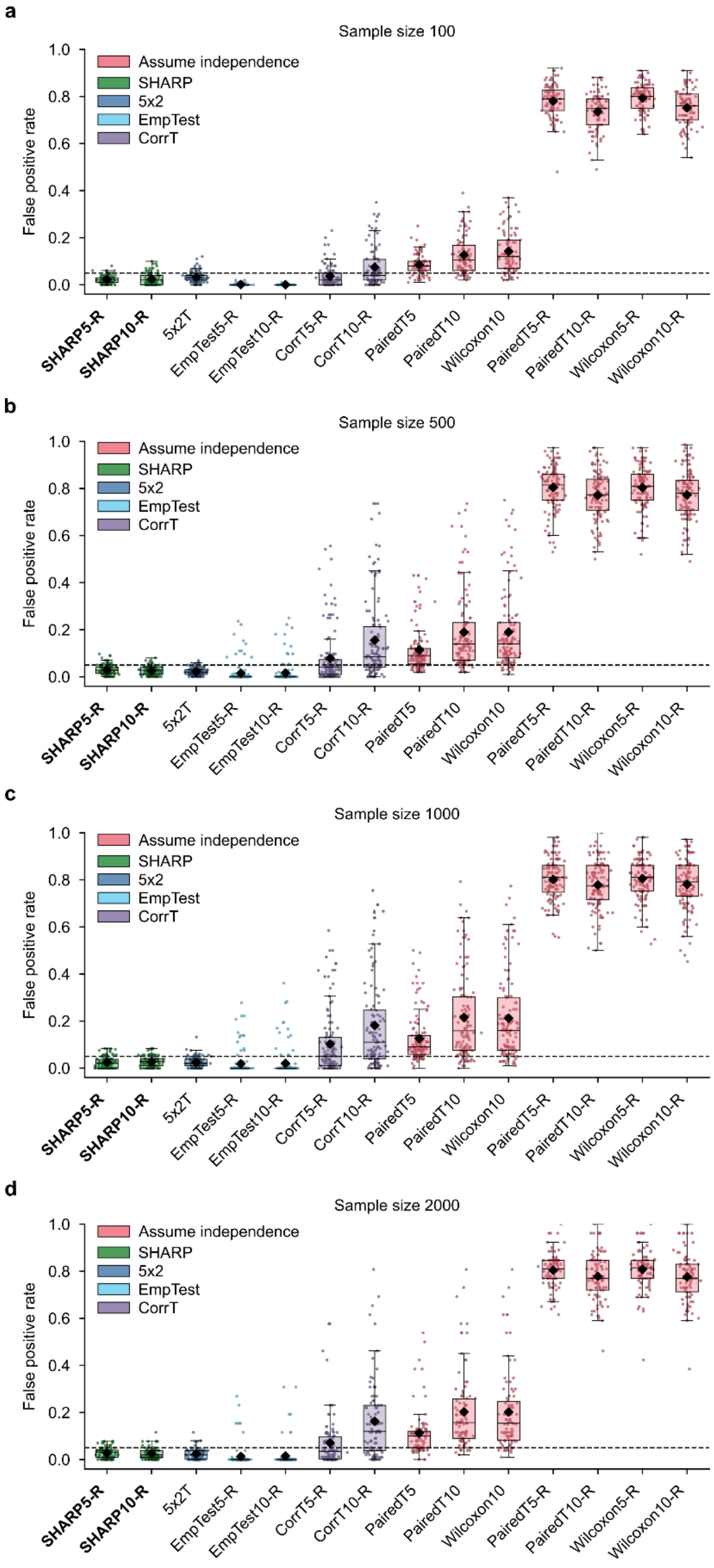


**Supplementary Fig. 4. False positive rates (FPRs) of various statistical tests across 420 scenarios broken down by sample sizes. a.** Boxplots of FPRs for simulations involving sample size of 100. Each boxplot comprises 90 datapoints, corresponding to 90 scenarios (3 datasets × 10 noise levels × 3 hyperparameter schemes). **b.** Boxplots of FPRs for simulations involving sample size of 500. Each boxplot comprises 120 datapoints, corresponding to 120 scenarios (4 datasets × 10 noise levels × 3 hyperparameter schemes). **c.** Boxplots of FPRs for simulations involving sample size of 1000. Each boxplot comprises 120 datapoints, corresponding to 120 scenarios (4 datasets × 10 noise levels × 3 hyperparameter schemes). **d.** Boxplots of FPRs for simulations involving sample size of 2000. Each boxplot comprises 90 datapoints, corresponding to 90 scenarios (3 datasets × 10 noise levels × 3 hyperparameter schemes). **Test naming conventions.** Numeric suffixes "5" and "10" denote 5-fold and 10-fold cross-validation. The "-R" suffix indicates repeated cross-validation: 5-fold repeated 60 times or 10-fold repeated 30 times, yielding 5 × 60 = 10 × 30 = 300 fold-level statistics. For example, "CorrT5-R" denotes the corrected resampled t-test evaluated under 60 repetitions of 5-fold cross-validation, while "PairedT10" denotes the naïve paired t-test under a single run of 10-fold cross-validation. "SHARP5-R" denotes the split-half procedure repeated 60 times with 5-fold cross-validation within each half; "SHARP10-R" denotes 30 repetitions with 10-fold cross-validation within each half. The number of repetitions was chosen such that the FPR of tests accounting for between-fold correlation had stabilized; tests that assume fold independence do not stabilize, with FPR increasing toward 1 as repetitions grow (Fig. 5d). "CorrT" denotes the corrected resampled t-test (Nadeau & Bengio, 2003). “EmpTest” denotes the empirical test of differences (Parkes et al., 2021a). For details on all tests, see Methods “Existing statistical tests for cross-validation” and Methods “SHARP test”.

#### FPR for two-algorithm scheme

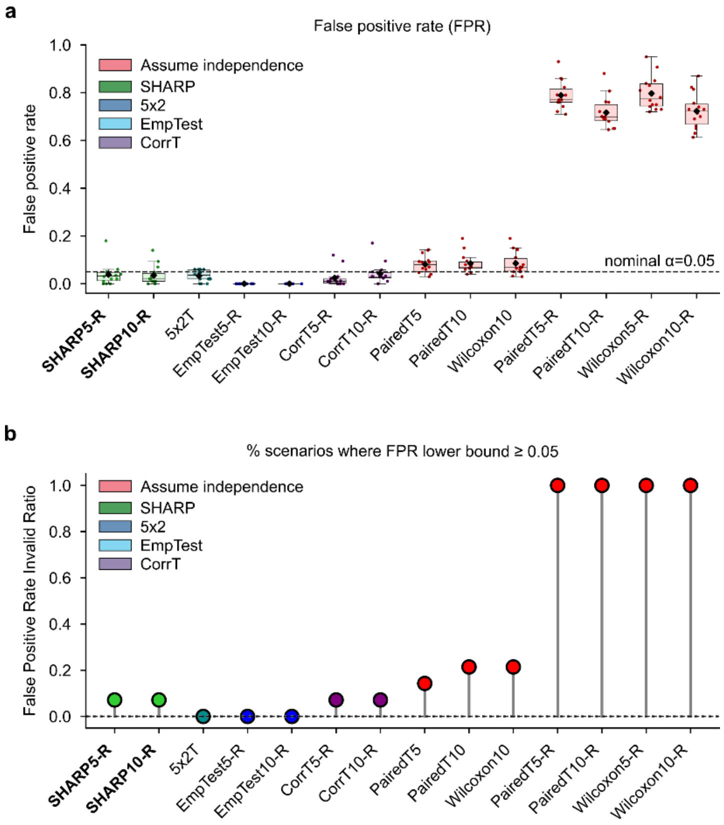


**Supplementary Fig. 5. False positive rates (FPRs) with the two-algorithm simulation scheme (Supplementary Methods S11). a.** Boxplots of FPR for each statistical test. Each boxplot contains 14 values, corresponding to the 14 scenarios. The black dot indicates the mean FPR across 14 scenarios. The dashed horizontal line marks the nominal FPR of 0.05. **b.** FPR inflation rate: the percentage of the 14 scenarios in which the lower bound of the 95% confidence interval for FPR exceeded 0.05, indicating inadequate control of the Type I error rate. **Test naming conventions.** Numeric suffixes "5" and "10" denote 5-fold and 10-fold cross-validation. The "-R" suffix indicates repeated cross-validation: 5-fold repeated 60 times or 10-fold repeated 30 times, yielding 5 × 60 = 10 × 30 = 300 fold-level statistics. For example, "CorrT5-R" denotes the corrected resampled t-test evaluated under 60 repetitions of 5-fold cross-validation, while "PairedT10" denotes the naïve paired t-test under a single run of 10-fold cross-validation. "SHARP5-R" denotes the split-half procedure repeated 60 times with 5-fold cross-validation within each half; "SHARP10-R" denotes 30 repetitions with 10-fold cross-validation within each half. The number of repetitions was chosen such that the FPR of tests accounting for between-fold correlation had stabilized; tests that assume fold independence do not stabilize, with FPR increasing toward 1 as repetitions grow (Fig. 5d). "CorrT" denotes the corrected resampled t-test (Nadeau & Bengio, 2003). “EmpTest” denotes the empirical test of differences (Parkes et al., 2021a). For details on all tests, see Methods “Existing statistical tests for cross-validation” and Methods “SHARP test”. Conclusions were the same as Fig. 6, except that the corrected resampled t-test reliably controlled FPR in this simulation.

**FPR for more tests**

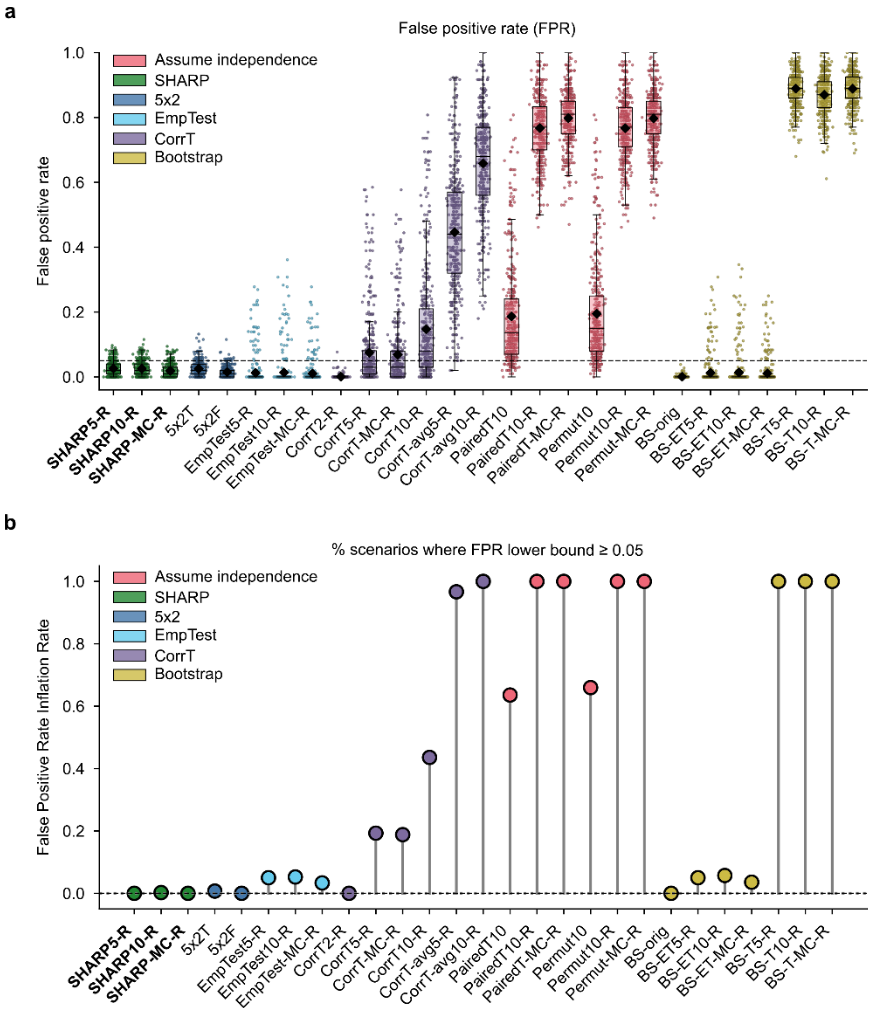


**Supplementary Fig. 6. False positive rate (FPR) across 420 simulation scenarios for an extended set of tests. a.** Boxplot of FPRs for each statistical test, with one value per scenario (n = 420). Black dots mark the mean FPR across scenarios. The dashed horizontal line marks the nominal level of 0.05. **b.** FPR inflation rate: the percentage of the 420 scenarios in which the lower bound of the 95% confidence interval for FPR exceeded 0.05, indicating inadequate control of the Type I error rate. **Test naming conventions.** Numeric suffixes "5" and "10" denote 5-fold and 10-fold cross-validation. The "-R" suffix indicates repeated cross-validation, with the number of repetitions chosen such that the FPR of tests accounting for between-fold correlation had stabilized. Under standard cross-validation, "-R" corresponds to 5-fold repeated 60 times or 10-fold repeated 30 times, yielding 5 × 60 = 10 × 30 = 300 fold-level statistics. For example, "EmpTest5-R" denotes the empirical test of differences under 60 repetitions of 5-fold cross-validation, while "PairedT10" denotes the naïve paired t-test under a single run of 10-fold cross-validation. For SHARP variants, each repetition produces one pair of statistics — one per half — so the number of pairs equals the number of repetitions: "SHARP5-R" denotes the split-half procedure repeated 60 times with 5-fold cross-validation within each half (yielding 60 pairs), and "SHARP10-R" denotes 30 repetitions with 10-fold cross-validation within each half (yielding 30 pairs). The "MC" suffix denotes Monte Carlo cross-validation, where the dataset is randomly split into 80% training and 20% test sets. Under Monte Carlo cross-validation, "-R" corresponds to 300 repetitions, again yielding 300 fold-level statistics; for "SHARP-MC-R", a single Monte Carlo split is performed within each half and the split-half procedure is repeated 300 times (yielding 300 pairs). "CorrT" denotes the corrected resampled t-test (Nadeau & Bengio, 2003); "CorrT-avg" denotes a misapplied variant (Supplementary Methods S8). "EmpTest" denotes the empirical test of differences (Parkes et al., 2021a). "Permut" denotes the paired permutation test (Supplementary Methods S5.3). "BS" denotes bootstrap with three variants (Supplementary Methods S5.8): bootstrap-orig ("BS-orig"), bootstrap-empirical-test-of-differences ("BS-ET") and bootstrap-t-test ("BS-T"). Two bootstrap variants ("BS-orig" and "BS-ET") reliably controlled FPR, but the third bootstrap variant ("BS-T") exhibited high FPR. For details about the various tests, see Methods "Existing statistical tests for cross-validation" and Methods "SHARP test".

**Power Simulation Scheme (Noisy-Model)**

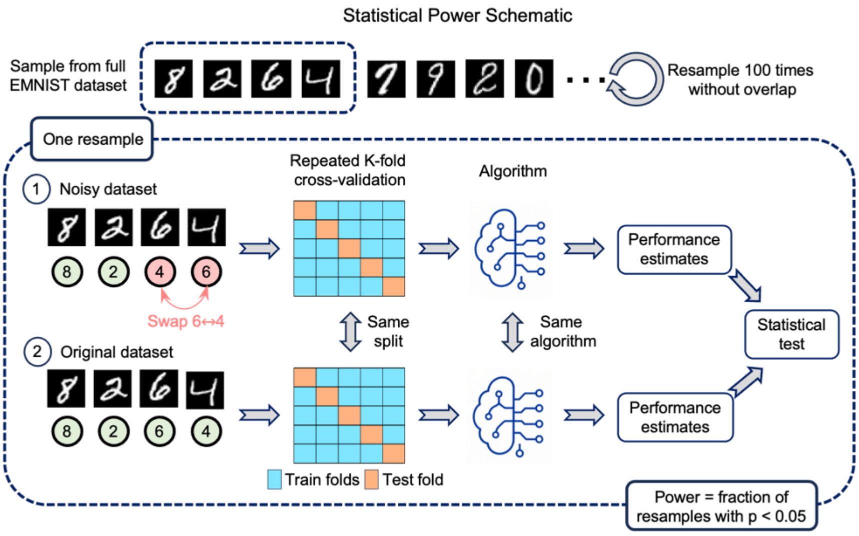


**Supplementary Fig. 7. Statistical power simulation scheme illustrated for EMNIST.** We randomly sampled a dataset (e.g., N = 1,000) from a full dataset, then generated a noisy version by permuting the target labels (ground-truth digits) for a fixed percentage of samples (e.g., 10%). For example, the noisy dataset (above) has the labels for digits 6 and 4 swapped. Using identical cross-validation splits and the same machine learning algorithm on the clean and noisy datasets, we obtained pairs of cross-validated performance estimates — e.g., 300 pairs for 10-fold cross-validation repeated 30 times. A statistical test was applied to the paired vector to obtain a p-value. Because only one dataset was corrupted, the model trained on the clean data had higher expected predictive performance by construction, so a sensitive test should reject the null hypothesis. This procedure was repeated up to 100 times using non-overlapping subsamples, with power computed as the percentage of repetitions in which the null was rejected. In total, there are 420 scenarios.

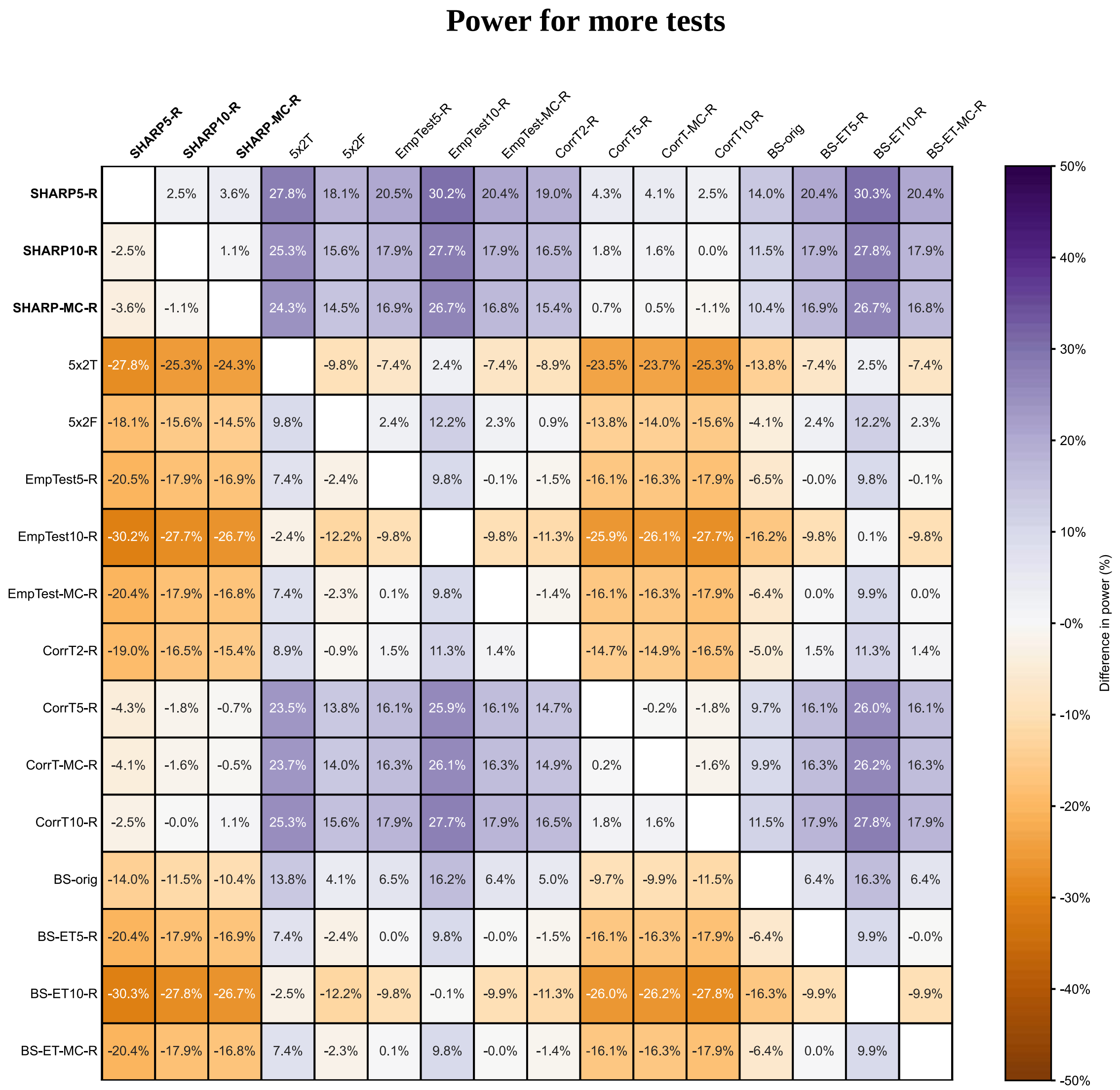


**Supplementary Fig. 8. Comparison of statistical power across 420 scenarios for an extended set of tests.** Difference in power between pairs of statistical tests, computed as the power of the statistical test (on the row) minus the power of the statistical test (on the column), averaged across 420 scenarios. A purple cell indicates that the test on the row achieved higher power than the test on the column; an orange cell indicates the opposite. The set of statistical tests was the same as Supplementary Fig. 6, except that only valid statistical tests are included. The SHARP test with repeated 5-fold cross-validation (SHARP5-R) demonstrated the best power, as indicated by the entirely purple SHARP5-R row. **Test naming conventions.** Numeric suffixes "5" and "10" denote 5-fold and 10-fold cross-validation. The "-R" suffix indicates repeated cross-validation, with the number of repetitions chosen such that the power of tests accounting for between-fold correlation had stabilized. Under standard cross-validation, "-R" corresponds to 5-fold repeated 60 times or 10-fold repeated 30 times, yielding 5 × 60 = 10 × 30 = 300 fold-level statistics. For example, "EmpTest5-R" denotes the empirical test of differences under 60 repetitions of 5-fold cross-validation. For SHARP variants, each repetition produces one pair of statistics — one per half — so the number of pairs equals the number of repetitions: "SHARP5-R" denotes the split-half procedure repeated 60 times with 5-fold cross-validation within each half (yielding 60 pairs), and "SHARP10-R" denotes 30 repetitions with 10-fold cross-validation within each half (yielding 30 pairs). The "MC" suffix denotes Monte Carlo cross-validation, where the dataset is randomly split into 80% training and 20% test sets. Under Monte Carlo cross-validation, "-R" corresponds to 300 repetitions, again yielding 300 fold-level statistics; for "SHARP-MC-R", a single Monte Carlo split is performed within each half and the split-half procedure is repeated 300 times (yielding 300 pairs). "CorrT" denotes the corrected resampled t-test (Nadeau & Bengio, 2003); "EmpTest" denotes the empirical test of differences (Parkes et al., 2021a). "BS" denotes bootstrap with two variants (Supplementary Methods S5.8): bootstrap-orig ("BS-orig") and bootstrap-empirical-test-of-differences ("BS-ET"). For details about the various tests, see Methods “Existing statistical tests for cross-validation” and Methods “SHARP test”.

**Power for two-algorithm scheme**

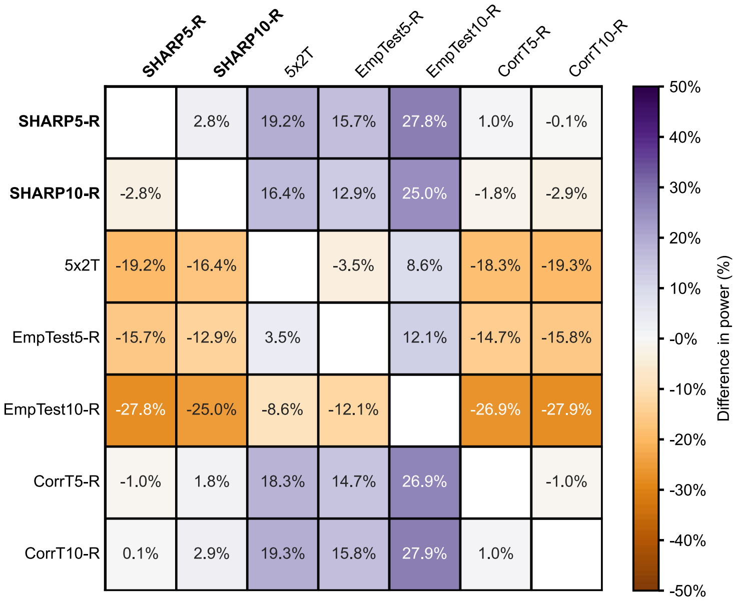


**Supplementary Fig. 9. Comparison of statistical power of statistical tests with the two-algorithm simulation scheme (Supplementary Methods S11).** Difference in power between pairs of statistical tests, computed as the power of the statistical test (on the row) minus the power of the statistical test (on the column), averaged across 14 scenarios. A purple cell indicates that the test on the row achieved higher power than the test on the column; an orange cell indicates the opposite. The set of statistical tests was the same as Fig. 7a. The SHARP test with repeated 5-fold cross-validation (SHARP5-R) and the corrected resampled t-test with repeated 10-fold cross-validation (CorrT10-R) had the best power among the tests that achieved nominal FPR control.

#### CI coverage for more tests

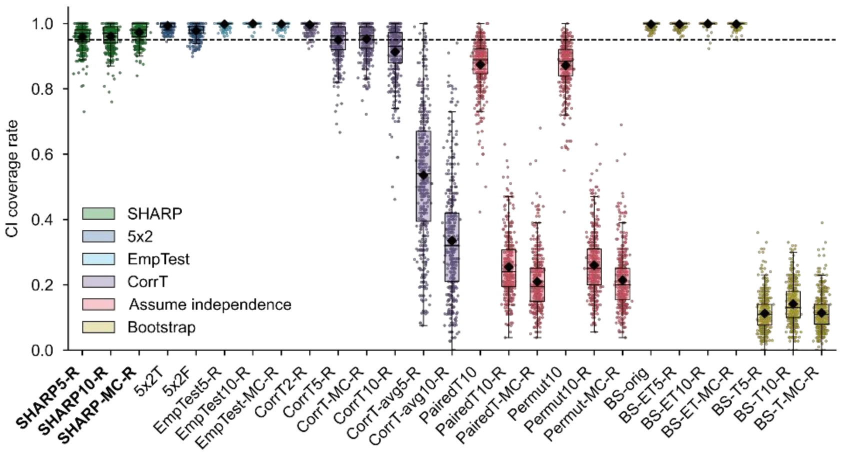


**Supplementary Fig. 10. Comparison of 95% confidence interval (CI) coverage rate across 420 scenarios for an extended set of tests.** The CI coverage rate of a scenario is defined as the percentage of non-overlapping subsamples in which the (estimated) true performance difference between models fell inside the 95% CI of a given test. For a well-calibrated test, the coverage rate should be exactly 95% (black dashed line). Each boxplot comprises 420 data points, each representing the CI coverage rate of one scenario. The black dot indicates the mean coverage rate across 420 scenarios. Tests that assumed fold independence (red boxplots) had overly narrow CIs, so their coverage rates were much lower than 95%. **Test naming conventions.** Numeric suffixes "5" and "10" denote 5-fold and 10-fold cross-validation. The "-R" suffix indicates repeated cross-validation, with the number of repetitions chosen such that the FPR of tests accounting for between-fold correlation had stabilized. Under standard cross-validation, "-R" corresponds to 5-fold repeated 60 times or 10-fold repeated 30 times, yielding 5 × 60 = 10 × 30 = 300 fold-level statistics. For example, "EmpTest5-R" denotes the empirical test of differences under 60 repetitions of 5-fold cross-validation, while "PairedT10" denotes the naïve paired t-test under a single run of 10-fold cross-validation. For SHARP variants, each repetition produces one pair of statistics — one per half — so the number of pairs equals the number of repetitions: "SHARP5-R" denotes the split-half procedure repeated 60 times with 5-fold cross-validation within each half (yielding 60 pairs), and "SHARP10-R" denotes 30 repetitions with 10-fold cross-validation within each half (yielding 30 pairs). The "MC" suffix denotes Monte Carlo cross-validation, where the dataset is randomly split into 80% training and 20% test sets. Under Monte Carlo cross-validation, "-R" corresponds to 300 repetitions, again yielding 300 fold-level statistics; for "SHARP-MC-R", a single Monte Carlo split is performed within each half and the split-half procedure is repeated 300 times (yielding 300 pairs). "CorrT" denotes the corrected resampled t-test (Nadeau & Bengio, 2003); "CorrT-avg" denotes a misapplied variant (Supplementary Methods S8). "EmpTest" denotes the empirical test of differences (Parkes et al., 2021a). "Permut" denotes the paired permutation test (Supplementary Methods S5.3). "BS" denotes bootstrap with three variants (Supplementary Methods S5.8): bootstrap-orig (“BS-orig”), bootstrap-empirical-test-of-differences (“BS-ET”) and bootstrap-t-test (“BS-T”). Two bootstrap variants (“BS-orig” and “BS-ET”) reliably controlled FPR, but the third bootstrap variant (“BS-T”) exhibited high FPR. For details about the various tests, see Methods “Existing statistical tests for cross-validation” and Methods “SHARP test”.

**CI coverage (two-algorithm)**

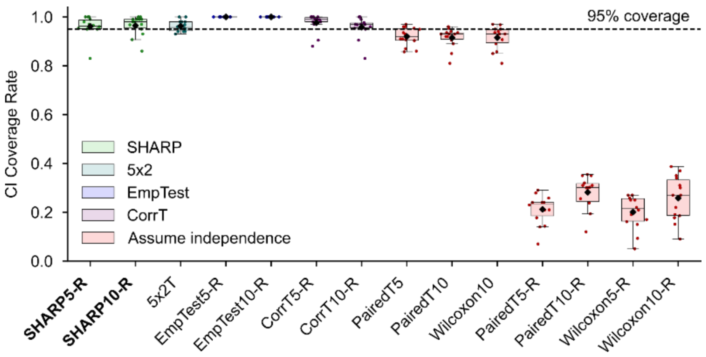


**Supplementary Fig. 11. Comparison of 95% confidence interval (CI) coverage rate across statistical tests under the two-algorithm simulation scheme (Supplementary Methods S11)**. The CI coverage rate of a scenario is defined as the percentage of non-overlapping subsamples in which the (estimated) true performance difference between models fell inside the 95% CI of a given test. For a well-calibrated test, the coverage rate should be exactly 95% (black dashed line). Each boxplot comprises 14 data points, each representing the CI coverage rate of one scenario. The black dot indicates the mean coverage rate across 14 scenarios. Tests that assumed fold independence (red boxplots) had overly narrow CIs, so their coverage rates were much lower than 95%. The test naming conventions are the same as Supplementary Fig. 5.

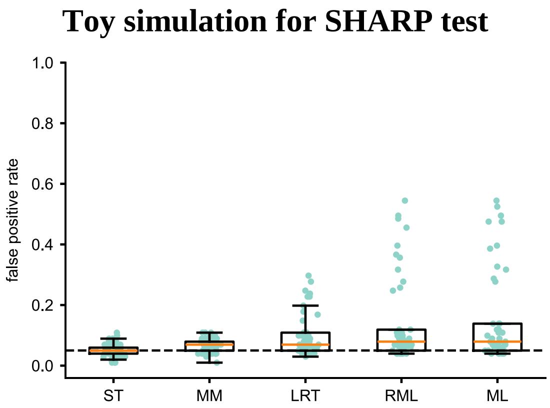


**Supplementary Fig. 12. Toy simulation to select the best variant of the SHARP test**. We randomly draw performance differences of two methods from a Gaussian distribution, $\boldsymbol{D} \sim N(\mu\mathbf{1}, \mathbf{\Sigma}(\sigma^2, \rho))$ with $\mu = 0, \sigma = 1$, while $\rho$ varies from 0.01 to 0.49. There are in total 49 dots in each box corresponding to the 49 values of $\rho$. Score test shows the best false positive control. Test abbreviations: ST: Score Test; MM: Method of Moments; LRT: Likelihood Ratio Test; RML: Restricted Maximum Likelihood Test; ML: Maximum Likelihood Test. The score test performed the best, so was used throughout the manuscript.

### References


Alfaro-Almagro, F., Jenkinson, M., Bangerter, N. K., Andersson, J. L. R., Griffanti, L., Douaud, G., Sotiropoulos, S. N., Jbabdi, S., Hernandez-Fernandez, M., Vallée, E., Vidaurre, D., Webster, M., McCarthy, P., Rorden, C., Daducci, A., Alexander, D. C., Zhang, H., Dragonu, I., Matthews, P. M., … Smith, S. M. (2018). Image processing and quality control for the first 10,000 brain imaging datasets from UK Biobank. *NeuroImage*, *166*, 400–424. https://doi.org/10.1016/j.neuroimage.2017.10.034

Ali, M. (2020). *PyCaret: An open source, low-code machine learning library in Python* (Version 1.0) [Computer software]. https://www.pycaret.org

Alpaydın, E. (1999). Combined 5 × 2 cv F test for comparing supervised classification learning algorithms. *Neural Computation*, *11*(8), 1885–1892. https://doi.org/10.1162/089976699300016007

Bergstra, J. S., Bardenet, R., Bengio, Y., & Kégl, B. (2011). Algorithms for hyper-parameter optimization. *Advances in Neural Information Processing Systems*, *24*, 2546–2554.

Bergstra, J., Yamins, D., & Cox, D. (2013). Making a science of model search: Hyperparameter optimization in hundreds of dimensions for vision architectures. *Proceedings of the 30th International Conference on Machine Learning*, *28*(1), 115–123. https://proceedings.mlr.press/v28/bergstra13.html

Blackard, J. (1998). *Covertype* [Data set]. UCI Machine Learning Repository. https://doi.org/10.24432/C50K5N

Bouckaert, R. R., & Frank, E. (2004). Evaluating the replicability of significance tests for comparing learning algorithms. In H. Dai, R. Srikant, & C. Zhang (Eds.), *Advances in knowledge discovery and data mining* (Lecture Notes in Computer Science Vol. 3056, pp. 3–12). Springer. https://doi.org/10.1007/978-3-540-24775-3_3

Cai, B., Luo, Y., Guo, X., Pellegrini, F., Pang, M., de Moor, C., Shen, C., Charu, V., & Tian, L. (2025). Bootstrapping the cross-validation estimate. *The Annals of Applied Statistics*, *19*(4), 2981–3002. https://doi.org/10.1214/25-AOAS2036

Cohen, G., Afshar, S., Tapson, J., & van Schaik, A. (2017). EMNIST: Extending MNIST to handwritten letters. *2017 International Joint Conference on Neural Networks (IJCNN)*, 2921–2926. https://doi.org/10.1109/IJCNN.2017.7966217

Collobert, R., Bengio, S., & Bengio, Y. (2001). A parallel mixture of SVMs for very large scale problems. *Advances in Neural Information Processing Systems*, *14*, 633–640.

DeLong, E. R., DeLong, D. M., & Clarke-Pearson, D. L. (1988). Comparing the areas under two or more correlated receiver operating characteristic curves: A nonparametric approach. *Biometrics*, *44*(3), 837–845. https://doi.org/10.2307/2531595

Dhamala, E., Jamison, K. W., Jaywant, A., Dennis, S., & Kuceyeski, A. (2021). Distinct functional and structural connections predict crystallised and fluid cognition in healthy adults. *Human Brain Mapping*, *42*(10), 3102–3118. https://doi.org/10.1002/hbm.25420

DiCiccio, T. J., & Efron, B. (1996). Bootstrap confidence intervals. *Statistical Science*, *11*(3), 189–228. https://doi.org/10.1214/ss/1032280214

Dietterich, T. G. (1998). Approximate statistical tests for comparing supervised classification learning algorithms. *Neural Computation*, *10*(7), 1895–1923. https://doi.org/10.1162/089976698300017197

Efron, B., & Tibshirani, R. J. (1994). *An introduction to the bootstrap*. Chapman & Hall/CRC. https://doi.org/10.1201/9780429246593

Fischl, B. (2012). FreeSurfer. *NeuroImage*, *62*(2), 774–781. https://doi.org/10.1016/j.neuroimage.2012.01.021

Fisher, R. A. (1992). Statistical methods for research workers. In S. Kotz & N. L. Johnson (Eds.), *Breakthroughs in statistics: Methodology and distribution* (Vol. 2, pp. 66–70). Springer. https://doi.org/10.1007/978-1-4612-4380-9_6

Grinsztajn, L., Oyallon, E., & Varoquaux, G. (2022). Why do tree-based models still outperform deep learning on typical tabular data? *Advances in Neural Information Processing Systems, 35*, 507–520.

Harrell, F. E., Jr. (2001). *Regression modeling strategies: With applications to linear models, logistic regression, and survival analysis*. Springer. https://doi.org/10.1007/978-1-4757-3462-1

He, T., An, L., Chen, P., Chen, J., Feng, J., Bzdok, D., Holmes, A. J., Eickhoff, S. B., & Yeo, B. T. T. (2022). Meta-matching as a simple framework to translate phenotypic predictive models from big to small data. *Nature Neuroscience*, *25*(6), 795–804. https://doi.org/10.1038/s41593-022-01059-9

Hogg, R. V., McKean, J. W., & Craig, A. T. (2013). *Introduction to mathematical statistics* (7th ed.). Pearson.

Japkowicz, N., & Shah, M. (2011). *Evaluating learning algorithms: A classification perspective*. Cambridge University Press. https://doi.org/10.1017/CBO9780511921803

Komer, B., Bergstra, J., & Eliasmith, C. (2014). Hyperopt-Sklearn: Automatic hyperparameter configuration for Scikit-Learn. *Proceedings of the 13th Python in Science Conference*, 32–37. https://doi.org/10.25080/Majora-14bd3278-006

Kramer, C. Y. (1956). Extension of multiple range tests to group means with unequal numbers of replications. *Biometrics, 12*(3), 307–310. https://doi.org/10.2307/3001469

Lehmann, E. L., & Casella, G. (1998). *Theory of point estimation* (2nd ed.). Springer. https://doi.org/10.1007/b98854

Massey, F. J., Jr. (1951). The Kolmogorov-Smirnov test for goodness of fit. *Journal of the American Statistical Association*, *46*(253), 68–78. https://doi.org/10.1080/01621459.1951.10500769

Nadeau, C., & Bengio, Y. (2003). Inference for the generalization error. *Machine Learning, 52*(3), 239–281. https://doi.org/10.1023/A:1024068626366

Naeem, M., & Asghar, S. (2011). *KEGG Metabolic Relation Network (Directed)* [Data set]. UCI Machine Learning Repository. https://doi.org/10.24432/C5CK52

Newcombe, R. G. (1998). Two-sided confidence intervals for the single proportion: Comparison of seven methods. *Statistics in Medicine*, *17*(8), 857–872. https://doi.org/10.1002/(SICI)1097-0258(19980430)17:8%3C857::AID-SIM777%3E3.0.CO;2-E

Parkes, L., Moore, T. M., Calkins, M. E., Cieslak, M., Roalf, D. R., Wolf, D. H., Gur, R. C., Gur, R. E., Satterthwaite, T. D., & Bassett, D. S. (2021a). Network controllability in transmodal cortex predicts positive psychosis spectrum symptoms. *Biological Psychiatry, 90*(6), 409–418. https://doi.org/10.1016/j.biopsych.2021.03.016

Parkes, L., Moore, T. M., Calkins, M. E., Cook, P. A., Cieslak, M., Roalf, D. R., Wolf, D. H., Gur, R. C., Gur, R. E., Satterthwaite, T. D., & Bassett, D. S. (2021b). Transdiagnostic dimensions of psychopathology explain individuals' unique deviations from normative neurodevelopment in brain structure. *Translational Psychiatry*, *11*(1), 232. https://doi.org/10.1038/s41398-021-01342-6

Pedregosa, F., Varoquaux, G., Gramfort, A., Michel, V., Thirion, B., Grisel, O., Blondel, M., Prettenhofer, P., Weiss, R., Dubourg, V., Vanderplas, J., Passos, A., Cournapeau, D., Brucher, M., Perrot, M., & Duchesnay, É. (2011). Scikit-learn: Machine learning in Python. *Journal of Machine Learning Research*, *12*, 2825–2830.

Peneder, P., Stütz, A. M., Surdez, D., Krumbholz, M., Semper, S., Chicard, M., Sheffield, N. C., Pierron, G., Lapouble, E., Tötzl, M., Ergüner, B., Barreca, D., Rendeiro, A. F., Agaimy, A., Boztug, H., Engstler, G., Dworzak, M., Bernkopf, M., Taschner-Mandl, S., … Tomazou, E. M. (2021). Multimodal analysis of cell-free DNA whole-genome sequencing for pediatric cancers with low mutational burden. *Nature Communications*, *12*(1), 3230. https://doi.org/10.1038/s41467-021-23445-w

Quade, D. (1979). Using weighted rankings in the analysis of complete blocks with additive block effects. *Journal of the American Statistical Association*, *74*(367), 680–683. https://doi.org/10.1080/01621459.1979.10481670

Rao, C. R. (1948). Large sample tests of statistical hypotheses concerning several parameters with applications to problems of estimation. *Mathematical Proceedings of the Cambridge Philosophical Society*, *44*(1), 50–57. https://doi.org/10.1017/S0305004100023987

Raschka, S. (2018). *Model evaluation, model selection, and algorithm selection in machine learning* (arXiv:1811.12808). arXiv. https://doi.org/10.48550/arXiv.1811.12808

Shannon, P., Markiel, A., Ozier, O., Baliga, N. S., Wang, J. T., Ramage, D., Amin, N., Schwikowski, B., & Ideker, T. (2003). Cytoscape: A software environment for integrated models of biomolecular interaction networks. *Genome Research*, *13*(11), 2498–2504. https://doi.org/10.1101/gr.1239303

Siegel, S. (1956). *Nonparametric statistics for the behavioral sciences*. McGraw-Hill.

Wilcoxon, F. (1945). Individual comparisons by ranking methods. *Biometrics Bulletin*, *1*(6), 80–83. https://doi.org/10.2307/3001968

Wright, S. (1921). Correlation and causation. *Journal of Agricultural Research*, *20,* 557–585.

Wulan, N., An, L., Zhang, C., Kong, R., Chen, P., Bzdok, D., Eickhoff, S. B., Holmes, A. J., & Yeo, B. T. T. (2024). Translating phenotypic prediction models from big to small anatomical MRI data using meta-matching. *Imaging Neuroscience*, *2*, 1–21. https://doi.org/10.1162/imag_a_00251

Yan, X., & Su, X. G. (2009). *Linear regression analysis: Theory and computing*. World Scientific. https://doi.org/10.1142/6986
